# A tale of two fusion proteins: understanding the metastability of human respiratory syncytial virus and metapneumovirus and implications for rational design of uncleaved prefusion-closed trimers

**DOI:** 10.1101/2024.03.07.583986

**Authors:** Yi-Zong Lee, Jerome Han, Yi-Nan Zhang, Garrett Ward, Keegan Braz Gomes, Sarah Auclair, Robyn L. Stanfield, Linling He, Ian A. Wilson, Jiang Zhu

## Abstract

Respiratory syncytial virus (RSV) and human metapneumovirus (hMPV) cause human respiratory diseases and are major targets for vaccine development. In this study, we designed uncleaved prefusion-closed (UFC) trimers for the fusion (F) proteins of both viruses by examining mutations critical to F metastability. For RSV, we assessed four previous prefusion F designs, including the first and second generations of DS-Cav1, SC-TM, and 847A. We then identified key mutations that can maintain prefusion F in a native-like, closed trimeric form (up to 76%) without introducing any interprotomer disulfide bond. For hMPV, we developed a stable UFC trimer with a truncated F_2_-F_1_ linkage and an interprotomer disulfide bond. Tens of UFC constructs were characterized by negative-stain electron microscopy (nsEM), x-ray crystallography (11 RSV-F and one hMPV-F structures), and antigenic profiling. Using an optimized RSV-F UFC trimer as bait, we identified three potent RSV neutralizing antibodies (NAbs) from a phage-displayed human antibody library, with a public NAb lineage targeting sites Ø and V and two cross-pneumovirus NAbs recognizing site III. In mouse immunization, rationally designed RSV-F and hMPV-F UFC trimers induced robust antibody responses with high neutralizing titers. Our study provides a foundation for future prefusion F-based RSV and hMPV vaccine development.

**ONE-SENTENCE SUMMARY:** The metastability analysis of fusion proteins has informed rational design of uncleaved prefusion-closed trimers for RSV and hMPV vaccine development.

## INTRODUCTION

Respiratory syncytial virus (RSV) and human metapneumovirus (hMPV) pose a significant threat to public health worldwide (*1, 2*). RSV is a major cause of lower respiratory tract (LRT) infections in infants, young children, the elderly, and immunocompromised individuals (*3–6*). The recently discovered hMPV affects the same population (*7–12*), causing upper and lower respiratory tract infections in mostly young children and individuals who suffer from asthma, pulmonary diseases, and cancer (*13–17*). Furthermore, hMPV is capable of coinfections with other respiratory viruses, including RSV (*11, 18, 19*), thus increasing mortality and morbidity in young children (*20*). Both RSV and hMPV are enveloped, non-segmented, negative-sense, single-stranded RNA viruses of the *Pneumoviridae* family (*1, 2*). Their viral genomes encode three surface proteins: glycoprotein (G) that is responsible for viral attachment to cell surface factors, fusion protein (F), which fuses viral and host cell membranes during viral entry (*21, 22*), and small hydrophobic (SH) protein, which is a viroporin. While both G and F can be recognized by host neutralizing antibodies (NAb) during natural infection (*23–26*), F is highly conserved among virus subtypes and represents a primary target for both RSV and hMPV vaccine development (*27, 28*).

RSV-F and hMPV-F are class I viral fusion glycoproteins with ∼30% sequence identity. The two F proteins share many structural and functional similarities in their pre- and post-fusion states, which have led to the design of a pan-pneumovirus F antigen (*29*). To enable cell entry, the RSV-F precursor, F_0_, is first cleaved by furin-like proteases at two sites to remove a 27-amino-acid (aa) peptide (p27) (*30, 31*) and generate two subunits: an N-terminal F_2_ that is attached to a larger C-terminal F_1_ subunit by two disulfide bonds to form a heterodimer, three of which assemble into a functional trimer (*27*). Some studies suggest that trimerization occurs after proteolytic activation (*32, 33*), which would make RSV-F distinct from other class I fusion proteins, such as HIV-1 envelope glycoprotein (Env), but it is unclear whether this prefusion F trimer adopts a fully closed conformation on virions. For hMPV, proteolytic cleavage at a single site by serine proteases (*34*) transforms F_0_ into F_2_ and F_1_ subunits, which form a functional prefusion hMPV-F trimer (*35*). The metastable prefusion RSV-F and hMPV-F then undergo irreversible refolding, during which the hydrophobic fusion peptides are ejected from the central cavity of the F trimer and insert into host cell membranes to facilitate virus-host membrane fusion and the rapid transition of F into a highly stable postfusion form (*27, 35*). An important lesson from decades of RSV vaccine research and clinical studies is that postfusion F elicits weak NAb or nonfunctional antibody responses with adverse effects, making it unsuitable for use as a vaccine antigen (*36*). In contrast, prefusion F induces superior NAb responses accounting for most of the RSV-neutralizing activity in human immune sera (*25, 26*). Passive transfer of prefusion F-specific NAbs, including maternal transfer of such antibodies, may effectively prevent or treat RSV infection (*37–39*).

Structure-based antigen design has played an essential role in the development of prefusion RSV-F vaccines (*40, 41*). Three prefusion F-specific NAbs (D25, AM22, and 5C4) were key to the determination of the first prefusion RSV-F structure (*42*), which led to a breakthrough in the first prefusion-stabilized F design, DS-Cav1(*43*). Based on the structural information, alternative designs were proposed to stabilize prefusion RSV-F in either cleaved or uncleaved forms (*33, 44, 45*). Structures of prefusion RSV-F in complex with diverse NAbs (*32, 46–52*) have defined six major antigenic sites, two of which (Ø and V) are prefusion-specific and can elicit NAbs that are 10-100 times more potent than those targeting sites (I, II, III, and IV) that are accessible in both pre- and postfusion states (*36, 53*). These studies paved the way for two approved RSV vaccines, ABRYSVO (GlaxoKlineSmith [GSK]) (*54*) and AREXVY (Pfizer) (*55*), as well as other vaccine candidates in various stages of preclinical and clinical development (*56–58*). Additionally, an mRNA vaccine showed 83.7% efficacy against RSV in older adults (*59*) that was comparable to an efficacy of 82.6-88.9% for the two marketed protein-based vaccines. For hMPV, similar design strategies (e.g., disulfide bond, proline, and cavity-filling substitutions) have recently been used to stabilize prefusion F for vaccine development (*35, 60–62*). In addition to 101F and MPE8, which are previously identified NAbs targeting both RSV and hMPV (*63, 64*), further hMPV NAbs were identified and structurally characterized using prefusion F (*65–67*). Although these prefusion hMPV-F constructs showed promising results in animal studies (*60, 61*), significant gaps remain between preclinical research and vaccine approval for human use. Nevertheless, structure-based rational antigen design will likely also be the driving force for hMPV vaccine development, as it has been for RSV.

Recently, we established a rational vaccine strategy for class-I viral fusion glycoproteins based on the analysis of their metastability (*68–71*). Such analyses were proposed to lead to general design principles applicable to diverse subtypes within the virus family to stabilize their surface antigens in a native-like, prefusion state. We also found that the causes of metastability could be encoded by different elements in the fusion domain. For HIV-1 Env, the N-terminal bend of heptad repeat 1 (HR1_N_) in gp41 was identified as the major cause of metastability (*70, 71*), whereas for Ebola virus (EBOV) glycoprotein (GP) and SARS-CoV-2 spike, the long stalk in GP2 and triple hinged HR2 in S2 were key contributors to metastability, respectively. Notably, metastability may have different forms: while wildtype and early-generation uncleaved HIV-1 Env constructs yielded non-native trimers (*72–74*), the unmutated EBOV GP with the mucin deletion (GPΔmuc) could not be retained as a closed trimer (*69*). For RSV-F and hMPV-F, the strong tendency to undergo a rapid pre-to-postfusion conformational rearrangement represents a distinct form of metastability. A recent study reported antibody-induced transient opening of the prefusion RSV-F trimer, suggesting another form of metastability (*50*). Although diverse designs have been reported to stabilize prefusion RSV-F (*33, 43–45*) and hMPV-F (*35, 60–62*), the causes of metastability have largely remained unclear for these two pneumovirus fusion proteins.

In this study, we designed prefusion RSV-F and hMPV-F based on metastability analysis. Negative-stain electron microscopy (nsEM) and x-ray crystallography were utilized to structurally characterize F proteins in addition to extensive biochemical, biophysical, and antigenic analyses. As controls, we first evaluated previously reported RSV-F designs: DS-Cav1 (*43*), SC-TM (*33*), sc9-10 DS-Cav1 (*44*), and 847A (*45*), with and without a His_6_ tag. We next designed and assessed, under the same experimental conditions as for the controls, uncleaved, prefusion-closed (UFC) RSV-F trimers derived from three “base” constructs, termed UFC_R1-3_ series. Introducing hydrogen bonds or hydrophobic interactions to an occluded acidic patch (^486^DEFD^489^) situated above the trimeric coiled-coil stalk in F_1_ substantially increased the ratio of prefusion-closed trimers from ∼4% to 76% in solution. A total of 11 crystal structures were determined to verify our RSV-F designs. We then applied a similar design strategy to hMPV-F. Indeed, a slightly shortened F_2_-F_1_ linkage and a well-positioned interprotomer disulfide bond could effectively stabilize hMPV-F in a prefusion-closed trimer, as confirmed by nsEM and a crystal structure at 6 Å resolution. To further evaluate these prefusion F constructs, we screened a human antibody library against an optimized RSV-F UFC trimer. Three potent NAbs were identified that shared the same epitopes as either the prefusion-specific NAb RSD5 targeting sites Ø and V (*51*), as confirmed by nsEM and a 4.0 Å-resolution crystal structure, or the cross-pneumovirus NAb MPE8 (*63*). The NAb A4 shared the same germline genes as RSD5 (*51*), thus defining a public antibody lineage. Lastly, we immunized mice with various RSV-F and hMPV-F constructs to evaluate their vaccine-induced antibody responses. In summary, the metastability analysis and UFC strategy presented in this study will facilitate next-generation vaccine development for RSV and hMPV.

## RESULTS

### Comparative analysis of previously reported prefusion RSV-F designs

Among four previously reported RSV-F designs (**Fig. S1a** and **Table S1a**), first-generation DS-Cav1 contains an engineered disulfide bond (S155C-S290C) in F_1_ and two hydrophobic mutations (S190F/V207L) (*43*). DS-Cav1 has been used to develop protein subunit, nanoparticle (NP), and mRNA vaccine candidates (*58, 59, 75, 76*) and has inspired other RSV-F designs (*33, 44, 45*). Unlike DS-Cav1, both SC-TM (*33*) and sc9-10 DS-Cav1 (*44*) are uncleaved and connect F_2_ and F_1_ with a GS linker, in addition to a proline mutation (S215P). Sc9-10 DS-Cav1 (*44*) also truncates the F_2_ C-terminus, removes the fusion peptide (FP), and includes an interprotomer disulfide bond (A149C-Y458C). 847A (*45*) represents another cleaved RSV-F design, containing a disulfide bond (T103C-I148C) between F_2_ and F_1_ and a D486S mutation in the acidic patch (^486^DEFD^489^). A foldon trimerization motif was appended to the F_1_ C-terminus (L513) with or without a His_6_ tag, producing a total of eight constructs for four RSV-F designs.

For our experiments, all eight RSV-F constructs were transiently expressed in ExpiCHO cells and purified either by an immobilized nickel affinity (Nickel) column or by immunoaffinity chromatography (IAC) using a site Ø-specific D25 antibody column (*42*). The purified protein was then characterized using size exclusion chromatography (SEC), differential scanning calorimetry (DSC), and nsEM (**Fig. S1b)**. For DS-Cav1 (*43*), the SEC profiles from both Nickel and D25 affinity columns contained a major aggregate peak at 8-9 ml, while the former resulted in a larger trimer peak relative to the latter (**Fig 1a**, left top). DS-Cav1-His_6_ produced two peaks in the DSC thermogram, with the first and second melting temperatures (T_m1_ and T_m2_) determined at 48.6°C and 71.7°C, respectively (**Fig 1a**, left bottom). The nsEM analysis revealed that the Nickel/SEC-purified trimer peak contained only a small fraction of closed trimers amidst a large population of dissociated prefusion trimers (**Fig. 1a**, right); similar results were seen for the D25/SEC-purified sample (**Fig. S1c**, left). For SC-TM (*33*), D25 purification resulted in a negligible yield, whereas a Nickel column produced a large trimer peak, as well as visible aggregate and monomer peaks (**Fig. 1b**, left top). Nickel/SEC-purified SC-TM-His_6_ was analyzed by DSC, which generated a similar double-peak thermograph with T_m1_ and T_m2_ determined at 50.7°C and 61.4°C, respectively (**Fig. 1b**, left bottom). In nsEM images, the Nickel/SEC-purified SC-TM-His_6_ sample contained both open and closed trimers, as well as monomers (**Fig. 1b**, middle left). The 2D classification analysis indicated that ∼28% of the trimers were in a prefusion-closed form, as further confirmed by fitting the crystal structure of SC-TM (PDB ID: 5C6B) into the nsEM density map (**Fig. 1b**, middle right). However, postfusion F could also be observed in EM micrographs (**Fig. 1b**, right). For the third design, sc9-10 DS-Cav1 (*44*), the two purification methods generated almost identical SEC profiles showing a single trimer peak with substantial yield and purity (**Fig. 1c**, left top). Greater thermostability was observed for the Nickel/SEC-purified sc9-10 DS-Cav1-His_6_ (**Fig. 1c**, left bottom), with T_m1_ and T_m2_ being ∼10-20°C higher than those of His-tagged DS-Cav1 and SC-TM. The nsEM analysis and structure fitting (PDB ID: 5K6I) indicated that sc9-10 DS-Cav1 contained ∼100% prefusion-closed trimers regardless of the purification method used (**Fig. 1c**, middle; **Fig. S1c**, right), likely attributed to the interprotomer disulfide bond. The last RSV-F design, 847A (*45*), showed little to no trimer yield after D25 purification but produced notable trimer and monomer peaks after Nickel purification (**Fig. 1d**, left top). 847A-His_6_ exhibited a unique DSC profile with a single T_m_ determined at 64.2°C (**Fig. 1d**, left bottom), which is close to the T_m1_ of sc9-10 DS-Cav1-His_6_. The nsEM analysis indicated that the closed trimer population was composed of both prefusion (∼94%) and postfusion (∼2%) F trimers (**Fig. 1d**, middle and right; **Fig. S1d**). The prefusion-closed form was confirmed by fitting the crystal structure of 847A (PDB ID: 7UJ3) into the nsEM density map (**Fig. 1d**, middle right). For the four His-tagged RSV-F constructs, the Nickel/SEC-purified trimer fractions were cross-linked using disuccinimidyl glutarate (DSG) and subjected to sodium dodecyl sulfate-polyacrylamide gel electrophoresis (SDS-PAGE) under reducing conditions (**Fig. S1e**). Regardless of the purification method used, His-tagged DS-Cav1, SC-TM, and 847A showed both trimer and monomer bands on the gel, as open trimers could not be cross-linked and would be reduced to monomers under denaturing conditions. In contrast, a concentrated trimer band was observed for sc9-10 DS-Cav1-His_6_.

**Fig. 1.**
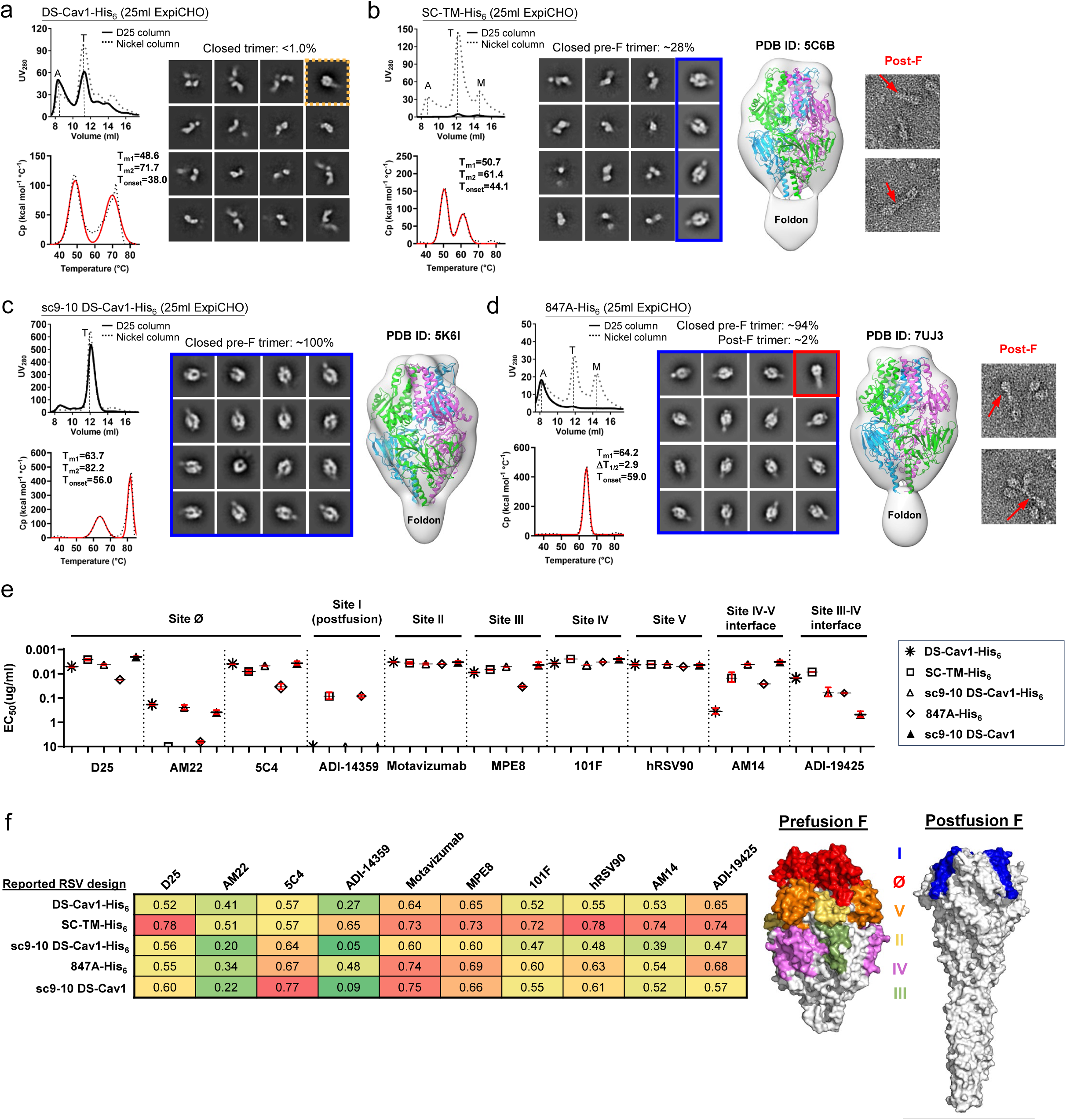
Comparative in vitro characterization of previously reported RSV-F designs. Size exclusion chromatography (SEC) profile (left top), differential scanning calorimetry (DSC) profile (left bottom), representative 2D classification images (middle, or right when no 3D models shown), and 3D reconstruction from the nsEM analysis (right) for (a) DS-Cav1, (b) SC-TM, (c) Sc9-10 DS-Cav1, and (d) 847A. All constructs contain a foldon trimerization motif with or without a C-terminal His_6_ tag, as indicated in the construct name with a suffix (-His_6_). A total of eight constructs were transiently expressed in 25 ml of ExpiCHO cells and purified using a Nickel column or a D25 antibody column. Major SEC peaks, such as aggregation (A), trimer (T), and monomer (A), are marked on the profile. The SEC profiles after Nickel and D25 antibody purification are shown as dotted and solid lines, respectively. For the 2D classification analysis, prefusion-closed trimers are circled in a blue line box, and closed trimers in the postfusion and non-prefusion states are circled in a red line box and a dotted orange line box, respectively. For SC-TM-His_6_ and 847A-His_6_, two micrographs are enlarged to show shapes characteristic of the postfusion F trimer. Prefusion RSV-F trimer structures (PDB ID: 5C6B, 5K6I, and 7UJ3) are used for structural fitting into the nsEM densities. (e) ELISA-derived EC_50_ (µg/ml) values of five RSV-F trimers binding to 10 antibodies that target six antigenic sites and two interface epitopes, which are labeled on the plots. (f) Biolayer interferometry (BLI) antigenic profiles of five RSV-F trimers binding to the same antibodies. Sensorgrams were obtained from an Octet RED96 instrument using an antigen titration series of six concentrations (starting at 600 nM followed by two-fold dilutions) and are shown in **Fig. S1g**. Peak values at the highest concentration are shown in a matrix, in which cells are colored in green (weak binding) to red (strong binding) (left). In (e) and (f), the four His-tagged RSV-F trimers were purified using a Nickel column, whereas sc9-10 DS-Cav1 was purified using a D25 antibody column. (g) Footprints of six antigenic sites colored on the surface representation of prefusion (PDB ID: 4JHW) and postfusion (PDB ID: 3RRR) RSV-F trimers (right).

Antigenicity was assessed using a panel of 10 antibodies targeting all major antigenic sites (defined in **Fig. S1a**), including D25, AM22, and 5C4 (site Ø) (*42, 46, 51*), ADI-14359 (site I) (*48*), motavizumab (site II) (*32, 77*), MPE8 (site III) (*63*), 101F (site IV) (*78*), hRSV90 (site V) (*47*), AM14 (the IV-V interface) (*32*), and ADI-19425 (the III-IV interface) (*48*). The four Nickel/SEC-purified RSV-F-His_6_ samples, together with a D25/SEC-purified sc9-10 DS-Cav1 sample, were analyzed by enzyme-linked immunosorbent assay (ELISA) (**Fig. 1e** and **Fig. S1f**). His-tagged DS-Cav1, SC-TM, and sc9-10 DS-Cav1 bound to prefusion-specific, site Ø-directed NAbs with higher affinities than 847A-His_6_, as indicated by the half maximal effective concentration (EC_50_). As expected, D25 purification resulted in an up to 2.1-fold improvement in EC_50_ for sc9-10 DS-Cav1 binding to NAb D25. Meanwhile, His-tagged SC-TM and 847A completely lost recognition by NAb AM22. Notably, His-tagged SC-TM and 847A showed strong affinities for the postfusion-specific, site I-directed non-NAb, ADI-14359, whereas negligible binding was observed for both DS-Cav1 and sc9-10 DS-Cav1 designs regardless of the purification method used. For most NAbs targeting sites II - V, the four designs showed similar binding profiles, with a relatively low affinity noted for Nickel/SEC-purified 847A-His_6_ and DS-Cav1-His_6_. For NAb MPE8 (*63*), which cross-neutralizes RSV and hMPV by targeting site III, 847A-His_6_ showed the lowest affinity among the five constructs. NAbs AM14 and ADI-19425 (*32, 48*) were included to probe the two interface epitopes. For AM14 (the IV-V interface), sc9-10 DS-Cav1 showed the highest binding affinity, whereas DS-Cav1-His_6_ defined the lowest with a fold difference of 14-110 in EC_50_, consistent with the open/dissociated trimers observed in the nsEM analysis (**Fig. 1a**, right). For ADI-19425 (the III-IV interface), sc9-10 DS-Cav1 exhibited low binding affinity, likely due to the interprotomer disulfide bond altering the III-IV interface (*44*). All five RSV-F samples were then subjected to biolayer interferometry (BLI) using the same antibody panel. The peak binding signals and equilibrium dissociation constant (*K*_D_) were determined to assess their antigenic properties (**Fig. 1f** and **Fig. S1g**). The BLI signals of these RSV-F constructs binding to the postfusion-specific ADI-14359 were consistent with the ELISA data (**Fig. 1e**). For sc9-10 DS-Cav1, D25 purification resulted in consistently higher NAb binding signals than Nickel purification (**Fig. 1f**).

Our results revealed unique properties associated with four previous RSV-F designs, which were used as controls. Overall, sc9-10 DS-Cav1 appeared to be the best performer, with nearly 100% prefusion closed trimers and well-preserved NAb epitopes, whereas two of the other RSV-F constructs (SC-TM-His_6_ and 847A-His_6_) expressed a detectable level of postfusion F. Our results did not always match previous data for these designs, which might be caused by differences in construct design, expression, purification, or sample handling; however, all of our experiments were conducted using the same methods throughout the study. This comparative analysis informed our design of a new generation of prefusion RSV-F.

### Design of uncleaved prefusion-closed (UFC) RSV-F trimers with minimum mutations

Sc9-10 DS-Cav1 (*44*) produces prefusion-closed trimers with high yield and purity but at the cost of introducing a large set of mutations and an interprotomer disulfide bond that alters a major NAb epitope. Here, we hypothesized that an uncleaved prefusion-closed (UFC) trimer could be designed for RSV-F involving a smaller set of mutations. We first derived a base construct, termed UFC_R1_, which contains the same F_2_-F_1_ connecting region as in sc9-10 DS-Cav1 (*44*), an intra-F_1_ disulfide bond (S155C-S290C), and S215P and E92D mutations (**Fig. S2a** and **Table S1b**). A foldon motif was attached to the F_1_ C-terminus in UFC_R1_ and all its derivatives. We further hypothesized that a second proline mutation (V185P, or named P2) in the HR1_N_-equivalent β3-β4 hairpin (*71*) might destabilize the postfusion state, resulting in a UFC_R1_-P2 construct. Lastly, we hypothesized that the acidic patch (^486^DEFD^489^) atop the coiled-coil F_1_ stalk is a major cause of RSV-F metastability, and a hydrogen bond (D486N/E487Q, or NQ) mutation or a hydrophobic (D486L/E487L, or L2) mutation might effectively maintain prefusion RSV-F in a closed trimer conformation.

Four constructs, UFC_R1_, UFC_R1_-P2, UFC_R1_-P2-NQ, and UFC_R1_-P2-L2, were characterized. Based on the overall more favorable antigenic properties of sc9-10 DS-Cav1 after D25 purification (**Fig. 1e-f**), all constructs were purified by a D25 affinity column following ExpiCHO expression. UFC_R1_ produced a similar SEC profile to sc9-10 DS-Cav1 with high trimer yield and purity (**Fig. 2a**, left top), but was less thermostable with lower T_m1_ (57.1°C) and T_m2_ (72.4°C) values (**Fig. 2a**, left bottom). In the nsEM analysis, only ∼4% of UFC_R1_ represented prefusion-closed trimers in 2D classes, which were used for 3D reconstruction and modeling (**Fig. 2a**, right). UFC_R1_-P2 generated a similar SEC profile with slightly reduced thermostability (1.3°C and 5.7°C lower for T_m1_ and T_m2_, respectively) compared with UFC_R1_ (**Fig. 2b**, left). In addition, UFC_R1_-P2 contained a similar fraction (∼4%) of prefusion-closed trimers in the nsEM analysis, although image quality was insufficient for 3D modeling (**Fig. 2b**, right). UFC_R1_-P2-NQ and UFC_R1_-P2-L2 represented two designs to avoid the interprotomer disulfide bond (A149C-Y458C) by altering the acidic patch. In wildtype RSV-F, this acidic patch creates “repulsive” charge-charge interactions around the trimer axis, thus destabilizing the closed trimer. For UFC_R1_-P2-NQ, the NQ mutation had little effect on trimer expression, generating a similar SEC profile to UFC_R1_ and UFC_R1_-P2 (**Fig. 2c**, left top). In the DSC analysis, the melting points remained comparable, but T_onset_ increased to 50°C (**Fig. 2c**, left bottom), suggesting a delayed denaturing step during heating. Notably, the nsEM analysis revealed a significantly increased ratio (73%) of prefusion-closed trimers, which allowed reliable 3D reconstruction and density fitting using a prefusion RSV-F structure (PDB ID: 4JHW) (**Fig. 2c**, right) (*42*). The effects of hydrophobic mutations at the acidic patch were examined using the UFC_R1_-P2-L2 construct. Overall, UFC_R1_-P2-L2 demonstrated a similar expression profile to other UFC_R1_ variants (**Fig. 2d**, left top). In terms of thermostability, UFC_R1_-P2-L2 had a further increased T_onset_ (52.1°C) compared with UFC_R1_-P2-NQ (**Fig. 2d**, left bottom). In the nsEM analysis, UFC_R1_-P2-L2 produced a slightly higher ratio of prefusion-closed trimers than UFC_R1_-P2-NQ, ∼76% *vs.* ∼73%, respectively (**Fig. 2d**, right). Based on these results, we combined a prefusion-specific NAb D25 and a postfusion-specific non-NAb ADI-14359 with nsEM to probe UFC_R1_-P2-NQ and UFC_R1_-P2-L2. In both cases, we observed D25 Fab-bound trimers, in addition to unbound D25 Fabs, unbound prefusion F monomers, and D25 Fab-bound prefusion F monomers (**Fig. 2e**, left; **Fig. S2b**, left). Both RSV-F trimers remained prefusion in the presence of ADI-14359, confirmed by the 2D classification and 3D reconstruction (**Fig. 2e**, right; **Fig. S2b**, right). Lastly, reducing SDS-PAGE analysis of cross-linked samples demonstrated consistent trimer bands on the gel, suggesting high homogeneity of the UFC_R1_ series (**Fig. S2c**).

**Fig. 2.**
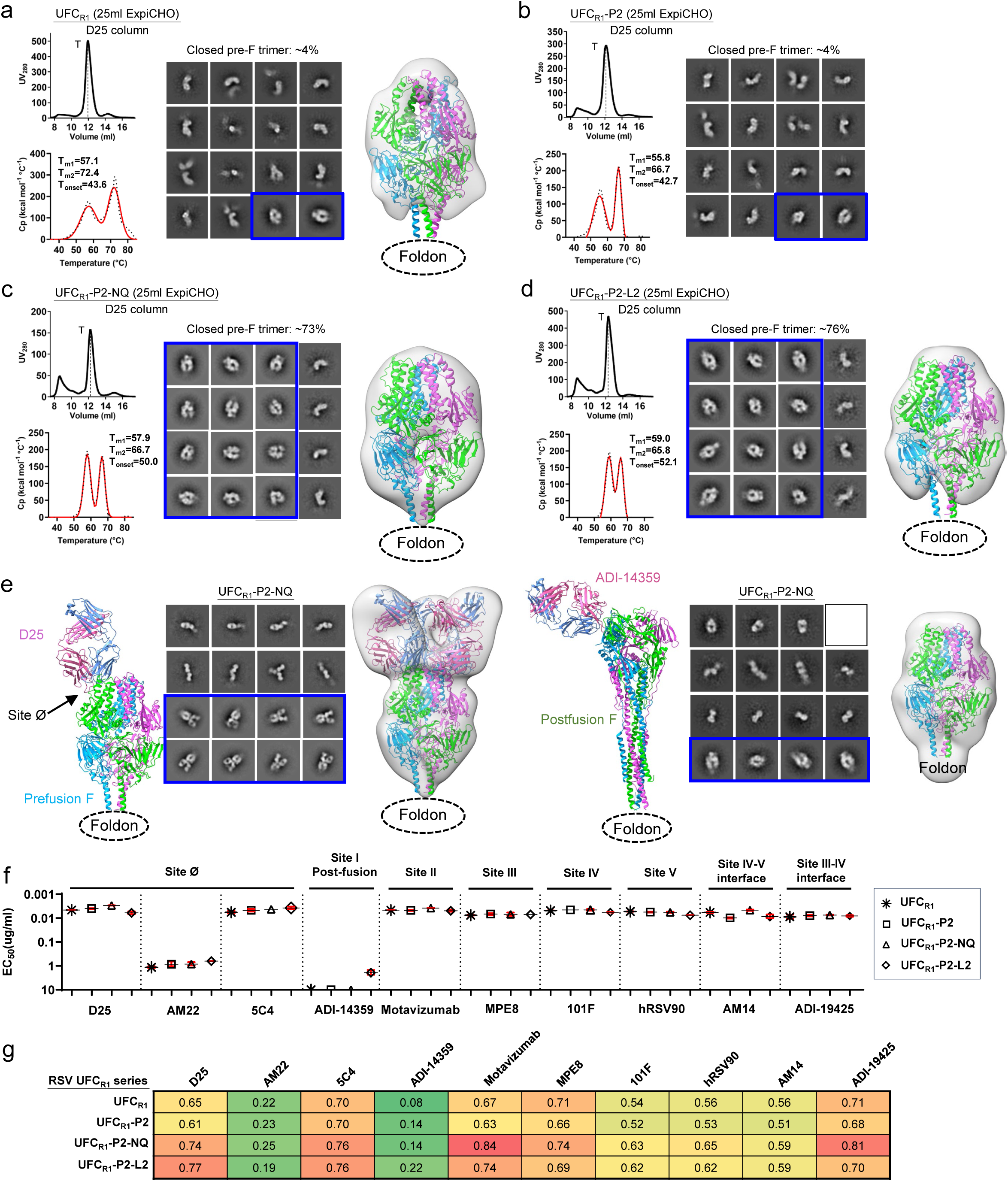
Design and in vitro characterization of RSV-F UFC_R1_ series. SEC profile (left top), DSC profile (left bottom), representative 2D classification images (middle, or right when no 3D models shown), and 3D reconstruction from nsEM analysis (right) for (a) UFC_R1_, (b) UFC_R1_-P2, (c) UFC_R1_-P2-NQ, and (d) UFC_R1_-P2-L2. All UFC_R1_ constructs were transiently expressed in 25 ml of ExpiCHO cells and purified using a D25 antibody column. The trimer (T) peak is marked on the profile. (e) The nsEM analysis of UFC_R1_-P2-NQ in the presence of prefusion-specific antibody D25 (left panel) or postfusion-specific antibody ADI-14359 (right panel). Each panel shows the ribbon model of the RSV-F/antibody complex (left), representative 2D classification images (middle), and 3D reconstruction (right). The 2D classes corresponding to prefusion-closed trimers (either ligand-free or antibody-bound) are circled in blue, and a prefusion RSV-F trimer (PDB ID: 4JHW) was used for structural fitting into the nsEM densities. (f) ELISA-derived EC_50_ (µg/ml) values of four UFC_R1_ constructs binding to 10 antibodies, as in Fig. 1e. (g) BLI-derived antigenic profiles of four UFC_R1_ constructs binding to 10 antibodies. Sensorgrams were obtained using the same protocol as in Fig. 1f and are shown in **Fig. S2e**. The matrix of peak values at the highest antigen concentration is shown, as in Fig. 1f.

Antigenicity of the four UFC_R1_ constructs was evaluated by ELISA and BLI using the 10-antibody panel. All four constructs showed high affinities for prefusion-specific, site Ø-targeting NAbs except AM22. In the ELISA, UFC_R1_-P2-NQ exhibited stronger binding to D25 than UFC_R1_-P2-L2 with EC_50_ values of 0.003 and 0.006 µg/ml, respectively (**Fig. 2f**, **Fig. S2d**). In BLI, UFC_R1_-P2-L2 showed a slightly higher D25-binding signal than UFC_R1_-P2-NQ (**Fig. 2g**, **Fig. S2e**). Notably, a detectable, albeit low, signal was observed for UFC_R1_-P2-L2 binding to the postfusion-specific site I-directed non-NAb, ADI-14359 (*48*), in both the ELISA and BLI. This unexpected result highlights the intricate balance between trimer stabilization and postfusion transition for mutations to the buried acidic patch. Because ADI-14359-bound UFC_R1_-P2-L2 trimers were not identified in any EM micrographs, a plausible explanation is that strong hydrophobic interactions introduced by the L2 mutation can cause conformational breathing that would transiently expose site I but could not escalate to an irreversible transition to the postfusion state. All four UFC_R1_ constructs showed high affinities for NAb ADI-19425 (*48*) (EC_50_ = 0.007-0.009 µg/ml), confirming that removal of the interprotomer disulfide bond can restore the III-IV interface epitope.

### Crystallographic characterization of UFC_R1_-series RSV-F trimers

Although low-resolution nsEM demonstrated that the NQ and L2 mutations substantially increased the ratio of prefusion-closed trimers, x-ray crystallography can provide atomic details of the altered interaction at the acidic patch. To achieve this goal, five crystal structures were obtained to validate RSV-F designs in a stepwise manner (**Tables S1b** and **S2**). The crystal structure of a ligand-free UFC_R1_ was determined at a resolution of 2.26 Å. The UFC_R1_ protomer showed root-mean-square deviations (RMSDs) of Cα atoms (Cα-RMSD) at 0.86 and 0.67 Å relative to DS-Cav1 (PDB ID: 4MMU) (*43*) and sc9-10 DS-Cav1 (PDB ID: 5K6I) (*44*), respectively, whereas a larger Cα-RMSD value of 1.13 Å was determined between DS-Cav1 and sc9-10 DS-Cav1 (**Fig. S3**). Notably, although the majority of UFC_R1_ adopted various open trimer forms in solution (**Fig. 2a**, middle), UFC_R1_ was seen in a fully closed trimer form in the crystal structure, suggesting that even a small percentage of closed trimers can crystallize (**Fig. 3a**). We also created a variant of UFC_R1_, with a 5GS flexible linker between RSV-F and a different trimerization motif (PDB ID: 1TD0), which was used in our previous studies (*68, 69, 79*). Indeed, a crystal structure at 2.28 Å resolution was obtained for UFC_R1_(1TD0), showing a Cα-RMSD of 0.50 Å relative to UFC_R1_ at the protomer level (**Fig. 3a**, right). Structural superimposition revealed largely similar backbone conformations (Cα-RMSD: 2.24 Å) for the loop connecting F_2_-S99 and F_1_-A149 (**Fig. 3a**, left), which would be buried within a prefusion-closed trimer. Next, we obtained a 2.70 Å-resolution crystal structure for UFC_R1_-P2-NQ, which largely resembled the UFC_R1_ structure with a Cα-RMSD of 0.53 Å at the protomer level (**Fig. 3b**, right). The V185P mutation disrupted a backbone hydrogen bond near the tip of the β3-β4 hairpin in the prefusion F (**Fig. 3b**, left) and would likely cause a “kink” in the extended α5 helix in the postfusion state (*80*). Similar proline mutations in HR1_N_ or HR1_N_-equivalent regions have been reported for prefusion-stabilized designs of HIV-1 Env (*70, 71, 81*) and EBOV GP (*69, 82*). The D486N/E487Q mutation replaced the repulsive charge-charge interaction with a hydrogen bond across the protomer interface (**Fig. 3b**, bottom), thus helping maintain the prefusion-closed trimer conformation. We also obtained a 2.30 Å-resolution crystal structure for UFC_R1_-L2 (**Fig. 3c**, left). UFC_R1_-L2 and UFC_R1_ shared high structural similarity with a protomer Cα-RMSD of 0.50 Å. In the UFC_R1_-L2 structure, two L486 residues of adjacent protomers form a hydrophobic contact across the interface, with Cγ-Cγ and Cδ-Cδ distances of 5.4 and 3.9 Å, respectively (**Fig. 3c**, right). By comparison, residues 486 and 487 do not interact in the crystal structures of SC-TM (*33*) or 847A (*45*), which contain E487Q and D486S mutations, respectively (**Fig. 3c**, bottom). Lastly, we examined whether adding the interprotomer disulfide bond (A149C-Y458C, or iSS) to UFC_R1_ would create a “simplified” version of sc9-10 DS-Cav1. A 2.30 Å-resolution crystal structure was obtained for this UFC_R1_-iSS construct (**Fig. 3d**, left). As expected, UFC_R1_-iSS closely resembled both UFC_R1_ and sc9-10 DS-Cav1 structures with protomer Cα-RMSDs of 0.54 and 0.61 Å, respectively. The interprotomer disulfide bond adopted a nearly identical geometry to that in sc9-10 DS-Cav1 (*44*) (**Fig. 3d**, right). Altogether, our crystallographic analyses provided detailed structural information supporting the UFC_R1_-series design principles.

**Fig. 3.**
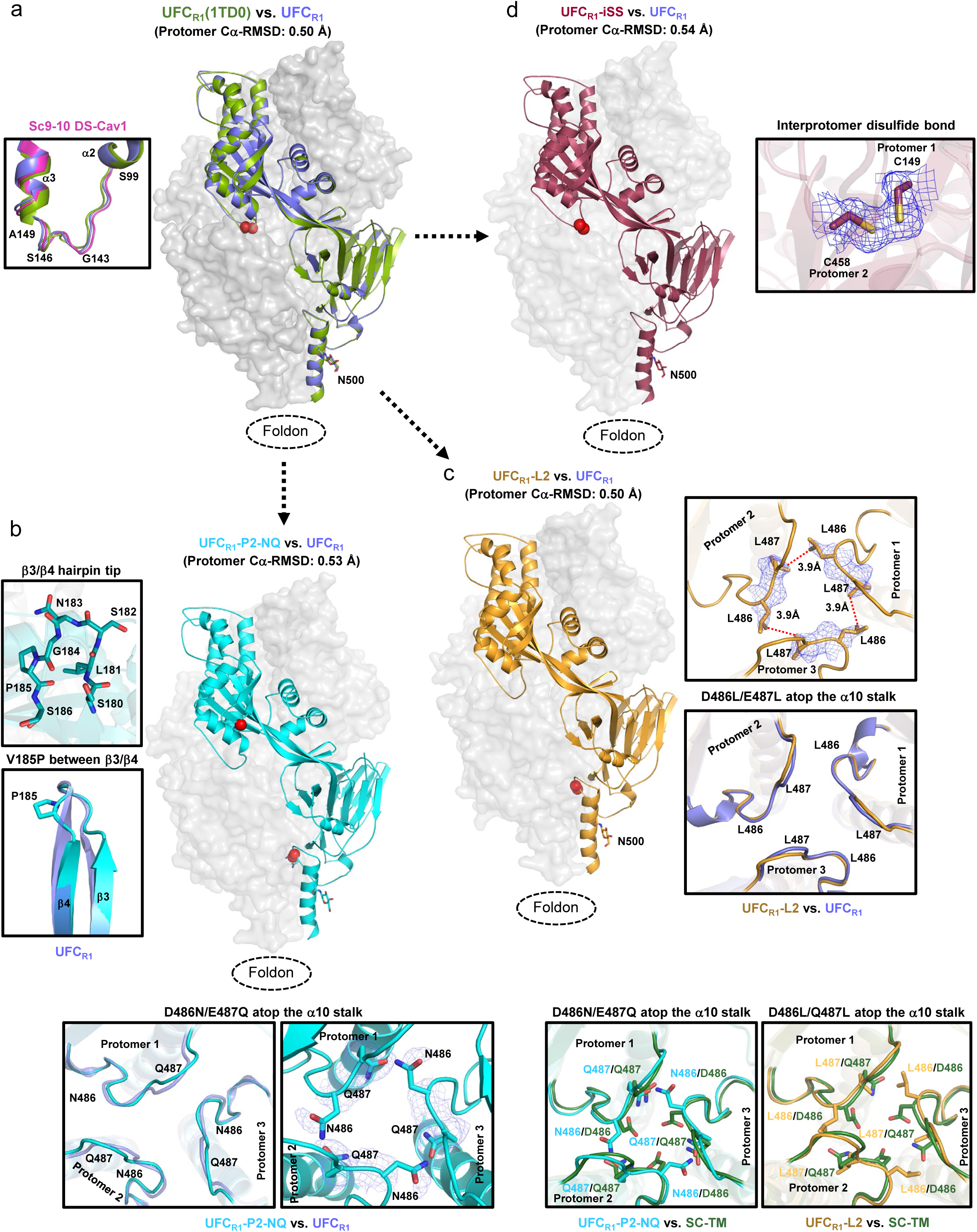
Crystallographic analysis of RSV-F UFC_R1_ series and variants. (a) Crystal structures of UFC_R1_ and UFC_R1_(1TD0) (2.26 and 2.28 Å) are superimposed and shown as green and blue ribbon models, respectively, within the gray trimer surface. The F_2_-F_1_ linkage is shown for UFC_R1_ and UFC_R1_(1TD0) with respect to sc9-10 DS-Cav1 (pink) in the left inset. (b) The crystal structure of UFC_R1_-P2-NQ (2.70 Å) is shown as a cyan ribbon model within the gray trimer surface. The atomic model of the β3/β4 hairpin tip and a close-up view of the V185P mutation between β3 and β4 are shown in the left insets, and the backbone and side chains of acidic patch mutations in UFC_R1_-P2-NQ (D486N-E487Q) are compared with UFC_R1_ in the bottom insets. (c) The crystal structure of UFC_R1_-L2 (2.30 Å) is shown as a gold ribbon model within the gray trimer surface. The backbone and side chains of acidic patch mutations in UFC_R1_-L2 (D486L-E487L) are compared with UFC_R1_ in the right insets, and structural details of this region in UFC_R1_-P2-NQ and UFC_R1_-L2 are compared with SC-TM in the bottom insets. (d) The crystal structure of UFC_R1_-iSS (2.3 Å) is shown as a red ribbon model within the gray trimer surface. A close-up view of the interprotomer disulfide bond (A149C-Y458C) within the density is shown in the right inset.

### Design and characterization of UFC_R2_-series RSV-F constructs

To investigate whether additional mutations can improve the ratio of prefusion-closed trimers, we created a second base construct, termed UFC_R2_, by adding two mutations (S46G and K465Q) to UFC_R1_. A total of four constructs, UFC_R2_, UFC_R2_-P2, UFC_R2_-P2-NQ, and UFC_R2_-P2-L2 (**Fig. S4a** and **Table S1c**), were evaluated using a similar strategy to UFC_R1_ (**Fig. 4**). For UFC_R2_, the SEC profile contained a shifted trimer peak (at ∼11.1 ml for UFC_R2_ vs. ∼11.9 ml for UFC_R1_) and a visible monomer peak (**Fig. 4a**, left top). Although UFC_R2_ appeared to be more thermostable than UFC_R1_, as indicated by higher T_m2_ (75.9°C *vs.* 72.4°C) and T_onset_ (51.4°C *vs.* 43.6°C) values (**Fig. 4a**, left bottom), no 2D classes representing prefusion-closed trimers were found by nsEM (**Fig. 4a**, right). In fact, D25-purified UFC_R2_ exhibited a tendency to dissociate into monomers. UFC_R2_-P2 behaved similarly to UFC_R2_ with less monomer content, as indicated by both SEC (**Fig. 4b**, left top) and nsEM (**Fig. 4b**, right), although its thermostability was slightly reduced, as indicated by DSC (T_m1_=56.2°C, T_m2_=69.9°C, and T_onset_=48.3°C) (**Fig. 4b**, left bottom). Incorporation of the NQ or L2 mutation generated a similar effect on the resulting UFC_R2_-P2-NQ and UFC_R2_-P2-L2 constructs, as it did on their UFC_R1_ counterparts (**Figs. 4c** and **4d**). Briefly, both mutations reduced the SEC peak corresponding to dissociated monomers, with UFC_R2_-P2-L2 showing the highest trimer purity (**Figs. 4c** and **4d**, left top). Similarly, these two mutations also improved RSV-F thermostability, with L2 slightly outperforming NQ in terms of T_m1_ (59.5°C vs. 58.5°C), which was ∼2-3°C higher than those of UFC_R2_ and UFC_R2_-P2 (**Figs. 4c** and **4d**, left bottom). However, nsEM revealed a significant difference in the ratio of prefusion-closed trimers between UFC_R2_-P2-NQ and -L2 following D25 and SEC purification, ∼6% and ∼28%, respectively (**Figs. 4c** and **4d**, middle). Nonetheless, fitting a prefusion RSV-F structure (PDB ID: 4JHW) (*43*) into the nsEM densities (**Figs. 4c** and **4d**, right) confirmed that both constructs produced prefusion-closed trimers. In reducing SDS-PAGE, cross-linked UFC_R2_ and UFC_R2_-P2 showed higher bands on the gel compared with UFC_R2_-P2-NQ and UFC_R2_-P2-L2 (**Fig. S4b**), indicative of an open form of the trimer structure. Therefore, two distant, seemingly unrelated mutations, S46G in β2 and K465Q in β22 (*44*), appeared to significantly reduce the ratio of prefusion-closed RSV-F trimers.

**Fig. 4.**
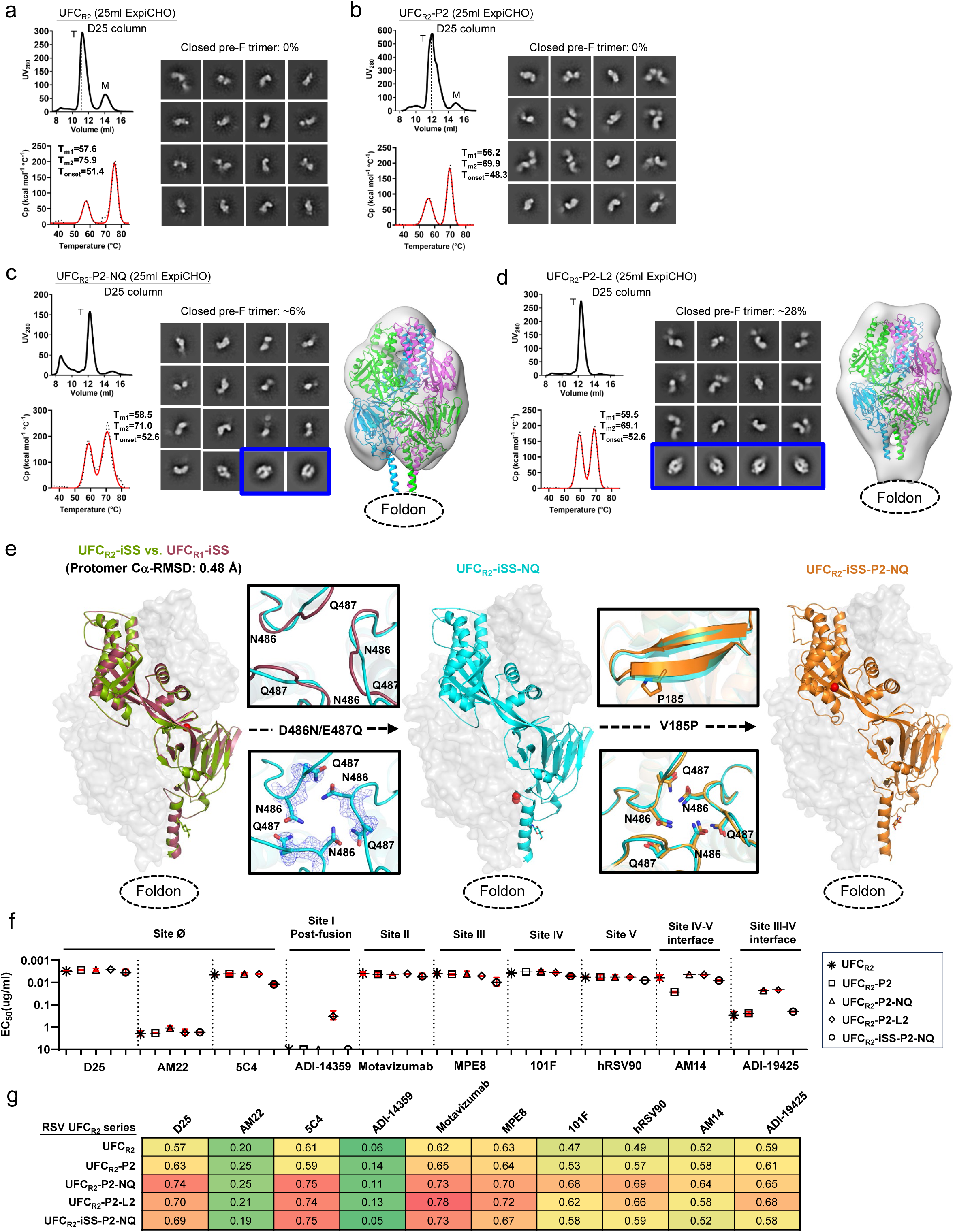
Design, in vitro characterization, and crystallographic analysis of RSV-F UFC_R2_ series. SEC profile (left top), DSC profile (left bottom), representative 2D classification images (middle, or right when no 3D models shown), and 3D reconstruction from nsEM analysis (right) for (a) UFC_R2_, (b) UFC_R2_-P2, (c) UFC_R2_-P2-NQ, and (d) UFC_R2_-P2-L2. All UFC_R2_ constructs were transiently expressed in 25 ml of ExpiCHO cells and purified using a D25 antibody column. The trimer (T) peak is marked on the profile. The 2D classification images corresponding to prefusion-closed trimers are circled in blue, and a prefusion RSV-F trimer (PDB ID: 4JHW) is used for structural fitting into the nsEM densities. (e) Crystallographic analysis of three UFC_R2_-derived constructs. Left: Crystal structures of UFC_R2_-iSS and UFC_R1_-iSS (2.83 and 2.30 Å) are superimposed and shown as green and red ribbon models, respectively, within the gray trimer surface. Middle: The crystal structure of UFC_R2_-iSS-NQ (2.30 Å) is shown as a cyan ribbon model within the gray trimer surface. The backbone and side chains of acidic patch mutations in UFC_R3_-iSS-NQ (D486N-E487Q) are compared with UFC_R2_-iSS in the insets to the left of the protomer/surface model. Right: The crystal structure of UFC_R2_-iSS-P2-NQ (2.30 Å) is shown as a gold ribbon model within the gray trimer surface. Details of the V185P mutation and acidic patch are shown in the insets to the left of the protomer/surface model. (f) ELISA-derived EC_50_ (µg/ml) values of five UFC_R2_ constructs binding to 10 antibodies, as in Fig. 1e. (g) BLI-derived antigenic profiles of five UFC_R2_ constructs binding to 10 antibodies. Sensorgrams were obtained using the same protocol as in Fig. 1f and are shown in **Fig. S4e**. The matrix of peak values at the highest antigen concentration is shown, as in Fig. 1f.

Although extensive screening of the UFC_R2_ constructs did not result in diffraction-quality crystals, the interprotomer disulfide bond (A149C-Y458C, iSS) stabilized the UFC_R2_ constructs and led to three crystal structures (**Tables S1c** and **S3**). UFC_R2_-iSS yielded Cα-RMSDs of 1.09 and 0.55 Å relative to DS-Cav1 and sc9-10 DS-Cav1, respectively (**Fig S4c**). It was not surprising that UFC_R2_-iSS was structurally more similar to sc9-10 DS-Cav1 because two additional mutations (S46G and K465Q) from sc9-10 DS-Cav1 were included in the UFC_R2_ construct. Structural superposition revealed nearly identical UFC_R1_-iSS and UFC_R2_-iSS backbones with a protomer Cα-RMSD of 0.48 Å (**Fig. 4e**, left), indicating that the S46G and K465Q mutations had little impact on the prefusion RSV-F structure. Compared with UFC_R2_-iSS, the NQ mutation resulted in the backbone of the acidic patch moving slightly outward toward each protomer, with a local Cα-RMSD of 1.40 Å, to make room for the hydrogen bond between N486 and Q487 across the protomer interface in UFC_R2_-iSS-NQ (**Fig. 4e**, middle and left inset). Lastly, a UFC_R2_-iSS-P2-NQ construct was created that showed nearly identical hydrogen bonding patterns to UFC_R2_-iSS-NQ and an unchanged β4 backbone despite the V185P (or P2) mutation (**Fig. 4e**, right and left inset). These results suggest that the S46G/K465Q mutation makes the UFC_R2_ backbone more flexible and potentially more amenable to global structural changes caused by mutations.

Antigenicity of five UFC_R2_-series constructs, including UFC_R2_, UFC_R2_-P2, UFC_R2_-P2-NQ, UFC_R2_-P2-L2, and UFC_R2_-iSS-P2-NQ, was evaluated by ELISA (**Fig. 4f, Fig. S4d**) and BLI (**Fig. 4g**, **Fig. S4e**) using the 10-antibody panel. Overall, the UFC_R2_ series generated similar antigenic profiles to their UFC_R1_ counterparts. In the ELISA, UFC_R2_-P2-L2 showed low levels of ADI-14359 binding (EC_50_ = 0.32 µg/ml) as did UFC_R1_-P2-L2, confirming the conformational breathing effect caused by the L2 mutation. In addition, UFC_R2_ and UFC_R2_-P2 showed lower affinities for ADI-19425 than UFC_R2_-P2-NQ and -L2 with a ∼12-fold difference in EC_50_, in line with the finding that UFC_R2_ and UFC_R2_-P2 did not produce any prefusion-closed trimers in nsEM and thus bound less favorably to NAbs targeting the III-IV interface. The iSS mutation (A149C-Y458C) adversely affected UFC_R2_-iSS-P2-NQ binding to ADI-19425 as this interprotomer disulfide bond might alter the structure of the III-IV interface required for ADI-19425 recognition.

### Alternative mutations to the HR1_N_-equivalent β3-β4 hairpin revealed by the UFC_R3_ series

In previous studies, the HR1_N_ bend, or HR1_N_-equivalent region, has been identified as the major contributor of metastability for HIV-1 Env (*70, 71*) and EBOV GP (*69, 82*). Notably, the HR1_N_-eqivalent region in prefusion RSV-F adopts a hairpin formed by β3 and β4, compared with a 21-aa unstructured loop in HIV-1 Env and an 8-aa turn (termed HR1_C_) between two HR1 helices in EBOV GP. In addition to the proline mutation (V185P), other mutations may be designed to target the β3-β4 hairpin. To this end, we created a third type of construct, termed UFC_R3_, which contains a longer linker (GS)_4_ between F_2_-T103 and F_1_-A147, an intra-F_1_ disulfide bond (S155C-S290C), the S215P mutation, and a second intra-F_1_ disulfide bond (A177C-T189C) specifically designed to lock β3 and β4 in the prefusion hairpin structure and prevent the pre-to-postfusion transition (**Fig. S5a** and **Table S1d**). We structurally characterized UFC_R3_ and its two variants using x-ray crystallography (**Table S4**). A crystal structure was obtained for UFC_R3_ at a resolution of 2.69 Å, which showed a Cα-RMSD of 0.74 Å with respect to sc9-10 DS-Cav1 at the protomer level (**Fig. S5b**, left). In the first variant, the S46G/E86D/K465Q triple mutation was added into UFC_R3_, resulting in a 2.31 Å-resolution structure for UFC_R3_-GDQ with a protomer Cα-RMSD of 0.76 Å relative to sc9-10 DS-Cav1 (**Fig. S5b**, middle). In the second variant, E86D was added to UFC_R3_ with viral capsid protein SHP (PDB ID: 1TD0) replacing the foldon as the trimerization motif, resulting in a 3.20 Å-resolution structure for UFC_R3_-D(1TD0) with a protomer Cα-RMSD of 0.89 Å relative to sc9-10 DS-Cav1 (**Fig. S5b,** right). In all three structures, the extended F_2_-F_1_ loop formed a 4-residue protrusion in the middle of the loop and was found buried within the cavity of the closed trimer (**Fig. S5b**, right, top inset), while β3 and β4 exhibited slightly twisted backbones that facilitated disulfide bond formation (**Fig. S5b**, right, bottom inset). Crystallographic analyses of these UFC_R3_ variants revealed that prefusion RSV-F can tolerate various mutations in the HR1_N_-eqivalent β3-β4 hairpin and F_2_-F_1_ linkage, in addition to different trimerization motifs.

### Design and in vitro characterization of prefusion hMPV-F constructs

Multiple designs have been proposed to stabilize prefusion hMPV-F (*35, 60–62*) (**Fig. S6a** and **Table S5a**). In the cleaved DS-CavEs2, Hsieh et al. introduced a disulfide bond (T365C-V463C) between β14 and α10 to stabilize this membrane-proximal region, which, however, could also destabilize the C-terminal stalk essential to F trimerization (*61*). As a result, the crystallographic analysis revealed that DS-CavEs2 is a prefusion monomer (*61*). Kwong and colleagues evaluated various disulfide bonds, proline mutations, and cleavage site linkers (*60, 62*), arriving at a construct containing a short F_2_-F_1_ linker and three disulfide bonds (*60*). In their construct design, the D454C-V458C mutation was initially introduced as an interprotomer disulfide bond, but cryo-EM revealed the formation of an intra-F_1_ disulfide bond that drastically altered the local structure around the trimer base (*60*). Here, we followed a minimalist approach, akin to what we used for RSV, to rationally design UFC trimers for hMPV-F. To this end, we developed a base construct, termed UFC_M1_, which places a G6 linker between the shortened F_2_ C-terminus (F_2_-E92) and fusion peptide (F_1_-103), in addition to A185P, E80D, and disulfide bond (T127C-N153C) mutations, as well as a C-terminal His_6_ tag (**Fig. S6b** and **Table S5b**). We then hypothesized that a second proline mutation (V155P, or P2) in the HR1_N_-equivalent region (*71*) can destabilize the postfusion state, yielding a UFC_M1_-P2 construct. We further hypothesized that a single interprotomer disulfide bond (A120C-Q426C, iSS) is sufficient to maintain a prefusion-closed trimer, leading to a UFC_M1_-P2-iSS construct. Lastly, we created a UFC_M1_-P2-F_2_C-VL construct, in which the shortened F_2_ C-terminus (residues 87-92) was modified to remove buried charges and the interprotomer disulfide bond mutation (iSS) was replaced with a hydrophobic contact (A120V/Q426L).

Four hMPV-F designs, UFC_M1_, UFC_M1_-P2, UFC_M1_-P2-iSS, and UFC_M1_-P2-F_2_C-VL, were validated using the same procedure established for RSV-F. While a Nickel column was used to capture all F species, an IAC column was generated using NAb MPE8 (*63*) to target the prefusion F in hMPV-F purification. For UFC_M1_, Nickel purification yielded a trimer peak at ∼11.9 ml in the SEC profile that was 5-fold higher than from an MPE8 column (**Fig. 5a**, left top), as measured by ultraviolet absorbance at 280 nm (UV_280_). DSC produced a thermogram with overlapping peaks, with T_m1_ and T_m2_ determined at 53.0⁰C and 59.6⁰C, respectively (**Fig. 5a**, left bottom). The Nickel/SEC-purified trimer fractions were analyzed by nsEM, in which no 2D classes of prefusion-closed trimers were identified (**Fig. 5a**, right). Meanwhile, no postfusion molecules were found in the EM micrographs. For UFC_M1_-P2, the P2 mutation between β3 and β4 notably increased the hMPV-F yield, as shown by the SEC profile following Nickel purification (**Fig. 5b**, left top). DSC generated comparable thermal parameters with ∼1 °C higher T_m1_ and T_onset_ (**Fig. 5b**, left bottom). All 2D classes obtained from nsEM corresponded to prefusion hMPV-F monomers with a similar shape to prefusion RSV-F monomers and contained no prefusion-closed trimers (**Fig. 5b**, right). UFC_M1_-P2-iSS showed a lower yield after Nickel and MPE8 purification, but with a higher ratio of prefusion-closed trimers within the total hMPV-F protein (**Fig. 5c**, left top). Furthermore, DSC demonstrated a single narrow peak with a single T_m_ of 72⁰C and a T_onset_ of 58.3⁰C, which were substantially higher (by ∼12-19 °C and ∼13-14 °C, respectively) than the melting points of UFC_M1_ and UFC_M1_-P2 (**Fig. 5c**, left bottom). Remarkably, almost all 2D classes in the nsEM analysis represented prefusion-closed hMPV-F trimers (**Fig. 5c**, middle), which was further confirmed by 3D reconstruction and structural fitting (PDB ID: 5WB0) (*35*) (**Fig. 5c**, right). The last construct, UFC_M1_-P2-F_2_C-VL, showed a low trimer yield after MPE8 purification, although a Nickel column produced a similar SEC profile to UFC_M1_ and UFC_M1_-P2 (**Fig. 5d**, left top). The DSC thermogram contained two peaks: while T_m1_ was comparable to those of UFC_M1_ and UFC_M1_-P2, T_m2_ increased to 83.6⁰C (**Fig. 5d**, left bottom). Interestingly, the nsEM analysis of Nickel/SEC-purified trimer fractions indicated the presence of prefusion-closed trimers, partially open trimers, and misfolded hMPV-F (**Fig. 5d**, middle). The 3D reconstruction revealed a tightened trimer apex and a widening around the base, suggesting an intermediate fusion state (**Fig. 5d**, right). In reducing SDS-PAGE, the cross-linked hMPV-F protein produced monomer, dimer, and trimer bands on the gel for all four constructs except UFC_M1_-P2-iSS, which displayed a single trimer band (**Fig. S6c**).

**Fig. 5.**
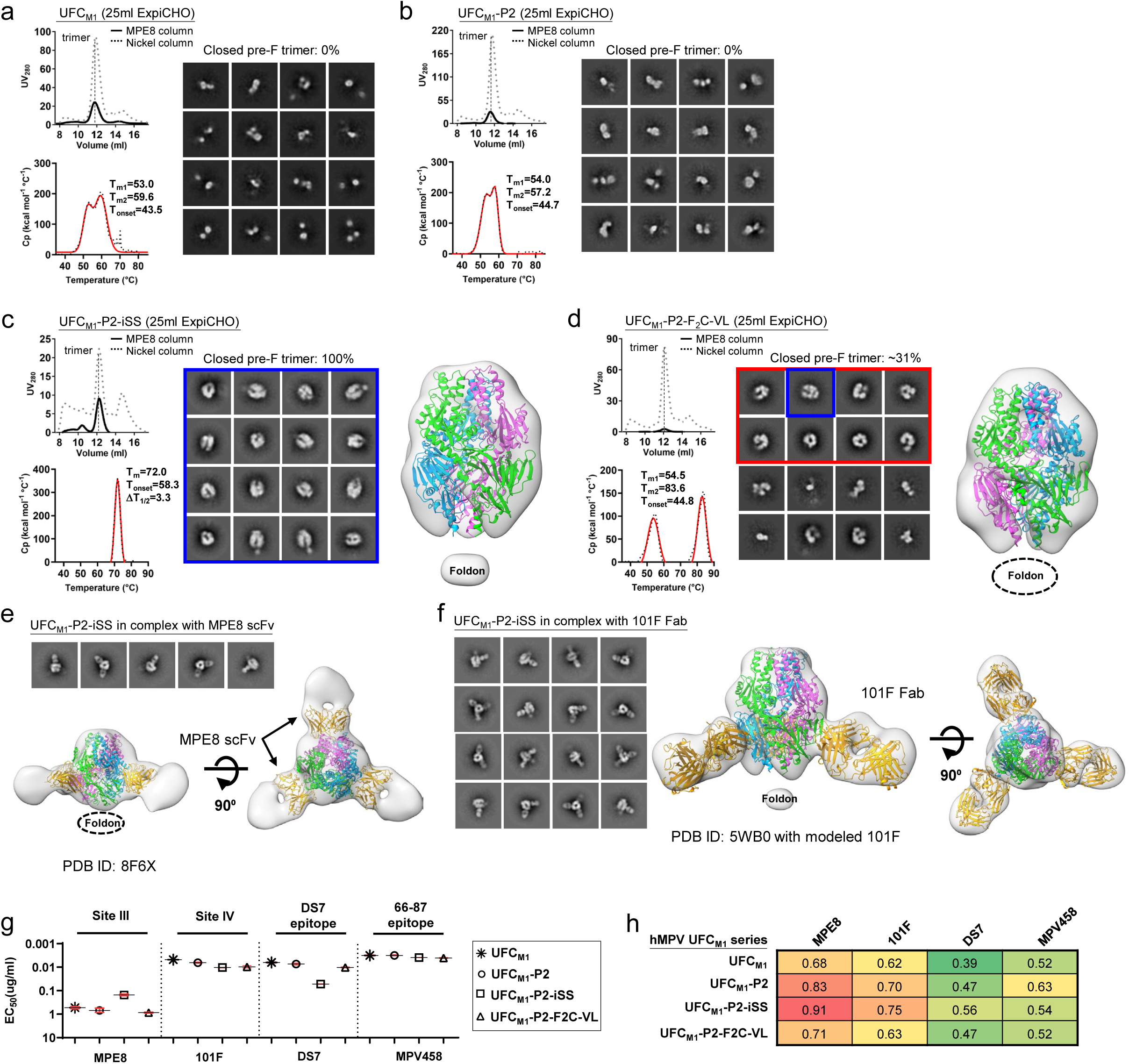
Design and in vitro characterization of hMPV-F UFC_M1_ series. SEC profile (left top), DSC profile (left bottom), representative 2D classification images (middle, or right when no 3D models shown), and 3D reconstruction from nsEM analysis (right) for (a) UFC_M1_, (b) UFC_M1_-P2, (c) UFC_M1_-P2-iSS, and (d) UFC_M1_-P2-F_2_C-VL. All UFC_M1_ constructs were transiently expressed in 25 ml of ExpiCHO cells and purified using an MPE8 antibody column and a Nickel column, as all constructs contain a His_6_ tag. The trimer (T) peak is marked on the profile. (e) The nsEM analysis of UFC_M1_-P2-iSS bound to antibody MPE8. Representative 2D classification images are shown on the top, and side and top views of 3D reconstruction of the complex are shown on the bottom left and right, respectively. A 3.25 Å-resolution cryo-EM model of MPE8 scFv-bound v3B Δ12_D454C-V458C (PDB ID: 8F6X) was used for density fitting. (f) The nsEM analysis of UFC_M1_-P2-iSS bound to antibody 101F. Representative 2D classification images are shown on the left, and side and top views of 3D reconstruction of the complex are shown on the right. A model of 101F Fab modeled onto a prefusion hMPV-F trimer (PDB ID: 5WB0) was used for density fitting. (g) ELISA-derived EC_50_ (µg/ml) values of four UFC_M1_ constructs binding to four antibodies, as in Fig. 1e. (h) BLI-derived antigenic profiles of four UFC_M1_ constructs binding to four antibodies. Sensorgrams were obtained using the same protocol as in Fig. 1f and are shown in **Fig. S6e**. The matrix of peak values at the highest antigen concentration is shown as in Fig. 1f.

To further characterize UFC_M1_-P2-iSS, we performed nsEM analyses of purified protein in complex with Fabs MPE8 (*63*) and 101F (*78*). The 3D reconstruction showed three MPE8 Fabs binding laterally to site III of a prefusion-closed trimer (**Fig. 5e**). The 3.25 Å-resolution cryo-EM model (EMDB-28891) of a recently reported hMPV-F design, v3B Δ12_D454C-V458C, bound to three single-chain variable fragments (scFv) of MPE8 (*60*) could be fitted into the EM density with an excellent match. The nsEM analysis indicated stronger 101F binding to UFC_M1_-P2-iSS, with more 2D classes showing two to three 101F Fabs binding to the hMPV-F trimer (**Fig. 5f**, left). Indeed, a 3D reconstruction with more structural detail was obtained for the 101F complex (**Fig. 5f**, right). The structural fitting of an hMPV-F/101F model revealed an upward angle of approach for 101F, which targets the exposed site IV. Together, our results indicate that UFC_M1_-P2-iSS can preserve important neutralizing epitopes on the prefusion-stabilized hMPV-F trimer.

Antigenicity of the four UFC_M1_ constructs was evaluated by ELISA and BLI using four NAbs with known complex structures, MPE8 (*63*), 101F (*78*), DS7 (*83*), and MPV458 (*66*). In the ELISA (**Fig. 5g** and **Fig. S6d**), UFC_M1_-P2-iSS bound preferably to MPE8 with a 3.5-5.7-fold higher EC_50_ than other UFC_M1_ constructs, consistent with the fact that MPE8 interacts with two protomers of a prefusion-closed trimer. In contrast, UFC_M1_-P2-iSS exhibited the lowest affinity for DS7, with a 5.0-8.3-fold difference in EC_50_ compared with other UFC_M1_ constructs (**Fig. 5g** and **Fig. S6d**). Further analysis of the DS7 complex structure (PDB ID: 4DAG) (*83*) revealed that its binding requires the displacement of β22 which only occurs in monomers or open trimers. The four hMPV-F constructs exhibited similar binding affinities for NAbs 101F and MPV458 (**Fig. 5f** and **Fig. S6d**). Overall, BLI demonstrated consistent patterns compared with ELISA, with UFC_M1_-P2-iSS showing the highest MPE8-binding signal (**Fig. 5g** and **Fig. S6e**).

### Crystallographic characterization of the hMPV-F UFC_M1_-P2-iSS trimer

We obtained a 6 Å-resolution structure for ExpiCHO-expressed, Nickel/SEC-purified UFC_M1_-P2-iSS using similar crystallization conditions to the first crystal structure of a prefusion hMPV-F design, 115-BV (*35*) (**Tables S5** and **S6**). The UFC_M1_-P2-iSS structure was superimposed onto the 115-BV structure (PDB ID: 5WB0) for comparison (**Fig. 6a**). In the asymmetric unit, UFC_M1_-P2-iSS adopted the same form as 115-BV, enabling the trimer structure to be modelled in a similar manner. UFC_M1_-P2-iSS yielded a Cα-RMSD of 1.22 Å with respect to 115BV at the protomer level. The three key elements of the UFC_M1_-P2-iSS design were compared to 115-BV, which has the most complete F structure bearing minimum mutations (**Fig. 6b**). The T127C-Q153C mutation was found to be critical to maintaining prefusion hMPV-F, with a C_β_-C_β_ distance of 4.2 Å in the DS-CavEs2 structure (*61*). From the fitted model, this disulfide bond had an estimated C_β_-C_β_ distance of 3.4 Å in UFC_M1_-P2-iSS, compared with a C_β_-C_β_ distance of 4.9 Å between T127 and Q153 in 115-BV (**Fig. 6b**, top). The V155P mutation appears to widen the β3-β4 turn in UFC_M1_-P2-iSS, which wouldlikely facilitate disulfide bond formation at positions 153 to 127 and destabilize postfusion F (**Fig. 6b**, middle). The interprotomer disulfide bond (C120-C426) had an estimated C_β_-C_β_ distance of 4.7 Å, thus locking hMPV-F in a prefusion-closed trimer conformation (**Fig. 6b**, bottom). Due to the limited resolution, the structure could not be resolved for the F_2_-F_1_ linkage, part of α8 (A344-S347), and part of α9-α10-β23 (V442-E457) (**Fig. S7**).

**Fig. 6.**
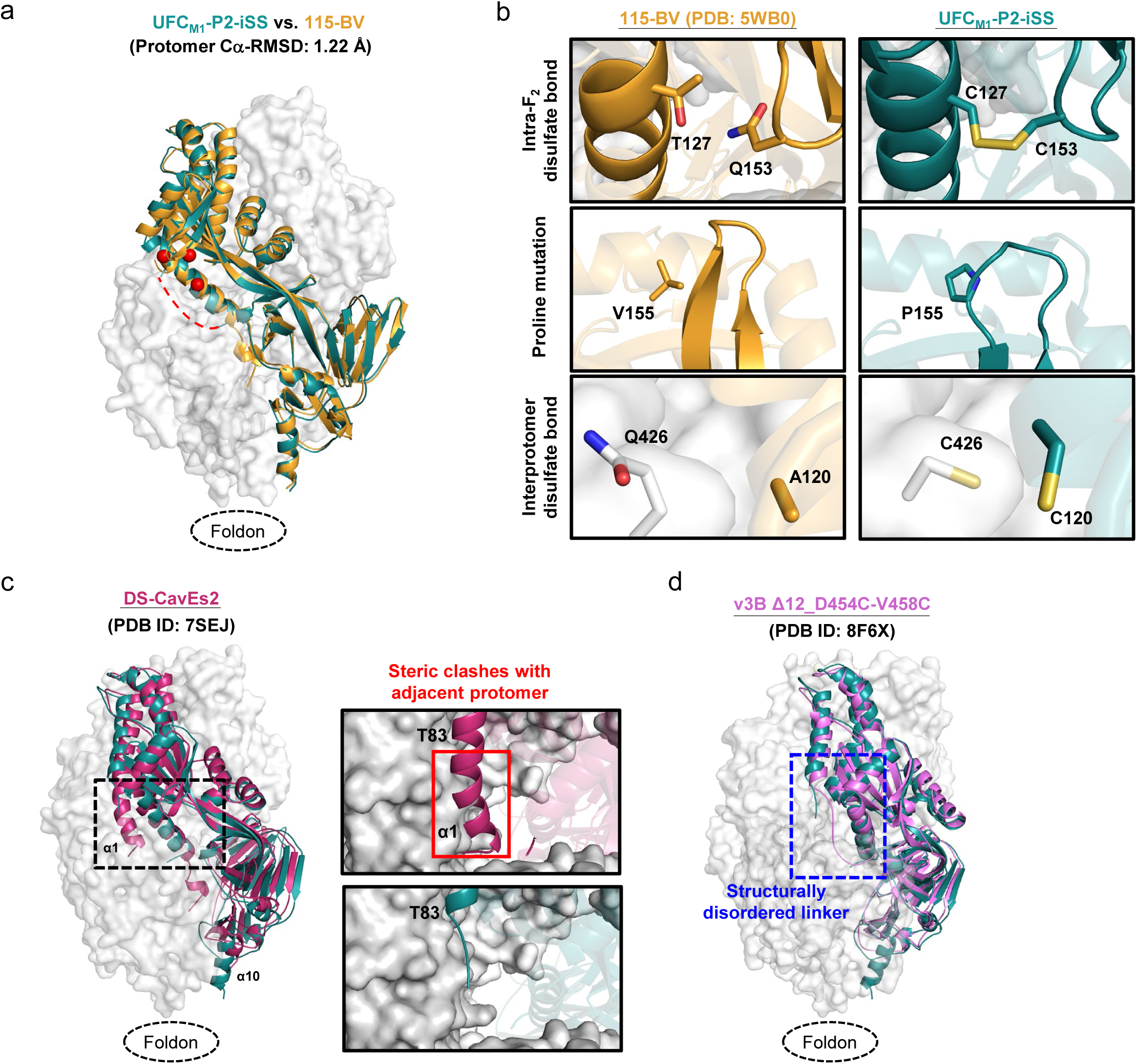
Crystallographic analysis of hMPV-F UFC_M1_-P2-iSS. (a) The crystal structure of UFC_M1_-P2-iSS (6.0 Å) is superimposed onto that of 115-BV (PDB ID: 5WB0), which are shown as green and gold ribbon models, respectively, within the gray trimer surface. Due to the limited resolution, structural details cannot be determined for the F_2_-F_1_ linker, A344-S347, and V442-E457. A red dotted line is added to show the expected approximate location of the missing F_2_-F_1_ linker. (b) Structural details of the intra-F_2_ disulfide bond T127C-Q153C, the V155P (P2) mutation inserted into the β3/β4 hairpin tip for destabilizing the postfusion state, and the interprotomer disulfide bond A120C-Q426C (iSS) are shown in the top, middle, and bottom insets, respectively. The crystal structure of 115-BV is included for comparison. (c) The crystal structures of UFC_M1_-P2-iSS and DS-CavEs2 (PDB ID: 7SEJ) are superimposed and shown as green and rouge pink ribbon models, respectively, within the gray trimer surface. The extended α1 helix in DS-CavEs2 that will clash with an adjacent protomer in a prefusion-closed trimer is circled in a black dotted line box. Close-up views of this region in DS-CavEs2 and UFC_M1_-P2-iSS are shown in the right insets. (d) Crystal structures of UFC_M1_-P2-iSS and v3B Δ12_D454C-V458C (PDB ID: 8F6X) are superimposed and shown as green and pink ribbon models, respectively, within the gray trimer surface. The F_2_-F_1_ linker region is circled in a blue dotted line box.

The UFC_M1_-P2-iSS structure was then compared to two leading prefusion hMPV-F designs: DS-CavEs2 (*61*) and v3B Δ12_D454C-V458C (*60*). The structural superposition of UFC_M1_-P2-iSS and DS-CavEs2 (PDB ID: 7SEJ) revealed differences in the stalk and α1 helix. Compared with a well-formed α10 helix in UFC_M1_-P2-iSS, DS-CavEs2 showed an incomplete α10 helix because of the intra-F_1_ disulfide bond (T365C-V463C) between β14 and α10, which destabilizes the C-terminal trimeric stalk (**Fig. 6c**, left). This may also explain why DS-CavEs2 crystalized as a monomer with an extended α1 helix that would clash with the adjacent protomer in a prefusion-closed trimer (**Fig. 6c**, right). Nonetheless, two cryo-EM structures showed trimeric DS-CavEs2 in complex with NAbs that interact with two protomers at the trimer interface, although the α10 helix was partially unstructured (*65, 67*). UFC_M1_-P2-iSS was then structurally superimposed onto v3B Δ12_D454C-V458C (PDB ID: 8F6X) (**Fig. 6d**). While both designs showed similar cleavage site linker structures, the trimer base (β23 and α10) in v3B Δ12_D454C-V458C adopted a non-native conformation due to the unintended intraprotomer disulfide bond (D454C-V458C) (*60*). In summary, our crystal structure, despite its modest resolution, validated the UFC_M1_-P2-iSS design and allowed for structural comparison with previously reported hMPV-F designs.

### Potent pneumovirus-neutralizing antibodies identified by RSV-F UFC_R1_-P2-NQ

As most children are infected by RSV before the age of two and will be reinfected throughout their adulthood (*84*), the healthy adult population provides a rich source of RSV NAbs. We hypothesize that if our lead RSV-F design, UFC_R1_-P2-NQ, can identify prefusion-specific RSV NAbs from a human antibody library, then it may induce similar NAbs in vaccination (**Fig. 7a**). Following our previously established protocol (*85*), we constructed a large phage-display scFv library using peripheral blood mononuclear cells (PBMCs) from 10 healthy donors and performed biopanning experiments to screen this human scFv library using UFC_R1_-P2-NQ as an antigen probe. After four biopanning steps, 96 clones were randomly selected for phage ELISA against sc9-10 DS-Cav1, a disulfide-locked prefusion-closed RSV-F trimer (*44*). A total of 36 scFv clones were sequenced (**Fig. S8a**), and eight with complete variable regions were selected as representative clones (**Fig. S8b**). Sequence analysis revealed that these eight clones are derived from five heavy chain variable (V_H_) genes and five λ/κ-light chain variable (V_L_/V_K_) genes (**Fig. 7b** and **Fig. S8c**). All eight clones in the scFv-Fc form could be expressed in HEK293 F cells with high yield except F1.

**Fig. 7.**
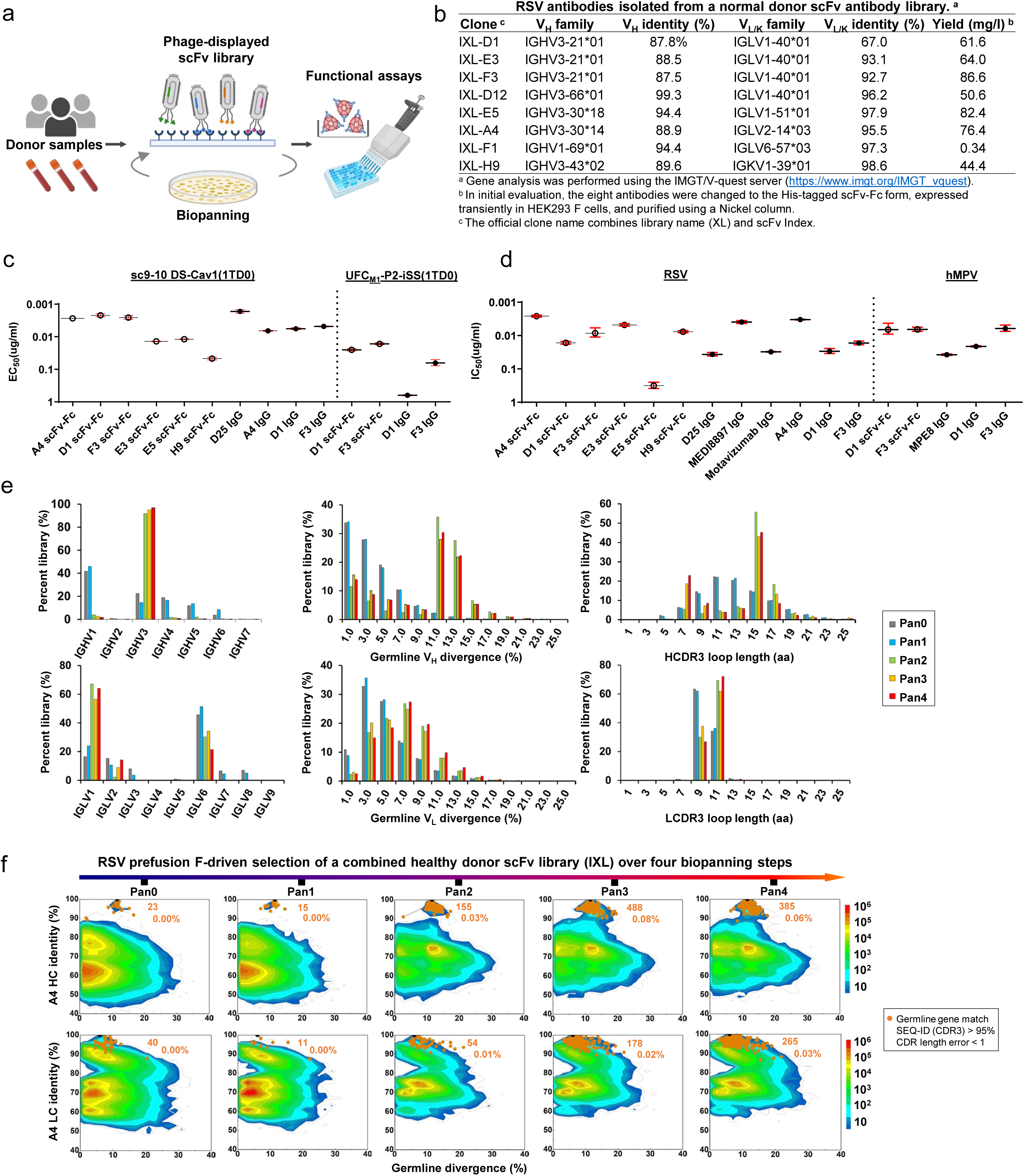
Potent pneumovirus neutralizing antibodies identified from a human antibody library. (a) Schematic representation of the phage display workflow. (b) Gene family analysis of eight scFv clones identified from a phage-displayed human antibody (scFv) library using RSV-F UFC_R1_-P2-NQ as a biopanning antigen. (c) ELISA-derived EC_50_ (µg/ml) values of library-derived antibodies binding to RSV-F sc9-10 DS-Cav1 and hMPV-F UFC_M1_-P2-iSS. (d) IC_50_ (µg/ml) values derived from live RSV and hMPV neutralization assays. (e) Distribution of germline gene usage, somatic hypermutation, and CDR3 length plotted for heavy chains (HCs) and λ-light chains (λ-LCs) of the scFv library during the biopanning process. (f) Identity-divergence analysis of the IXL-A4 (or A4) within the scFv library during the biopanning process. The sequence datasets used in (e) and (f) were obtained from next-generation sequencing (NGS) of the scFv libraries on an Ion GeneStudio S5 platform. For the heatmaps in (f), after data processing using an Antibodyomics pipeline, each sequence is plotted as a function of sequence identity from a reference antibody chain and sequence divergence from the assigned germline gene. Color indicates sequence density at a particular point on the 2D plot. Sequences with the same germline gene, a CDR3 identity ≥ 95%, and a 1-residue error margin of CDR length calculation with respect to the reference antibody chain are plotted as orange dots with the number of sequences and gene family percentage labeled. The schematic representation of phage-based antibody isolation was created with BioRender.com.

These eight clones were evaluated in an ELISA against RSV-F using sc9-10 DS-Cav1 (*44*) and hMPV-F using UFC_M1_-P2-iSS (**Fig. 7c** and **Fig. S8d**). In the scFv-Fc form, A4, D1, and F3 showed higher affinities for sc9-10 DS-Cav1 than other RSV-F reactive clones, such as E3, E5, and H9, with a ∼4.8-18.9-fold difference in EC_50_. The binding affinities of A4, D1, and F3 scFv-Fc antibodies were largely comparable to D25 IgG (*42*). In the IgG form, these three clones displayed similar affinities for sc9-10 DS-Cav1, but slightly lower than D25, with a 2.9-3.9-fold difference in EC_50_. When tested against UFC_M1_-P2-iSS, only D1 and F3 exhibited any measurable binding to this prefusion hMPV-F trimer. Specifically, D1 and F3 had similar affinities for hMPV-F in the scFv-Fc form, which were reduced 25- and 3.8-fold, respectively, when changed to the IgG form. These phage library-derived antibodies demonstrated different potencies in live RSV and hMPV neutralization assays (**Fig. 7d** and **Fig. S8e**). In the scFv-Fc form, A4 appeared to be the best RSV neutralizer with a half maximal inhibitory concentration (IC_50_) of 0.0021 µg/ml, 1.8-fold higher than a highly optimized therapeutic antibody, MEDI8897 (Nirsevimab) (*38, 86*). In the IgG form, A4 showed a nearly identical IC_50_ to MEDI8897, F3 yielded a comparable IC_50_ to D25 (0.013 µg/ml *vs.* 0.011 µg/ml, respectively), and D1 exhibited similar potency to a site II-directed NAb, Motavizumab (*87*). Both D1 and F3 IgGs neutralized live hMPV, with F3 showing ∼11-fold higher potency, estimated by the IC_50_ value, than a widely studied cross-NAb, MPE8 (*63*).

To understand how these functional antibodies were selected during the biopanning process, we pooled the pre-panning and four post-panning scFv libraries for next-generation sequencing (NGS) on the Ion GeneStudio S5 platform using an Ion 530 chip. NGS yielded over ∼18.8 million raw reads, which were processed using the Antibodyomics 2.0 pipeline (*85*) to generate full-length V_H_ and V_L/K_ reads for bioinformatics analyses (**Fig. S8f**). A distinct pattern of antibody enrichment and a rapid convergence after two panning rounds were observed in the quantitative library profiles (**Fig. 7e** and **Fig. S8g**). In terms of germline gene usage, IGHV3 and IGLV1 accounted for 97% and 64% of the converged library, respectively, consistent with the finding that six of eight selected scFv clones were derived from the combination of these two genes (**Fig. 7b**). In terms of somatic hypermutation (SHM), both V_H_ and V_L/K_ distributions shifted from a germline-like (SHM: 1-3%) pre-panning population toward a more mature population after convergence, peaking at 11-13% and 7-9%, respectively. In terms of heavy chain complementarity-determining region 3 (HCDR3) length, the converged library contained two prevalent scFv families with 7-9 aa (22.9%) and 15-17 aa (45.2%) HCDR3 loops, compared with a normal distribution with an average of 13 aa in the pre-panning library. The κ-light chains showed little change in germline gene usage, SHM, and κ- chain CDR3 (LCDR3) length (**Fig. S8g**), suggesting that κ-light chains were not used by RSV-specific scFv clones. A 2D identity-divergence analysis was then conducted to visualize the scFv-derived heavy- and light-chain (HC and LC, respectively) populations during biopanning (**Fig. 7f** and **Fig. S8h**). The 2D plots revealed the rapid enrichment of D1, E3, and D12 clones and modest expansion of A4, F3, and E5 clones after two panning steps, but little selection pressure was noted for F1 and H9. Between the two most potent neutralizers, F3 exhibited more pronounced expansion than A4, which accounted for 1.0% and 0.03% of their germline gene families, respectively.

### Structural characterization of RSV and hMPV-neutralizing human antibodies

Of the eight scFv clones, A4, D1, and F3 displayed distinct binding and neutralization profiles. To investigate how these human antibodies recognize prefusion RSV-F, we generated A4, D1, and F3 Fabs and formed complexes with sc9-10 DS-Cav1 (*44*) for structural analysis. We first used nsEM to identify their epitopes on the prefusion trimer (**Figs. 8a** and **8b**). For A4, the 2D classification revealed sc9-10 DS-Cav1 trimers with Fabs bound to an epitope near the trimer apex (**Fig. 8a**, left). Compared with the 2D classes obtained for the D25-bound UFC_R1_-P2-NQ trimer (**Fig. 2e**, left), A4 shifted sideward and created a larger angle relative to the trimer axis in the side views (**Fig. 8a**, left). The visual inspection of previously reported RSV-F NAbs revealed that a germline version of NAb RSD5 (RSD5-GL) in complex with DS-Cav1 (PDB ID: 6DC3 (*51*)) could be fitted into the A4/sc9-10 DS-Cav1 density with a nearly perfect match (**Fig. 8a**, right). The crystal structure (PDB ID: 6DC3) revealed a distinctive RSD5 epitope (*51*) that mainly overlaps with site Ø, as defined by D25 (*42*), but also extends to site V, as defined by hRSV90 (*47*), 01.4B and ADI-14442 (*52*). For D1 and F3, nsEM revealed an angle of approach resembling that of the site III-specific NAb, MPE8 (*63*) (**Fig. 8b**, left), which was confirmed by fitting the MPE8/DS-Cav1 complex (*63*) into the nsEM densities of D1- and F3-bound sc9-10 DS-Cav1 (**Fig. 8b**, right). The MPE8-like epitope specificity also explained their reactivity with both RSV-F and hMPV-F.

**Fig. 8.**
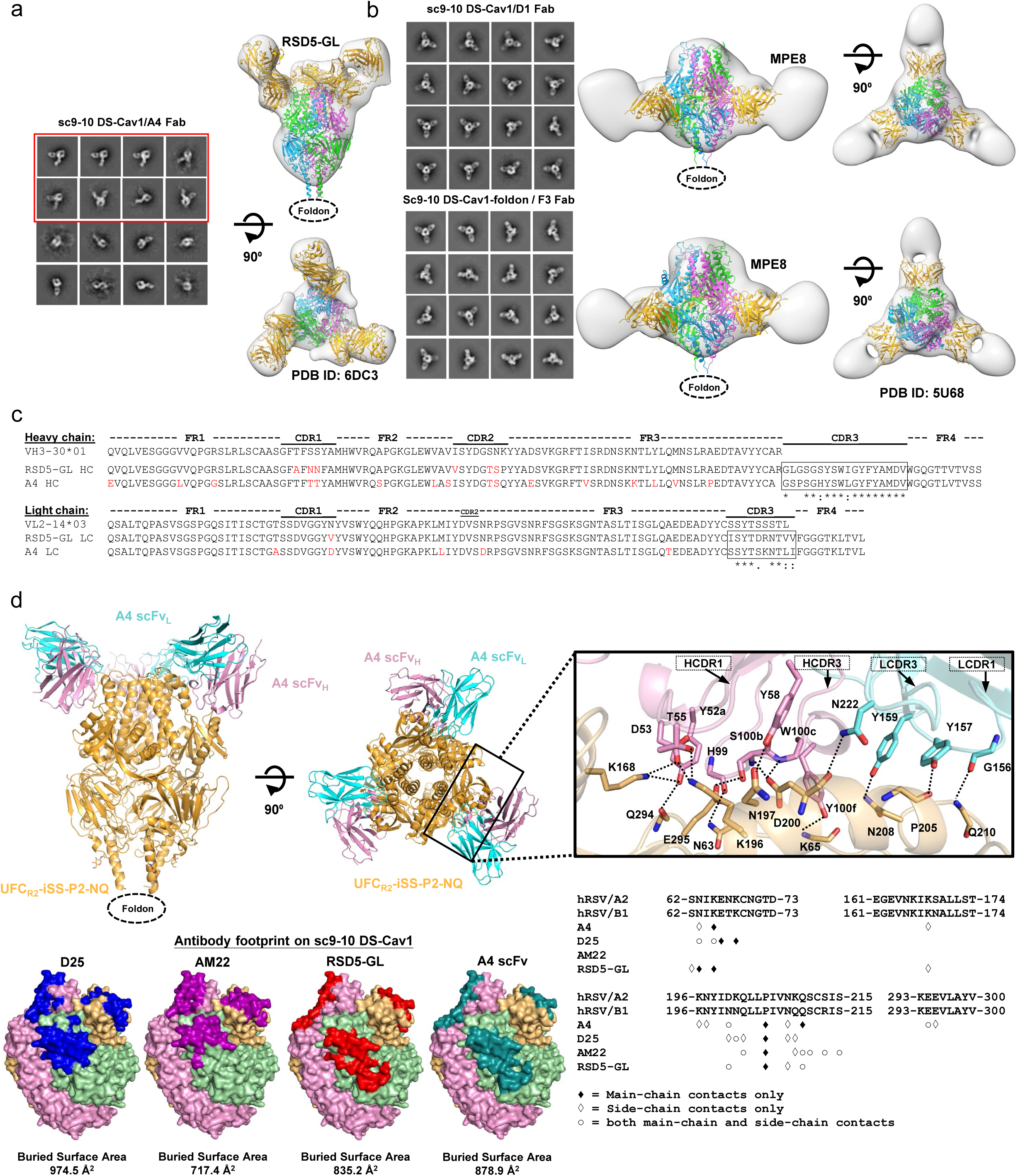
Structural characterization of potent pneumovirus neutralizing human antibodies. (a) The nsEM analysis of sc9-10 DS-Cav1 in complex with A4. Representative 2D classification images are shown on the left, and side and top views of 3D reconstruction of the complex are shown on the left. A 3.50 Å-resolution crystal structure of RSD5-bound DS-Cav1 (PDB ID: 6DC3) was used for density fitting. The 2D classification images containing two or more than two bound A4 Fabs are circled in a red line box. (b) The nsEM analysis of sc9-10 DS-Cav1 in complex with D1 and F3. Representative 2D classification images are shown on the left, and side and top views of 3D reconstruction of the complex are shown on the right. A 3.08 Å-resolution crystal structure of MPE8-bound DS-Cav1 (PDB ID: 5U68) was used for density fitting. (c) Sequence analysis of A4 and RSD5 heavy and light chains with alignment to respective germline genes. Mature antibody residues that differ from the germline are colored in red. (d) Crystallographic analysis of UFC_R2_-iSS-P2-NQ in complex with A4 scFv and structural epitope mapping. Top left: A 4.0 Å-resolution crystal structure UFC_R2_-iSS-P2-NQ in complex with A4 scFv is shown as ribbon models, with UFC_R2_-iSS-P2-NQ in gold and A4 heavy and light chains in pink and cyan, respectively. Top right: Close-up view of the A4/RSV-F interface. Side chains are shown for residues involved in hydrogen bond interactions across the A4/RSV-F interface, which were identified based on the estimated donor-acceptor distances and are indicated by dotted black lines. HCDR1, HCDR3, LCDR3 and LCDR1 loops are indicated. Bottom left: Surface models of prefusion RSV-F trimer showing footprints of D25, AM22, RSD5, and A4 colored in blue, rouge pink, red, and teal blue, respectively. Bottom right: RSV-F sequence with antibody-interacting residues labeled for D25, AM22, RSD5, and A4. Three types of contact are considered using a cutoff distance of 5 Å: main-chain contacts, side-chain contacts, and both.

Given the high potency of A4 and its similar angle of approach compared with RSD5 (*51*), we examined the gene families of these two NAbs (**Fig. 8c**). Consistent with the same binding and approach angle, A4 and RSD5 appeared to originate from the same V_H_-V_L_ combination (IGHV3-30*01-IGLV2-14*03), showing CDR3 sequence identities of 78% (14/18) and 50% (5/10) for HC and LC, respectively. This finding suggests that RSD5 and A4 may belong to a “public” antibody lineage targeting this unique epitope that overlaps both sites Ø and V. Similar public antibody lineages have been reported for SARS-CoV-2 (*88–90*). We obtained a crystal structure for A4 scFv-bound UFC_R2_-iSS-P2-NQ (**Table S7**). The RSD5-GL/DS-Cav1 structure (PDB ID: 6DC3) (*51*) was used as a template to build the A4/UFC_R2_-iSS-P2-NQ complex structure by molecular replacement, which was refined to a final model with a resolution of 4.0 Å (**Fig. 8d**, top left). At this resolution, charged or long aliphatic side chains could not be modeled as accurately as aromatic side chains. Nonetheless, our crystal structure revealed a tightly packed interface with more than 10 potential hydrogen bonds based on the estimated donor-acceptor distances (**Fig. 8d**, top right). Specifically, A4 employs HCDR1 (4 residues), HCDR3 (4 residues), LCDR1 (3 residues), and LCDR3 (1 residue) to interact with key residues of the RSV-F α4 helix (site Ø), such as K196, N197, D200, L204, P205, N208, and Q210 (n.b., the helix kinks at residue 203 prior to P205). Additional contacts were made with N63, K65 (β2-α1 loop, site Ø), K168 (α3), E294, and E295 (β5-β6 turn) of surrounding structural elements. The superimposition of A4/UFC_R2_-iSS-P2-NQ and RSD5-GL/DS-Cav1 using various fitting schemes revealed closely matched RSV-F α4 helices in site Ø and antibody HCDR3 loops (**Fig. S9a**). When the Cα atoms of α4 residues L195-L207 were used for fitting, we obtained a Cα-RMSD of 1.0 Å for HCDR3 (13 residues, H100-V102 for A4 and S100-V102 for RSD5-GL), which formed nearly identical interactions with RSV-F (**Fig. S9b**). Using the PDBePISA webtool (*91*), the A4 footprint on RSV-F was also compared with those of D25 (*42*), AM22, and RSD5-GL (*51*), showing the closest match to the RSD5-GL footprint (**Fig. 8d**, bottom left). Further analysis indicated that A4 appears to engage more residues on RSV-F than D25, AM22, and RSD5-GL, interacting with 12, 10, 7, and 8 residues, respectively, using both backbone and sidechain contacts (**Fig. 8d**, bottom right). Our structure thus provided critical insights into how A4 interacts with RSV-F to achieve a comparable potency to MEDI8897 (Nirsevimab) (*38, 86*).

### Antibody responses induced by rationally designed RSV-F trimer vaccines in mice

We assessed the immunogenicity of seven RSV-F constructs, including four UFC designs (UFC_R1_-P2-NQ/-L2 and UFC_R2_-P2-NQ/-L2) and three “control” designs (DS-Cav1Δp27 (*43*), SC-TM (*33*), and sc9-10 DS-Cav1 (*44*)), in BALB/c mice (**Fig. 9a**). In this in vivo study, the p27 peptide, which hinders F trimerization if not properly removed (*33*), was not included in DS-Cav1, resulting in a DS-Cav1Δp27 construct with a single cleavage site. Compared with Nickel/SEC-purified DS-Cav1-His_6_, D25/SEC-purified DS-Cav1Δp27 showed similar in vitro properties (**Fig. S10a-b**). Based on the antigenic data (**Fig. 1e** and **Fig. S10b**), D25/SEC-purified DS-Cav1Δp27 and sc9-10 DS-Cav1, along with Nickel/SEC-purified SC-TM-His_6_, were included as control antigens for comparison. A mouse immunization protocol used in our previous SARS-CoV-2, HIV-1, and influenza studies (*79, 92, 93*) was adopted with the number of animals per group increased to 10 to improve power in the statistical analysis. RSV-F antigens were formulated with AddaVax, an oil-in-water emulsion adjuvant, and administered intradermally through footpad injections (4 footpads, 2.5 μg/footpad) at 3-week intervals. Serum was isolated from blood draws obtained 2 weeks after each immunization. RSV-F binding antibody responses were determined using a sc9-10 DS-Cav1 1TD0 probe in the ELISA, with EC_50_ titers calculated for comparison (**Fig. 9b** and **Figs. S10a**-**S10b**). Overall, all groups demonstrated comparable binding antibody titers at each of the four studied time points except the SC-TM-His_6_ group, which showed significantly lower EC_50_ titers in most cases. This pattern was noted as early as week 2, where SC-TM-His_6_ showed a 7.1-28.7-fold lower EC_50_ titer than other antigens. Sc9-10 DS-Cav1 elicited the highest binding antibody titers at week 5, showing 3.6- and 46.1-fold higher EC_50_ titers than DS-Cav1Δp27 and SC-TM-His_6_, respectively. Among our four designs, UFC_R2_-P2-L2 showed the highest EC_50_ titer at week 5, which was 3.1- and 39.6-fold higher than DS-Cav1Δp27 and SC-TM-His_6_, respectively. Most mouse groups reached saturated EC_50_ titers after three immunizations at week 8. Interestingly, although UFC_R1_-P2-NQ and UFC_R1_-P2-L2 showed lower binding antibody titers than their UFC_R2_ counterparts at week 2, this pattern changed at week 5 and was reversed at weeks 8 and 11. Furthermore, L2 outperformed NQ regardless of the base construct, UFC_R1_ or UFC_R2_, at all four time points. These results correlated with the ratio of prefusion-closed trimers obtained from nsEM (**Figs. 1, 2**, and **4**). The week-11 sera were also analyzed against hMPV-F using the UFC_M1_-P2-iSS(1TD0) probe (**Fig. 9c** and **Fig. S10c**). A few mice in the DS-Cav1Δp27 and SC-TM-His_6_ groups showed nonspecific signals, as indicated by absorbance at 450 nm (A450).

**Fig. 9.**
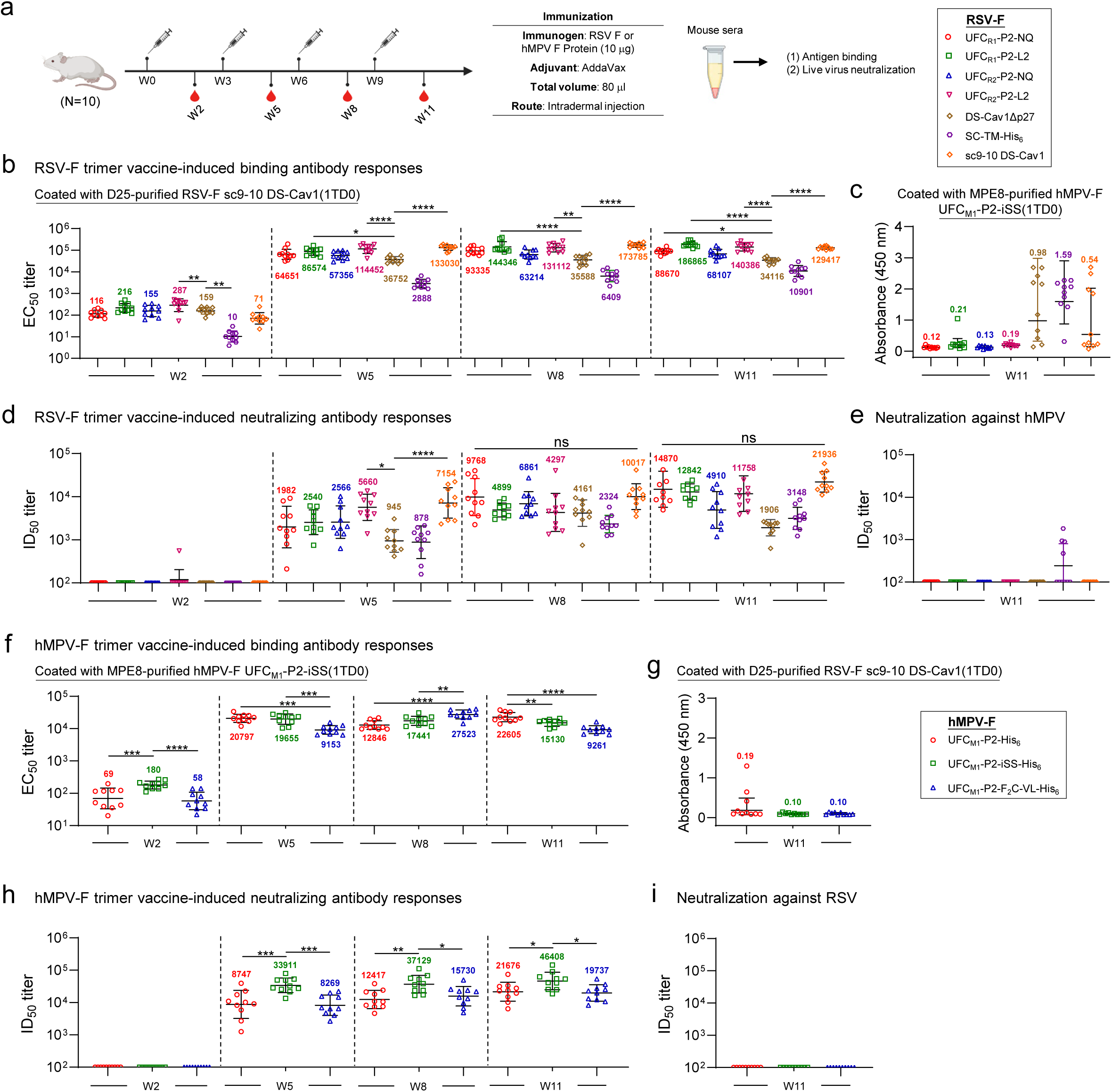
Antibody responses to rationally designed RSV and hMPV-F trimer vaccines in mice. (a) Schematic representation of the mouse immunization regimen for both RSV-F and hMPV-F vaccines (n = 10 mice/group). (b, c) RSV-F vaccine-induced binding antibody responses against RSV-F sc9-10 DS-Cav1(1TD0) and hMPV-F UFC_M1_-P2-iSS(1TD0). (d, e) RSV-F vaccine-induced neutralizing antibody responses against live RSV-A2-GFP and live hMPV-GFP. In (b)-(e), DS-Cav1Δp27, SC-TM-His_6_, and sc9-10 DS-Cav1 are included for comparison. (f, g) HMPV-F vaccine-induced binding antibody responses against hMPV-F UFC_M1_-P2-iSS(1TD0) and RSV-F sc9-10 DS-Cav1(1TD0). (h, i) HMPV-F vaccine-induced neutralizing antibody responses against live hMPV-GFP and live RSV-A2-GFP. EC_50_ values (b, c, f, g) were derived from the ELISA analysis of mouse serum against coating antigens, with geometric mean EC_50_ values labeled on the plots. ID_50_ titers were derived from the live RSV and hMPV neutralization assays, with geometric mean ID_50_ values labeled on the plots. Notably, the ID_50_ values were derived by setting the lower/upper constraints of % neutralization at 0.0/100.0. The data were analyzed using one-way ANOVA, followed by Tukey’s multiple comparison *post hoc* test for each timepoint. The statistical significance is indicated as the following: ns (not significant), \**p* < 0.05, \*\**p* < 0.01, \*\*\**p* < 0.001, and \*\*\*\**p* < 0.0001. In (b) and (d), statistical analyses of EC_50_ and ID_50_ values were performed by comparing individual RSV-F vaccine with the control, DS-Cav1Δp27. Detailed ELISA and neutralization data and the complete statistical analysis are shown in **Fig. S10.** The schematic representation of the mouse immunization protocol was created with BioRender.com.

Serum NAb responses were assessed using a live RSV neutralization assay (*94*), with the 50% inhibitory dilution (ID_50_) calculated for comparison. None of the RSV-F vaccines elicited a detectable NAb response at week 2 except for one mouse in the UFC_R1_-P2-L2 group, but the ID_50_ titers were detectable at weeks 5, 8, and 11 and continued to increase throughput these timepoints (**Fig. 9d** and **Fig. S10d**-**S10e**). Consistent with the binding antibody titers, the DS-Cav1Δp27 and SC-TM-His_6_ groups showed the lowest NAb titers across all timepoints. In contrast, sc9-10 DS-Cav1 yielded the highest NAb titers of 7154, 10017, and 21936 at weeks 5, 8, and 11, respectively, which were 7.6-, 2.4-, and 11.5-fold higher than the NAb titers elicited by DS-Cav1Δp27. Among our four designs, the UFC_R1_-P2-NQ group showed the highest ID_50_ titer at week 8, which was comparable to the sc9-10 DS-Cav1 group. At weeks 8 and 11, the two UFC_R2_ designs consistently underperformed their UFC_R1_ counterparts in terms of ID_50_ titers. These results confirmed that a high ratio of prefusion-closed trimers (e.g., 73% for UFC_R1_-P2-NQ) (**Fig. 2c**) may benefit the elicitation of potent NAbs but also demonstrated that the L2 mutation, which transiently exposes site I, may dampen NAb responses (**Figs. 2f** and **4f**). Lastly, the week-11 serum samples were tested using a live hMPV neutralization assay (*95*) (**Fig. 9e** and **Fig. S10f**). All groups showed negligible NAb responses against hMPV except SC-TM-His_6_, likely caused by the weak NAbs induced by the postfusion form of this antigen.

Our results highlighted the advantage of stabilized, prefusion-closed RSV-F trimer in NAb elicitation. The first-generation DS-Cav1 and SC-TM induced low NAb titers in these experiments. Overall, sc9-10 DS-Cav1 appeared to be the best performer among all RSV-F constructs, with UFC_R1_-P2-NQ ranked the second-best performer at most time points in the immunization.

### Antibody responses induced by rationally designed hMPV-F trimer vaccines in mice

The immunogenicity of three hMPV-F design constructs, UFC_M1_-P2, UFC_M1_-P2-iSS, and UFC_M1_-P2-F_2_C-VL, was assessed in mice (**Fig. 9a**). The hMPV-F-specific binding antibody responses were measured by ELISA using a UFC_M1_-P2-iSS(1TD0) probe for all timepoints. The EC_50_ titers were calculated and plotted longitudinally for comparison (**Fig. 9f** and **Fig. S10g**-**S10h**). Overall, all groups demonstrated strong binding antibody responses, with EC_50_ values ≥ 9153 after two vaccine doses. The UFC_M1_-P2 group reached the highest EC_50_ titer at week 5, comparable to the UFC_M1_-P2-iSS group and 2.3-fold higher than the UFC_M1_-P2-F_2_C-VL group. Interestingly, the UFC_M1_-P2-F_2_C-VL group yielded higher EC_50_ values than the other two groups at week 8. Cross-reactive antibody responses were assessed using week-11 sera against an RSV-F sc9-10 DS-Cav1 1TD0 probe (**Fig. 9g** and **Fig. S10i**). All three groups showed negligible signals except some nonspecific signals observed for the UFC_M1_-P2 group. Live hMPV neutralization assays (*95*) were used to assess serum NAb responses elicited by various hMPV-F constructs (**Fig. 9h** and **Fig. S10j**-**S10k**). As expected, none of the three hMPV-F constructs elicited NAb responses against autologous hMPV at week 2. Importantly, the UFC_M1_-P2-iSS group showed the most potent NAb titers, with ID_50_ titers of 33911, 37129, and 46408, which were 3.9-, 3.0-, and 2.1-fold higher than those elicited by UFC_M1_-P2 at weeks 5, 8, and 11, respectively. This confirmed the importance of our rationally designed mutations for producing a stabilized, prefusion-closed hMPV-F trimer, which in turn induced a potent NAb response against hMPV. We evaluated the cross-neutralizing activity of hMPV-F-induced mouse sera using live RSV assays, which did not detect NAb titers against RSV (**Fig. 9i** and **Fig. S10**l). Altogether, our results indicate that UFC_M1_-P2-iSS, as a fully closed prefusion hMPV-F trimer, can induce a potent NAb response in mice.

## DISCUSSION

Both RSV and hMPV cause LRT infections in infants, young children, and the elderly (*1, 2*). Although RSV-F and HMPV-F share only ∼30% sequence identity, their F proteins are structurally similar in both prefusion and postfusion states (*96*). Over the last decade, at least four rational RSV-F designs have been reported and extensively characterized in vitro and in vivo (*33, 43–45*). Some of these structurally optimized prefusion RSV-F constructs gave rise to the recently licensed RSV vaccines: ABRYSVO™ (GSK) (*54*) and AREXVY (Pfizer) (*45, 97*). Inspired by the success of these RSV-F designs, similar structure-based design principles have been applied to hMPV-F to stabilize the prefusion trimer and facilitate vaccine development (*60–62*). However, outcomes from these studies suggest that rational hMPV-F design is still in its infancy. As a result, there are no rationally designed hMPV-F trimer vaccines approved for human use.

The recently approved RSV vaccines have been celebrated as a success of structure-based rational vaccine design (*41*), but the cause of RSV-F metastability largely remains unclear. In addition, an in-depth comparative analysis of previous RSV-F designs has not been reported. Our study addressed these issues with a systematic approach. Notably, use of the site Ø-specific NAb D25 (*42*) for purification allowed us to quantify and extract the prefusion fraction of RSV-F from mammalian cell expression. Both SC-TM and 847A had a low yield using a D25 NAb column but not a Nickel affinity column, suggesting the presence of non-prefusion F species. While x-ray crystallography validated our RSV-F designs in atomic detail, nsEM was extensively used to probe RSV-F structures and fusion states. Indeed, nsEM demonstrated that His-tagged SC-TM and 847A contained detectable postfusion F trimers after Nickel/SEC purification. This finding is rather concerning because a postfusion F vaccine failed to prevent RSV illness in older adults (*98*). Another key element in our study was the use of a pair of antibodies, prefusion site Ø-specific NAb D25 (*42*) and postfusion site I-specific non-NAb ADI-14359 (*48*), to probe fusion states of an RSV-F construct. Indeed, ADI-14359 in the ELISA confirmed the presence of postfusion F in Nickel/SEC-purified SC-TM-His_6_ and 847A-His_6_ samples. Overall, sc9-10 DS-Cav1 (*44*) appeared to be the best of the four RSV-F designs. Our results also revealed two forms of RSV-F metastability: a tendency to undergo a rapid pre-to-postfusion change and a tendency for prefusion trimers to open or dissociate in solution. The F_2_-F_1_ junction appeared to be another key contributor to metastability, as sc9-10 DS-Cav1 differs the most from others in this region.

Systematic analyses of various RSV-F designs based on the UFC_R1_, UFC_R2_, and UFC_R3_ series allowed us to examine a set of diverse design principles and identify alternate mutations to the interprotomer disulfide bond for maintaining prefusion RSV-F in a closed trimer. A number of important conclusions were drawn. First, an RSV-F construct with limited mutations can produce prefusion (albeit open) trimers with high yield and high purity, as indicated by UFC_R1_. The success of this “minimalist approach” provided a base construct to investigate RSV-F metastability and assess various solutions. Second, the buried acidic patch (^485^DEFD^488^) in β23 appears to trigger the prefusion RSV-F trimer to open because of the repulsive charge-charge interactions atop the trimeric α10 stalk. Either a hydrogen bond (NQ) or hydrophobic (L2) mutation could effectively maintain up to 76% of the RSV-F protein in a prefusion-closed conformation. Third, this acidic patch is critical to both forms of metastability and responds to remote mutations. For example, a hydrophobic (L2) mutation of this acidic patch likely results in breathing motions of RSV-F that transiently expose site I in both UFC_R1_ and UFC_R2_ contexts, suggesting that the two forms of metastability are intrinsically connected through this acidic patch. In addition, S46G and K465Q mutations in the UFC_R2_ base, more than 35 Å away from the acidic patch, drastically reduced the stabilizing effect of NQ and L2 mutations. Fourth, the β3-β4 hairpin in RSV-F, which is equivalent to HR1_N_ – the fundamental cause of HIV-1 Env metastability (*70, 71*), has little impact on RSV-F metastability. The V185P (P2) mutation neither improved nor worsened any RSV-F properties in the UFC_R1_ and UFC_R2_ series. A similar pattern was observed for the interstrand disulfide bond (A177C-T189C), which was intended to lock the prefusion β3-β4 hairpin conformation. The extensive crystallographic analyses, in concert with other experimental approaches, validated our hypotheses on structural designs and RSV-F metastability at the atomic level.

Despite structural similarity to RSV-F, hMPV-F appears to not be confined by the same metastability principles, presenting a new challenge for antigen design and potentially explaining the suboptimal outcomes for recent hMPV-F designs (*60–62*). First, RSV-F and hMPV-F differ significantly around the F_2_-F_1_ region, with a p27 peptide and double cleavage site in RSV-F *vs.* lack of the p27 peptide and a single cleavage site in hMPV-F, respectively, resulting in different sensitivity of the prefusion F design to truncation or linkage in the F_2_-F_1_ region. In our base design, UFC_M1_, a short truncation at the F_2_ C-terminus coupled with a G_6_ linker resulted in prefusion-open trimers. However, a larger truncation in the F_2_-F_1_ region would produce postfusion hMPV-F trimers, as demonstrated by v3B_Δ18 and v3B_Δ15 (*60*). Second, unlike RSV-F, for which the P2 mutation displays no visible effect, the equivalent mutation (A185P) improves the expression of prefusion hMPV-F (albeit being open trimers). Third, unlike RSV-F, which contains an acidic patch essential to metastability, hMPV-F has a non-acidic ^454^DQFN^457^ segment in β23, suggesting a different mechanism to trigger the prefusion trimer to open. Although an interprotomer disulfide bond in UFC_M1_-P2-iSS could effectively lock hMPV-F in a prefusion-closed trimer, our attempt to replace it with a non-covalent interaction proved unsuccessful, resulting in a mixture of F species in UFC_M1_-P2-F_2_C-VL. Compared with the two recent designs (*60, 61*), UFC_M1_-P2-iSS provides a simple and robust construct to produce prefusion-closed trimers for hMPV-F vaccine development. However, the causes of hMPV-F metastability warrant more in-depth investigations.

The biopanning of a phage antibody library provided a direct in vitro approach to evaluate our top RSV-F design. Our hypothesis was that a prefusion-closed RSV-F trimer should select out both prefusion- and trimer-specific NAbs from a human antibody library that contains infection-induced RSV NAbs. Indeed, this was confirmed by our functional and structural characterization of a subset of scFv clones. A4 represents a “public” antibody lineage targeting a prefusion-specific epitope that overlaps sites Ø and V like RSD5 (*51*), showing the highest IgG potency on par with MEDI8897 (Nirsevimab) (*86*). D1 and F3 are MPE8-like NAbs directed to site III at the protomer interface (*63*), with F3 IgG demonstrating comparable potency to D25 against RSV (*42*) and 10-times higher potency than MPE8 against hMPV. We performed a mouse immunization study to assess antibody responses to various RSV-F and hMPV-F constructs. Overall, our serological analysis revealed a strong and positive correlation between binding and NAb responses. Among all RSV-F constructs, DS-Cav1 (*43*) and SC-TM (*33*) elicited the lowest EC_50_ and ID_50_ titers at all timepoints, which correlated with trimer dissociation and the presence of postfusion F, respectively. The second-generation sc9-10 DS-Cav1 elicited the highest binding antibody and NAb titers. For our four RSV-F constructs, serum binding and NAb responses correlated with their *in vitro* and structural properties, thus confirming our design hypotheses. Within our three hMPV-F constructs, UFC_M1_-P2-iSS elicited the most potent NAb responses with the highest ID_50_ titers at all timepoints, stressing the importance of a prefusion-closed conformation to NAb elicitation. Interestingly, some RSV-F-immunized mouse samples showed heterologous NAb responses against hMPV.

Our future studies may focus on the following directions. First, more in-depth examination of metastability, as well as construct optimization, are required for hMPV-F. The introduction of an interprotomer disulfide bond led to a reduced trimer yield for UFC_M1_-P2-iSS. Thus, it remains imperative to understand the causes of hMPV-F metastability and design alternative mutations to maintain a prefusion-closed hMPV-F trimer. Second, the optimized RSV-F and hMPV-F trimers can be displayed on our single-component, self-assembling protein nanoparticles (1c-SApNPs) to further improve immunogenicity and NAb elicitation. DS-Cav1 has been displayed on protein NPs for vaccine development (*75, 76*). We have successfully designed 1c-SApNP vaccines for HIV-1 (*70, 93, 99*), HCV (*100*), EBOV (*69*), SARS-CoV-2 (*68, 92*), and influenza (*79*). Lastly, a bivalent RSV-F/hMPV-F vaccine may warrant careful evaluation. Although monovalent immunogens did not elicit heterologous NAb responses in our mouse study, a combination of optimized RSV-F and hMPV-F UFC trimers might generate an effective cross-pneumovirus NAb response.

## METHODS

### Design, expression, and purification of RSV-F and hMPV-F

The amino acid sequences of RSV-F and hMPV-F were based on subtype A (strain A2, UNIPROT ID: P03420) and strain NL/1/00 (A1 sublineage, UNIPROT ID: Q1A2Z0), respectively. Structural modeling was performed using UCSF Chimera v1.13 software. All redesigned RSV-F and hMPV-F constructs were subcloned into a CMV/R vector and transiently expressed in ExpiCHO cells (Thermo Fisher, catalog no. A29133) as previously reported (*69, 93*). The cells were thawed and incubated with ExpiCHO^TM^ Expression Medium (Thermo Fisher) in a shaker incubator at 37°C, 135 rpm, and 8% CO_2_. The cells were allowed to reach a cell density of 1 × 10^7^/ml. On the day of transfection, the cells were diluted to 6 × 10^6^/ml in ExpiCHO^TM^ Expression Medium, after which ExpiFectamine^TM^ CHO/plasmid DNA complexes were prepared for 25-ml transfection according to the manufacturer’s instructions. For all RSV-F and hMPV-F constructs tested in this study, 25 μg F antigen plasmid and 80 μl of ExpiFectamine^TM^ CHO reagent were mixed in 1.9 ml of cold OptiPRO™ medium (Thermo Fisher). After the first feed on day 1, ExpiCHO cells were further cultured in the shaker incubator at 32°C, 120 rpm, and 8% CO_2_ following the Max Titer protocol, with an additional feed on day 5 (Thermo Fisher). Culture supernatants were harvested 12-14 days after transfection, clarified by centrifugation at 3,724 × g for 25 min, and filtered using a 0.45-μm filter (Millipore). Some RSV-F and hMPV-F constructs were purified using D25 (*42*) and MPE8 (*63*) NAb columns, respectively, whereas others were purified using Ni Sepharose excel resin (Nickel, Cytiva) as an alternative approach. For NAb columns, bound proteins were eluted three times, each with 5 ml of 0.2 M glycine (pH 2.2) and neutralized with 0.5 ml of Tris-Base (pH 9.0), and then exchanged into phosphate-buffered saline (PBS; pH 7.2). For Nickel columns, bound proteins were eluted two times, each with 15 ml of 0.5 M imidazole, and then exchanged into Tris-buffered saline (TBS; pH 7.2). The protein samples were then further purified by size exclusion chromatography (SEC) using a Superdex 200 Increase 10/300 GL column (Cytiva) mounted on an AKTA pure system (Cytiva). All SEC data were collected using UNICORN 7.5 software. Protein concentrations were determined by UV_280_ with theoretical extinction coefficients.

### Antibody expression and purification

All antibodies in IgG and Fab forms in this study were transiently expressed in ExpiCHO cells (Thermo Fisher). For IgGs and Fabs, 12-14 days post-transfection, the cells were centrifuged at 3,724 × g for 25 min, and the supernatants were filtered using a 0.45-μm filter (Millipore). IgGs were further purified using protein A affinity resin (Cytiva) and eluted in 0.3 M citric acid (pH 3.0), and Fabs were purified using Protein G affinity resin (Cytiva) and eluted in 0.2 M glycine (pH 2.0). The pH of the elution was immediately titrated to a neutral pH of 7.0 by addition of 2 M Tris-Base (pH 9.0). The eluate was concentrated and exchanged into phosphate buffered saline (PBS) using an Amicon 10 kDa filter (Millipore). The antibody concentration was quantified by UV_280_ absorption with theoretical extinction coefficients.

Phage library-derived scFv-His and scFv-Fc antibodies were all transiently expressed in HEK293F (293F) cells using FreeStyle^TM^ 293 Expression Medium. Briefly, HEK293F cells were thawed and incubated with FreeStyle^TM^ 293 Expression Medium (Life Technologies, CA) in a shaker incubator at 37°C at 135 rpm with 8% CO_2_. When the cells reached a density of 2.0 × 10^6^/ml, expression medium was added to reduce the cell density to 1.0 × 10^6^/ml for polyethyleneimine (PEI; Polysciences) transfection. For 50-ml HEK293F transfection, 45 μg expression plasmid in 1.25 ml of Opti-MEM transfection medium (Life Technologies, CA) was mixed with 1.25 ml of OptiMEM containing 0.5 ml of PEI-MAX (1.0 mg/ml). After 25 min of incubation, the DNA-PEI-MAX complex was added to HEK293F cells. Culture supernatants were harvested 5 days after transfection and centrifuged at 3,724 × g for 15 min. Nickel and protein A resins were then used to purify scFv-His and scFv-Fc antibodies, respectively, from the clarified supernatant. Protein was then concentrated, and buffer-exchanged into PBS using an Amicon 10 kDa filter. Purified antibodies were quantified using NanoDrop and stored at -80°C until use.

### Crosslinking of RSV-F and hMPV-F trimers in SDS-PAGE

SEC-purified RSV-F and hMPV-F proteins were cross-linked by disuccinimidyl glutarate (DSG) (Thermo Scientific, Catalog No. 20593) and analyzed by SDS-PAGE under reducing conditions. Briefly, 5 μg of each protein was prepared in 20 μl of PBS and mixed with 5 μl of 25 mM DSG in dimethylsulfoxide (DMSO). The mixture was incubated at room temperature for 30 min before the addition of 1.25 μl of quench buffer (1 M Tris-HCl, pH 7.0) and further incubated for 10 min. Next, 5 μl of 6× loading buffer was added to the sample and heated at 100°C for 5 min. Thereafter, 12 μl of sample was loaded into a 4-20% Mini-PROTEAN TGX^TM^ PRECAST GEL (Bio-Rad). SDS-PAGE gels were then run for 50 min at 150 V in 1× Tris/glycine/SDS running buffer (Bio-Rad, Catalog No. 1610732). Lastly, the gels were stained using InstantBlue Coomassie protein stain (Abcam) for 2 h and imaged by a ChemiDoc MP Imaging System (Bio-Rad).

### Differential scanning calorimetry

Thermal melting curves of all RSV-F and hMPV-F proteins following antibody/Nickel column and SEC purification were obtained from a MicroCal PEAQ-DSC Man instrument (Malvern). Briefly, purified proteins in PBS buffer were adjusted to 1-5 μM before analysis. Melting was probed at a scan rate of 60°C/h from 20°C to 100°C. Data processing, including buffer correction, normalization, and baseline subtraction, was conducted using MicroCal PEAQ-DSC software. Gaussian fitting was performed using GraphPad Prism 10.0.2 software.

### Enzyme-linked immunosorbent assay (ELISA)

Costar^TM^ 96-well, high-binding, flat-bottom, half-area plates (Corning) were first coated with 50 µl of PBS containing 0.1 μg of the appropriate RSV-F and hMPV-F antigens. The plates were incubated overnight at 4 °C and then washed five times with PBST wash buffer containing PBS and 0.05% (v/v) Tween 20. Each well was then coated with 150 µl of blocking buffer consisting of PBS and 40 mg/ml blotting-grade blocker (Bio-Rad). The plates were incubated with the blocking buffer for 1 h at room temperature, and then washed five times with PBST wash buffer. For antigen binding, antibodies (in either IgG or scFv form) were diluted in blocking buffer to a maximum concentration of 10 μg ml^-1^ followed by a 10-fold dilution series. For each antibody dilution, a total volume of 50 μl was added to the appropriate wells. For mouse sample analysis, serum was diluted 40-fold in blocking buffer and subjected to a 10-fold dilution series. For each sample dilution, a total volume of 50 μl was added to the wells. Each plate was incubated for 1 h at room temperature and then washed five times with PBST wash buffer. For antibody binding, a 1:5000 dilution of goat anti-human IgG antibody (Jackson ImmunoResearch Laboratories), or for mouse sample analysis, a 1:3000 dilution of horseradish peroxidase (HRP)-conjugated goat anti-mouse IgG antibody (Jackson ImmunoResearch Laboratories), was then made in the PBST wash buffer, with 50 μl of this diluted secondary antibody added to each well. The plates were incubated with the secondary antibody for 1 h at room temperature, and then washed six times with PBST wash buffer. Finally, the wells were developed with 50 μl of 3,3’,5,5’-tetramethylbenzidine (TMB; Life Sciences) for 3-5 min before stopping the reaction with 50 μl of 2 N sulfuric acid. The plates were immediately read on a BioTek Synergy plate reader at a wavelength of 450 nm. EC_50_ values were then calculated from full curves using GraphPad Prism 10.0.2 software. When antibody binding signals (OD_450_) were lower than 0.5, the binding was considered weak and the EC_50_ values were set to 10 µg/ml in **Figs. 1e**, **2f**, and **4f** to facilitate EC_50_ plotting and comparison.

### Bio-layer interferometry

Antigenic profiles of RSV-F and hMPV-F were measured using Octet RED96 (FortéBio, Pall Life Sciences) against a panel of 10 antibodies for RSV-F and four antibodies for hMPV-F, all in IgG forms. All assays were performed with agitation set to 1000 rpm in FortéBio 1× kinetic buffer. The final volume for all solutions was 200 μl per well. Assays were performed at 30 °C in solid black 96-well plates (Geiger Bio-One). For all antigens, 5 μg/ml antibody in 1× kinetic buffer was loaded onto the surface of anti-human Fc Capture Biosensors (AHC) for 300 s. Next, a 2-fold concentration gradient of antigen, starting at 600 nM, was used in a dilution series of six. A 60-s biosensor baseline step was applied before the analysis of association of the antibody on the biosensor to the antigen in solution for 200 s. Dissociation of the interaction was followed for 300 s, after which the correction of baseline drift was performed by subtracting the mean value of shifts recorded for controls: a sensor loaded with antibody + no antigen and a sensor without an antibody + an antigen. Octet data were processed by FortéBio’s data acquisition software v.8.1. Peak signals at the highest antigen concentration were summarized in a matrix and color-coded accordingly to allow comparisons between different constructs. Experimental data for each antigen-antibody pair were fitted with the binding equations describing a 1:1 interaction, and three datasets that showed the optimal fitting were then grouped to determine the K_on_ and K_D_ values.

### Protein crystallization and data collection

Freshly purified RSV-F proteins were used for crystallization experiments using the sitting drop vapor diffusion method on a Cryschem M Plate (Hampton Research) with protein concentrations between 5 and 10 mg/ml. For hMPV-F UFC_M1_-P2-iSS-foldon-His6 and RSV-F sc9-10 DS-Cav1-foldon complexes with A4 scFv, ∼ 8 mg/ml protein was used for crystallization experiments using the sitting drop vapor diffusion method using our automated CrystalMation robotic system (Rigaku) at 20°C at The Scripps Research Institute (*101*). The reservoir solution details are listed in **Table S8**. Diffraction-quality crystals were obtained after 2 weeks at 20°C. All RSV-F crystals were cryoprotected with a well solution containing 25% (*v/v*) ethylene glycol, mounted on a nylon loop, and flash cooled in liquid nitrogen. Diffraction data were collected for all RSV-F crystals at Stanford Synchrotron Radiation Lightsource (SSRL) beamlines 12-1 and 12-2 and processed with HKL-2000 (*102*). Diffraction data for all RSV-F designs were indexed in P4_3_21, hMPV-F UFC_M1_-P2-iSS-foldon-His6 in I213, and the complex of UFC_R2_-iSS-P2-NQ with A4 scFv in R32.

### Structure determination and refinement

Structures of all RSV-F constructs were determined by molecular replacement (MR) using Phaser-MR from Phenix (version 1.19.2-4158) (*103*) with coordinates of the uncleaved prefusion structure of sc9-10 DS-Cav1 (PDB ID: 5K6I). The hMPV-F UFC_M1_-P2-iSS-foldon-His_6_ structure was determined by molecular replacement (MR) using prefusion hMPV-F (PDB ID: 5WB0) as the MR model. The structure of UFC_R2_-iSS-P2-NQ in complex with A4 scFv was determined by molecular replacement using the prefusion RSV-F in complex with RSD5 Fab (PDB ID: 6DC3). Polypeptide chains were manually adjusted into electron density using Coot (*104*), refined with phenix-refine (*103*), and validated using the wwPDB Validation System (*105*). The final data processing and refinement statistics are described in **Tables S2-S4**, **S6**, and **S7**. All images for crystal structures shown in the figures were generated using PyMOL v2.3.4 software.

### Negative-stain electron microscopy (nsEM)

The nsEM analysis of RSV-F and hMPV-F trimers, as well as their antibody-bound complexes, was performed by the Core Microscopy Facility at The Scripps Research Institute. Briefly, samples were prepared at a concentration of 0.008 mg/ml. Carbon-coated copper grids (400 mesh) were glow-discharged, and 8 µl of each sample was then adsorbed for 2 min. Next, excess sample was removed, and grids were negatively stained with 2% uranyl formate for 2 min. Excess stain was wicked away and the grids were allowed to dry. Samples were then analyzed at 120 kV with a Talos L120C transmission electron microscope (Thermo Fisher), and images were acquired using a CETA 16 M CMOS camera. Computational analysis of the imaging data was performed using the high-performance computing core facility at The Scripps Research Institute. The nsEM images were processed using EMAN2 (*106*) and cryoSPARC v4.3.0 (*107*). Briefly, micrographs were contrast transfer function (CTF) corrected, and particles were selected using a Blob/template picker and later extracted for 2D classification. Three-dimensional (3D) models were generated by *ab initio* reconstruction and refined by heterogeneous and homogeneous refinement. All nsEM and fitted structure images were generated by UCSF Chimera (*108*) and Chimera X (*109–111*).

### Preparation and biopanning of a healthy donor antibody phage library

Peripheral blood mononuclear cells (PBMC) from 10 healthy donors were purchased from iXCells Biotechnologies (San Diego, CA) and used for scFv library construction following our previously described method (*85*). Briefly, total RNA was extracted from ∼10 million PBMCs for single-stranded cDNA synthesis using SuperScript^TM^ IV (Invitrogen) with random hexamer and oligo(dT) 12-18 primers. Antibody heavy chain (HC) and light chain (LC) variable regions were obtained by polymerase chain reaction (PCR) using mixed HuJ reverse primers and separate HuV forward primers (*85*). To generate HC-LC fragments, overlap PCR was performed in 25 × 50 μl reactions (10 cycles) with 50 ng of gel-purified HC and 50 ng of gel-purified LC (equal amounts of κ and λ LC) without primer. Subsequent PCR was performed in 50 × 50 μl reactions (15 cycles) with SfiI-F and SfiI-R primers using 100 ng of gel-purified HC-LC as a template to obtain full-length scFv inserts. The scFv inserts and phagemid vector, pAdLTM-20c (Antibody Design Labs), were then digested with SfiI and purified through gel extraction. The scFv DNA (256 ng) was ligated into the phagemid vector (400 ng) using the T4 Ligase Kit (New England BioLabs) in 25 × 40 μl reactions and incubated at 16°C overnight. After purification, the phagemids were electroporated into phage-competent TG1 bacterial cells (Lucigen) with the MicroPulsuer^TM^ system (Bio-Rad). Specifically, 1 μl of phagemid and 25 μl of competent cells were incubated in a 0.1 cm cuvette for electroporation with the preset program at 1.8 kV. The transformed bacteria were spread on 2YT agar plates supplemented with 100 μg/ml carbenicillin and 2% (w/v) glucose, which were then incubated at 37°C overnight. Bacteria were then scraped from the plates for phage culture with the helper phage CM13 (Antibody Design Labs) for biopanning. The purified UFC_R1_-P2-NQ trimer was conjugated with Dynabeads^TM^ M-270 beads before biopanning. A total of four biopanning steps were performed, incorporating up to 5 washes in each step to eliminate phages that were either not bound or showed low affinity for the RSV-F trimer. Lastly, plasmids from each biopanning step were extracted from 50 ml of the bacterial suspension and used for the subsequent deep sequencing and bioinformatics analyses of scFv libraries.

### Phage enzyme-linked immunosorbent assay

Phage ELISA was performed to select scFv clones reactive to sc9-10 DS-Cav1 using a previously described protocol (*112*). Briefly, wells of Costar^TM^ 96-well assay plates (Corning) were coated with 50 μl of PBS containing 0.2 μg sc9-10 DS-Cav1 and incubated overnight at 4°C. The next day, the plates were washed five times with PBST wash buffer and blocked with 150 μl/well of nonfat milk blocking buffer for 1 h at room temperature. The plates were then washed five times, and 50 μl of scFv-displaying phage was added to the appropriate wells. Each plate was incubated for 1 h and then washed five times with PBST wash buffer. Next, a 1:5,000 dilution of HRP-conjugated mouse anti-M13 antibody (Sino Biological) was made in the wash buffer, and 50 μl of diluted antibody was added to each well. The plates were incubated for 1 h and washed five times with PBST wash buffer. Finally, the wells were developed with 50 μl of TMB for 5 min before the reaction was stopped with 50 μl of 2 N sulfuric acid. The plate readout was performed as described above for the ELISA analysis of antigen-antibody interactions. RSV-F reactive scFv clones were subjected to Sanger sequencing (Eton Bioscience) to extract nucleotide sequences.

### Next-generation sequencing (NGS) and bioinformatics analysis

Two pairs of primers were used to generate NGS libraries. To amplify HC fragments, a forward primer containing an A adaptor, an Ion Xpress^TM^ barcode (Life Technologies), and a cloning site-targeting motif, and a reverse primer containing a P1 adaptor and a GS linker-targeting motif, were used (listed in **Fig. S8f**). To amplify V_L_ or V_K_ fragments, a forward primer containing a P1 adaptor and a GS linker-targeting motif, and a reverse primer containing an A adaptor, an Ion Xpress^TM^ barcode (Life Technologies), and a cloning site-targeting motif were used (**Fig. S8f**). One scFv library at each step of the biopanning process (pre-panning, Pan-0, Pan-1, Pan-2, and Pan-4) would produce three antibody chain libraries (V_H_, V_L_, and V_K_). Three antibody chain libraries from each biopanning step (with the same barcode) were then quantitated using a Qubit® 2.0 Fluorometer with the Qubit®dsDNA HS Assay Kit and pooled. Finally, the five pooled libraries were mixed at a ratio of 5:1:1:1:1 for further processing. Template preparation and Ion 530 chip loading were performed on Ion Chef using the Ion 520/530 Ext Kit, followed by a single NGS run on the Ion GeneStudio S5 system, as previously described (*113*). The Antibodyomics 2.0 pipeline (*113*) was used to process the raw NGS data and determine distributions for germline gene usage, somatic hypermutation, germline divergence, and CDR3 length. The 2D identity-divergence plots were used to visualize selected scFv lineages in the context of the antibody library at each biopanning step. The CDR3 identity of 95% and a 1-residue error margin of CDR length calculation were used to identify potential somatic variants of a reference antibody clone. These variants are shown as orange dots on the 2D plots (**Fig. 7f** and **Fig. S8h**).

### Animal immunizations and sample collection

We utilized a similar immunization protocol from our previous vaccine studies (*79, 92, 93*). All animal experiments followed the Association for the Assessment and Accreditation of Laboratory Animal Care (AAALAC) guidelines and used protocols that were approved by the Institutional Animal Care and Use Committee (IACUC) of The Scripps Research Institute (TSRI). Six-week-old female BALB/c mice were purchased from The Jackson Laboratory and housed in ventilated cages in environmentally controlled rooms in the Immunology building of The Scripps Research Institute. For the immunogenicity study, the mice were injected at weeks 0, 3, 6, and 9 with 80 μl of antigen/adjuvant mix containing 10 μg of vaccine antigen in 40 μl of PBS and 40 μl of AddaVax adjuvant (InvivoGen). Intradermal immunization in mice was performed through injections into four footpads, each with 20 μl of antigen/adjuvant mix using a 29-gauge insulin needle under 3% isoflurane anesthesia with oxygen. Blood was drawn from the maxillary/facial vein 2 weeks after each immunization. Serum was isolated from blood after centrifugation at 14000 rpm for 10 min. Serum was then heat-inactivated at 56°C for 30 min and the supernatant was collected after centrifugation at 8000 rpm for 10 min. Heat-inactivated serum was used in antigen binding and neutralization assays to determine vaccine-induce antibody responses.

### Virus production and neutralization assays

The propagation of viruses and microneutralization assays for RSV and hMPV were based on previous protocols with modification (*94, 114, 115*). The following reagents were obtained from ViraTree: RSV-A2-GFP and hMPV-GFP (parental strain CAN97-83). For the propagation of RSV-A2-GFP, 3.0 × 10^6^ HEp-2 cells (American Type Culture Collection [ATCC], Catalog No. CCL-23) cells were plated in 100 cm cell culture dishes. The next day, cells were washed twice with PBS and infected with a multiplicity of infection (MOI) between 0.01 and 1 of RSV-A2-GFP for 2 h. Next, 10 ml of growth media (2.5% fetal bovine serum [FBS] in Dulbecco’s Modified Eagle Medium [DMEM]) was added to the plates, which were then incubated for 5 days at 37°C. On the day of harvest, cells were scraped, resuspended in viral growth media, and centrifuged at 4000 rpm for 10 min at 4°C. Equal volumes of viral supernatant and 50% w/v sucrose in PBS were combined, aliquoted, and frozen at -80°C until use. For hMPV propagation, 3.0 × 10^6^ VeroE6 cells (ATCC Catalog No. CRL-1586) were plated in 100 cm cell culture dishes overnight. The next day, cells were washed twice with PBS and infected with a MOI between 0.01 and 1 of hMPV-GFP for 1 h. Next, inoculum was removed before the addition of 10 ml of virus growth media consisting of serum-free DMEM, 0.2% w/v bovine serum albumin (BSA; VWR), and 1 μg/ml L-1-tosylamido-2-phenylethyl chloromethyl ketone (TPCK)-treated trypsin (Sigma Aldrich). Cells were incubated for 6 days at 37°C. On the day of harvest, cells were scraped, resuspended in the viral growth media, and centrifuged at 4000 rpm for 10 min. The supernatant was aliquoted and frozen at -80°C until use. For neutralization assays, RSV-A2-GFP or hMPV-GFP was incubated with serial dilutions of heat-inactivated mouse serum in black, clear-bottom 96-well plates (Catalog No. 3603, Corning) in DMEM containing 2.5% FBS or serum-free DMEM containing 10 μg/ml TPCK-treated trypsin, respectively, for 1 h at 37°C. Assays were performed in duplicate with 3-fold dilutions of antibodies (scFv-Fc and IgG) starting at 0.1-10 µg/ml and mouse serum starting at a 1:300 dilution. For RSV-A2-GFP neutralization, after 1 h of virus/serum incubation, 1.5 × 10^4^ VeroE6 cells in DMEM containing 2.5% FBS were then added to each well, after which the plates were incubated for 72 h. For hMPV-GFP neutralization, after virus/serum incubation, 1.5 × 10^4^ VeroE6 cells in DMEM containing 2.5% FBS were added to each well, producing a final concentration of 5 μg/ml TPCK-treated trypsin and 1.25% FBS. The plates were incubated for 72 h at 37°C. On the day of the readout, supernatant was removed from the plates, 100 μl of PBS was added to each well, and the plates were read for GFP on a BioTek Synergy plate reader using Gen5 software. Values from experimental wells were compared against wells containing virus only, with both subtracting background fluorescence from cell control wells. Dose-response neutralization curves were then fitted by nonlinear regression in GraphPad Prism 10.0.2 software, with the IC_50_ and ID_50_ values calculated with constraints set between 0 and 100% neutralization.

### Statistical analysis

Data were collected from 10 mice per group in the immunization studies. All assays, including serum binding and live RSV and hMPV neutralization, were performed in duplicate. Different vaccine constructs, adjuvant-formulated RSV-F and hMPV-F immunogens, were compared using one-way analysis of variation (ANOVA), followed by Tukey’s multiple comparison *post hoc* test. Statistical significance is indicated as the following in the figures: ns (not significant), \**p* < 0.05, \*\**p* < 0.01, \*\*\**p* < 0.001, and \*\*\*\**p* < 0.0001. The graphs were generated, and statistical analyses were performed using GraphPad Prism 10.0.2 software.

## ACKNOWLEDGEMENTS

Diffraction data were collected at the Stanford Synchrotron Radiation Lightsource (SSRL) beamlines 12-1 and 12-2. Use of the SSRL and Stanford Linear Accelerator Center (SLAC) National Accelerator Laboratory is supported by the U.S. Department of Energy, Office of Science, Office of Basic Energy Sciences under Contract No. DE-AC02-76SF00515. The SSRL Structural Molecular Biology Program is supported by the Department of Energy (DOE) Office of Biological and Environmental Research, and by the National Institutes of Health, National Institute of General Medical Sciences (P30GM133894). We thank V. Tong and M. Arends for proofreading the manuscript.

## Funding

This work was supported by Ufovax/SFP-2018-1013 (J.Z.).

## Author contributions

Project design by Y.-Z.L., J.H., Y.-N.Z., K.B.G., L.H., and J.Z.; protein design by J.Z.; protein expression and purification by G.W., Y.-Z.L., and L.H.; EM analysis by Y.-Z.L., L.H., and J.Z.; crystal structures by Y.-Z.L., R.L.S., and I.A.W; mouse immunization, sample collection, and serum binding analysis by Y.-N.Z; live RSV and hMPV propagation and neutralization assay by G.W., K.B.G., and L.H.; manuscript written by Y.-Z.L., J.H., Y.-N.Z., K.B.G., S.A., L.H., and J.Z. All authors were asked to comment on the manuscript. The Scripps Research Institute manuscript No. is 30254.

## Competing interests

The authors declare that they have no competing interests.

## Data and material availability

All data to understand and assess the conclusions of this research are available in the main text and Supplementary Materials.

## SUPPLEMENTAL LEGENDS

**Fig. S1.**
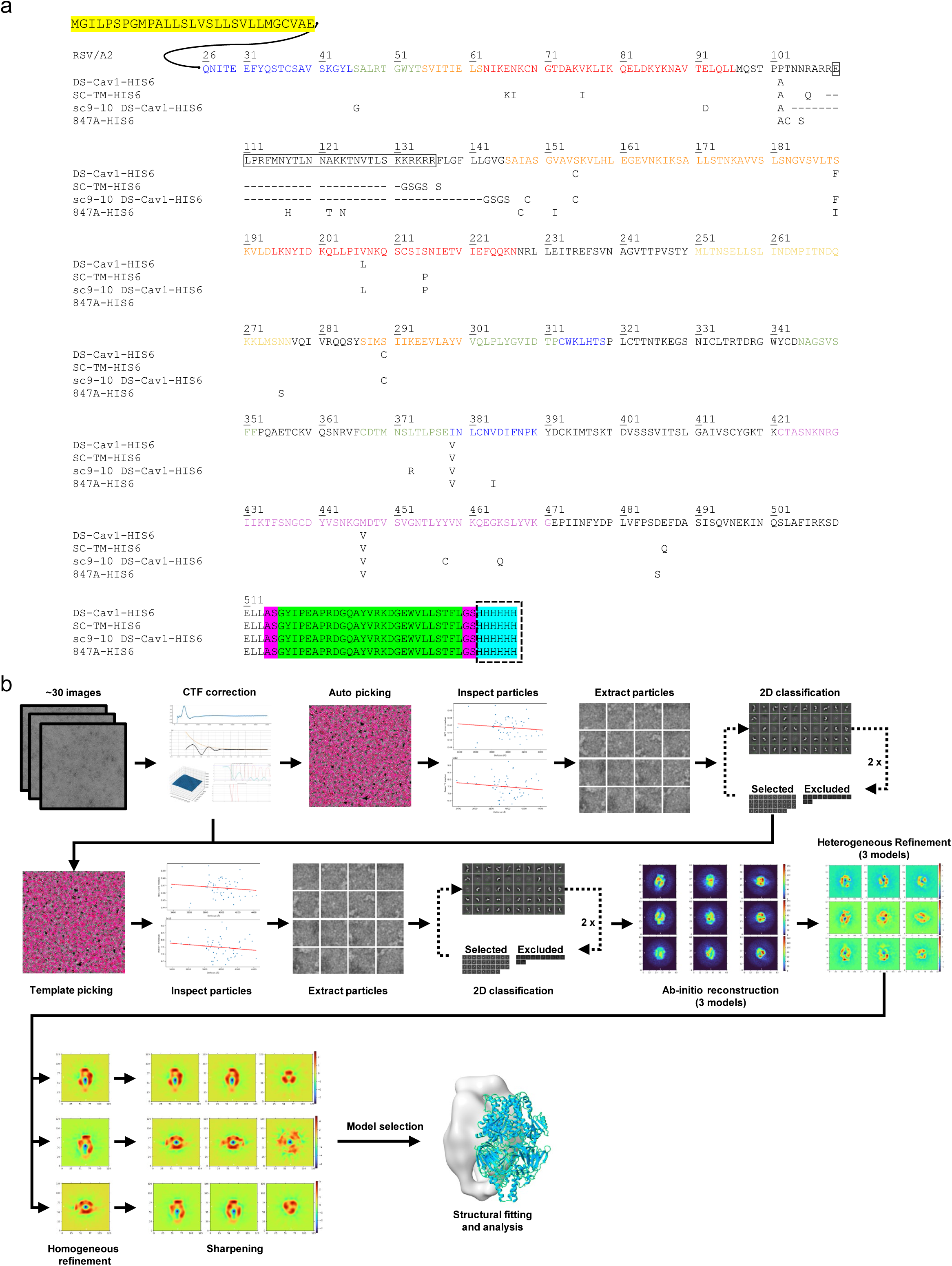
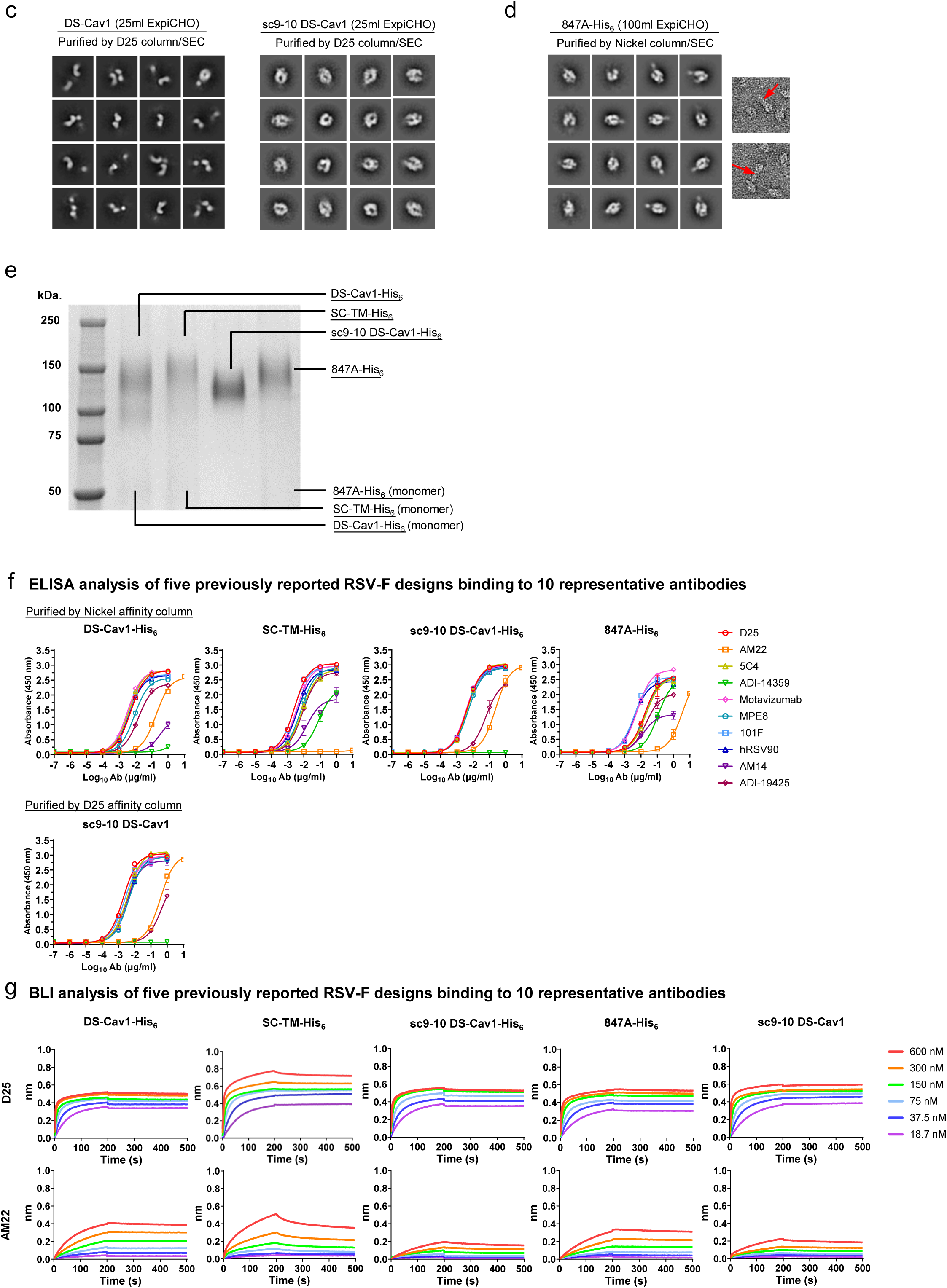
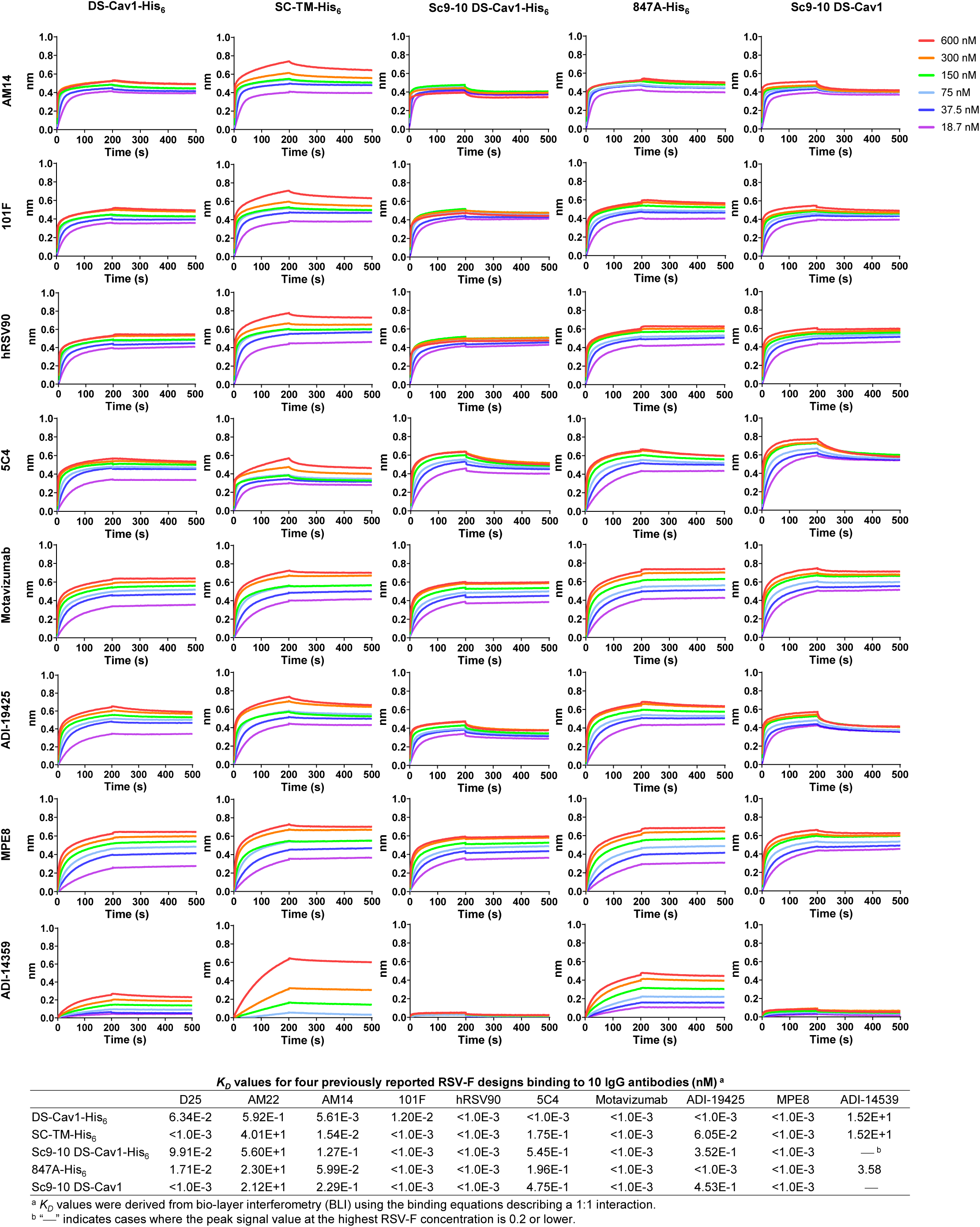
In vitro characterization of previously reported RSV-F designs. (**a**) Sequence alignment of four previously reported RSV-F designs based on the RSV A2 strain backbone. The six major antigenic sites are color-coded as follows: site Ø in red, I in blue, II in yellow, III in green, IV in purple, and V in orange. The p27 peptide is circled in black line boxes. The leader sequence used in the expression constructs is shown atop the sequence alignment and highlighted in yellow shade. The restriction site, foldon trimerization motif, and His_6_ tag are highlighted in pink, green, and cyan shades. The His_6_ tag is included to facilitate immobilized nickel affinity (termed Nickel) purification and is circled in a black dotted line box. (**b**) Schematic representation of image processing, 2D classification, and 3D reconstruction of negative stain EM (nsEM) data obtained for RSV-F samples using CryoSPARC. (**c**) Representative 2D classification images of D25/SEC-purified DS-Cav1 and sc9-10 DS-Cav1. (**d**) Representative 2D classification images of Nickel/SEC-purified 847A-His_6_ from a different production run expressed in 100 ml ExpiCHO cells. (**e**) SDS-PAGE analysis of various RSV-F constructs under reducing conditions. Prior to the analysis, proteins were treated with 5 mM disuccinimidyl glutarate (DSG) for crosslinking F protomers. Each well was loaded with 2 μg of the appropriate antigen. (**f**) ELISA binding curves of various RSV-F constructs with 10 representative antibodies. Briefly, each well were coated with 0.1 μg of the appropriate antigen and IgG antibodies were diluted from a starting concentration of 1 μg/ml with a 10-fold dilution series except for AM22, which started from a concentration of 10 μg/ml. (**g**) BLI analysis of five RSV-F antigens using 10 representative antibodies. Octet binding curves are shown for five RSV-F antigens with 10 antibodies in the IgG form. Sensorgrams were obtained from an Octet RED96 instrument using the AHC biosensor. A two-fold concentration gradient of antigen, starting at 600 nM, was used in a dilution series of six. *K_D_* values were derived from a 1:1 fitting model. In (f) and (g), the four His-tagged RSV-F trimers were purified using a Nickel column, and Sc9-10 DS-Cav1 trimers were purified using a D25 antibody column.

**Fig. S2.**
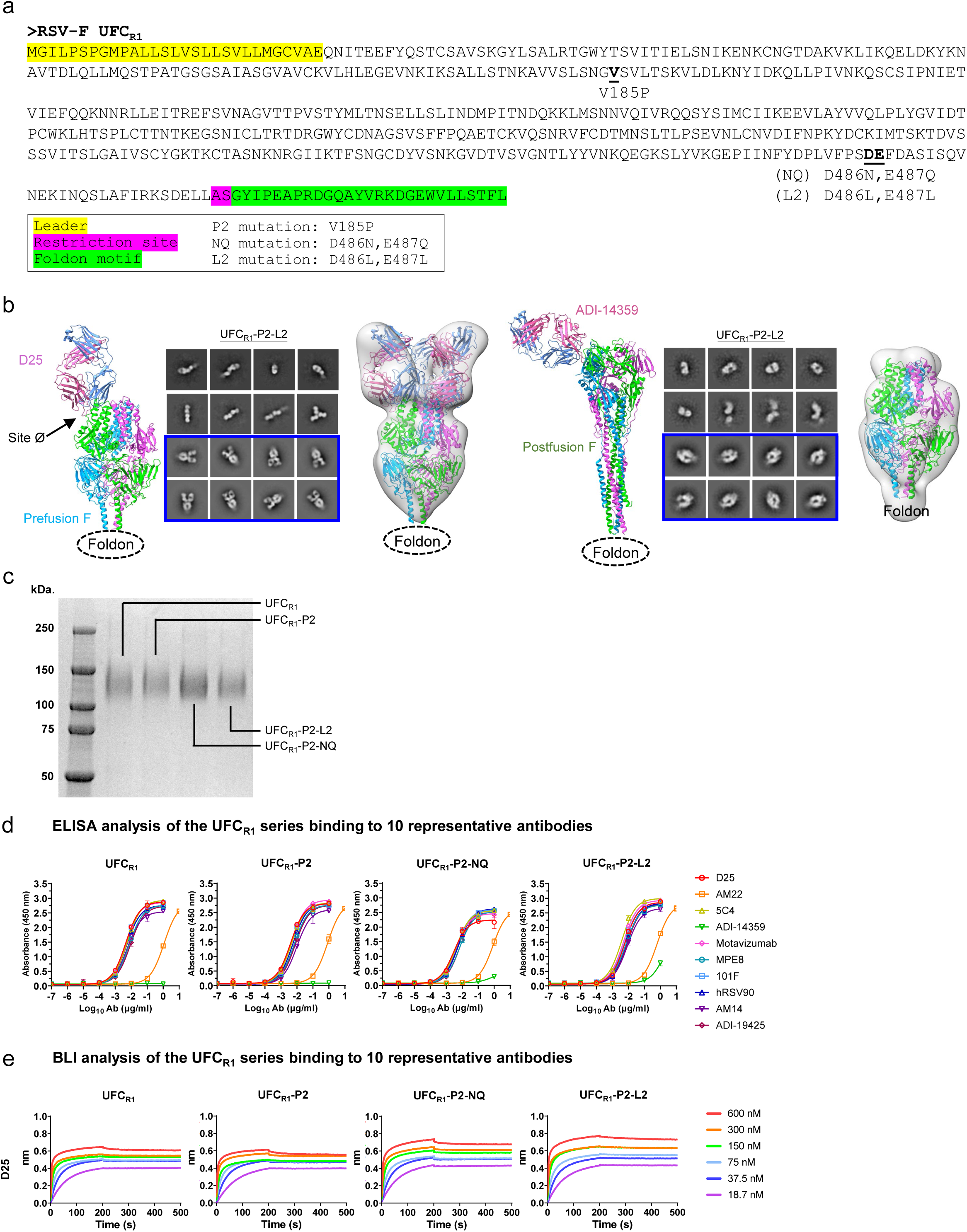
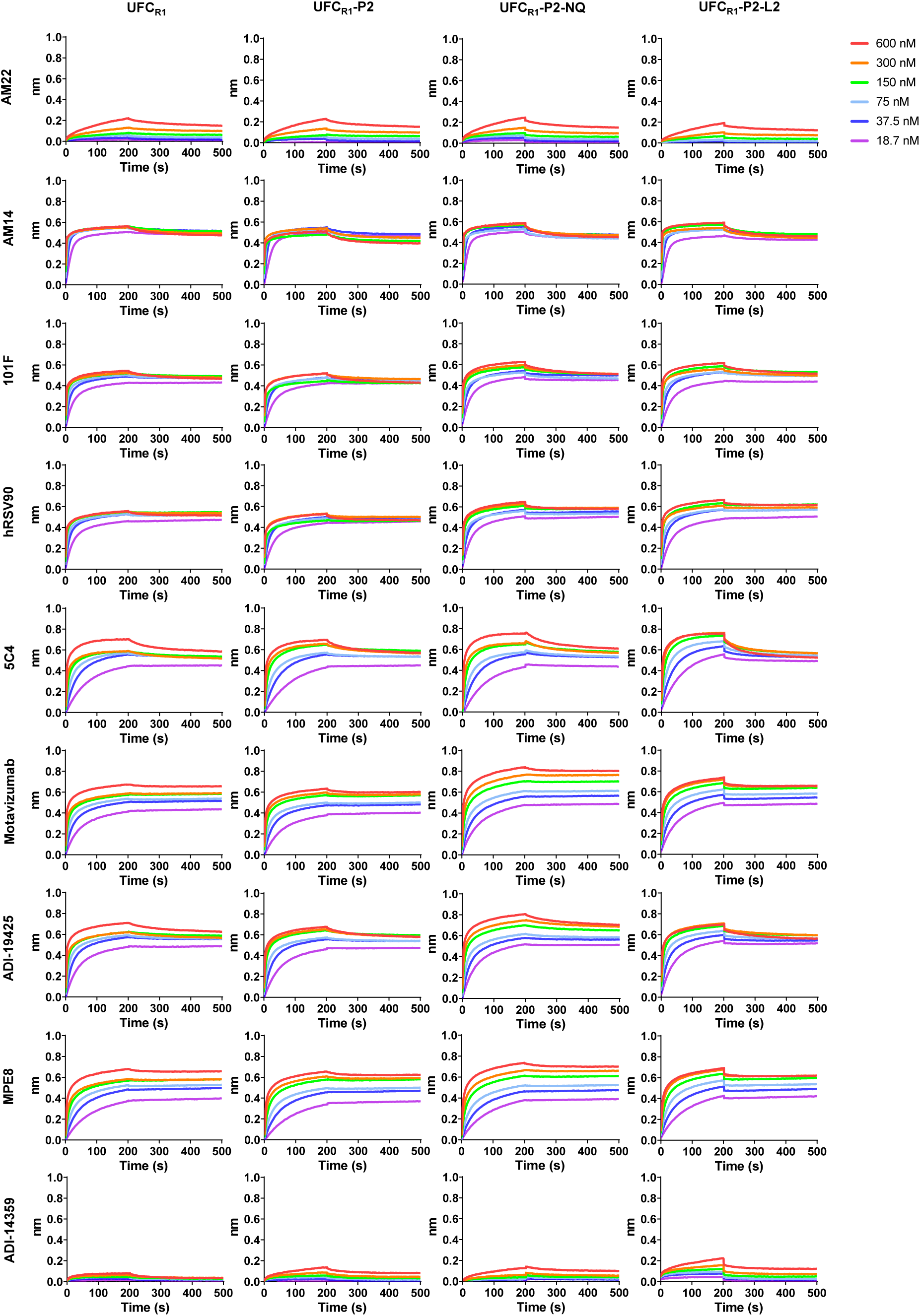
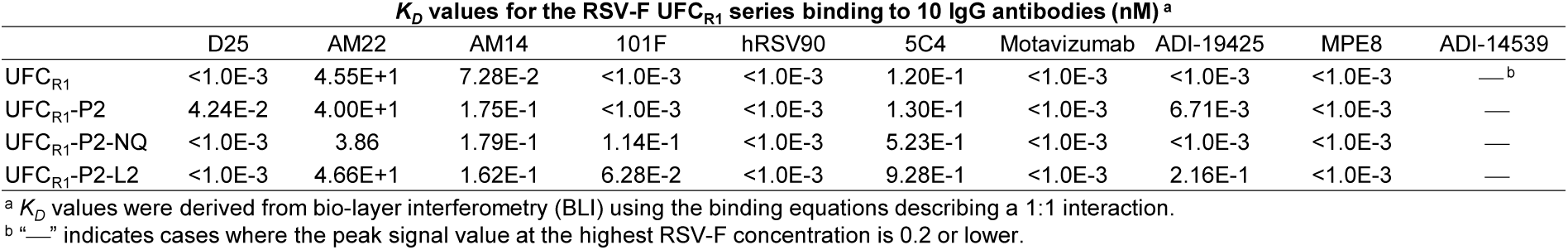
In vitro characterization of the RSV-F UFC_R1_ series. (**a**) Construct design of UFC_R1_, UFC_R1_-P2, UFC_R1_-P2-NQ, and UFC_R1_-P2-L2. The sequence of UFC_R1_ is shown with the P2, NQ, and L2 mutations labeled. The leader sequence, restriction site, and foldon trimerization motif are highlighted in yellow, pink, and green shades. (**b**) The nsEM analysis of UFC_R1_-P2-L2 in the presence of prefusion-specific antibody D25 (left panel) or postfusion-specific antibody ADI-14359 (right panel). Each panel shows the ribbons model of the RSV-F/antibody complex on the left, representative 2D classification images in the middle, and 3D reconstruction on the right. The 2D class images corresponding to prefusion-closed trimers (either ligand-free or antibody-bound) are circled in blue, and a prefusion RSV-F trimer (PDB ID: 4JHW) is used for structural fitting into the nsEM densities. Notably, UFC_R1_-P2-L2/ADI-14359 complexes were not identified in nsEM micrographs and subsequent 2D classification images. (**c**) SDS-PAGE analysis of four UFC_R1_ trimers under reducing conditions. Prior to the analysis, UFC_R1_ trimers were treated with 5 mM disuccinimidyl glutarate (DSG) for crosslinking F protomers. Each well was loaded with 2 ug of the appropriate antigen. (**d**) ELISA binding curves of four UFC_R1_-series constructs with ten representative antibodies. Briefly, each well were coated with 0.1 μg of the appropriate antigen and antibodies were diluted from a starting concentration of 1 μg/ml with a 10-fold dilution series except for AM22, which started from a concentration of 10 μg/ml. (**e**) BLI analysis of four UFC_R1_-series constructs using 10 representative antibodies. Octet binding curves are shown for four UFC_R1_-series constructs with 10 antibodies in the IgG form. Sensorgrams were obtained from an Octet RED96 instrument using the AHC biosensor. A two-fold concentration gradient of antigen, starting at 600 nM, was used in a dilution series of six. *K_D_* values were derived from a 1:1 fitting model.

**Fig. S3.**
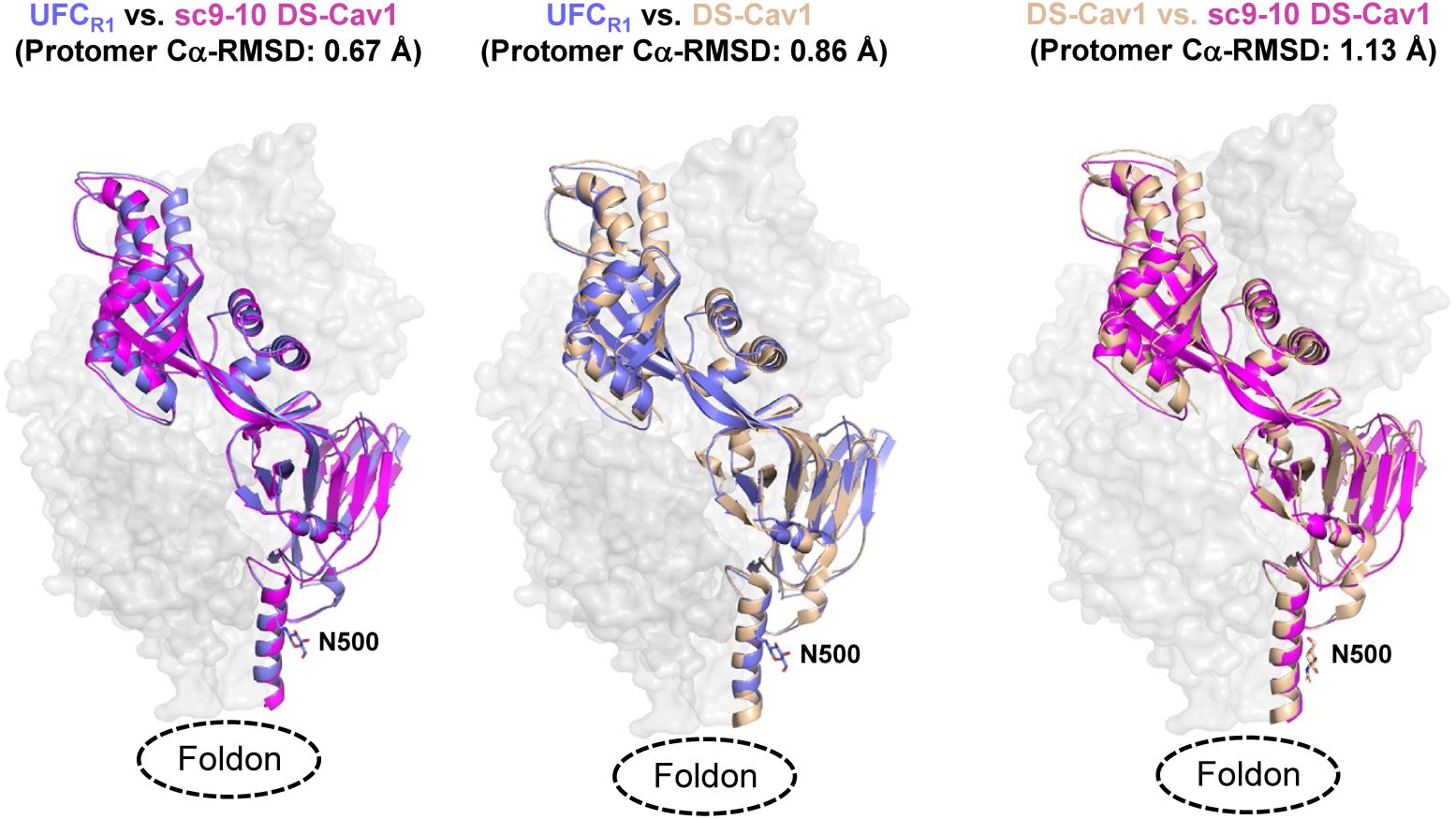
Additional structural analysis of the RSV-F UFC_R1_ series. Structural comparison between UFC_R1_ and the two DS-Cav1 design variants. Left: Crystal structures of UFC_R1_ and sc9-10 DS-Cav1 are superimposed and shown as blue and pink ribbons models, respectively, within the gray trimer surface. The Cα-RMSD between UFC_R1_ and sc9-10 DS-Cav1 is 0.67 Å. Middle: Crystal structures of UFC_R1_ and DS-Cav1 superimposed and shown as blue and gold ribbons models, respectively, within the gray trimer surface. The Cα-RMSD between UFC_R1_ and DS-Cav1 is 0.86 Å. Right: Crystal structures of sc9-10 DS-Cav1 and DS-Cav1 superimposed and shown as pink and gold ribbons models, respectively, within the gray trimer surface. The Cα-RMSD between sc9-10 DS-Cav1 and DS-Cav1 is 1.13 Å.

**Fig. S4.**
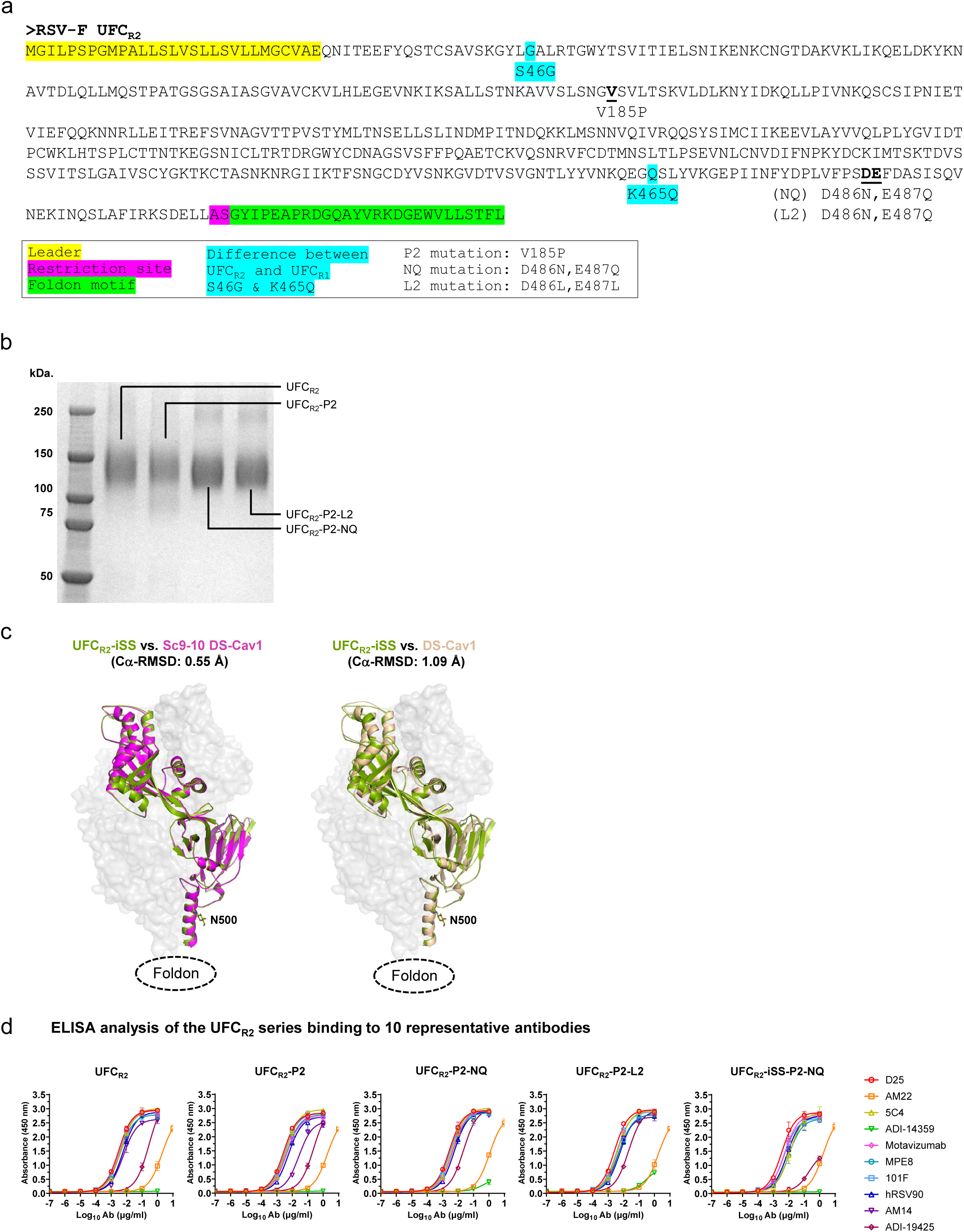
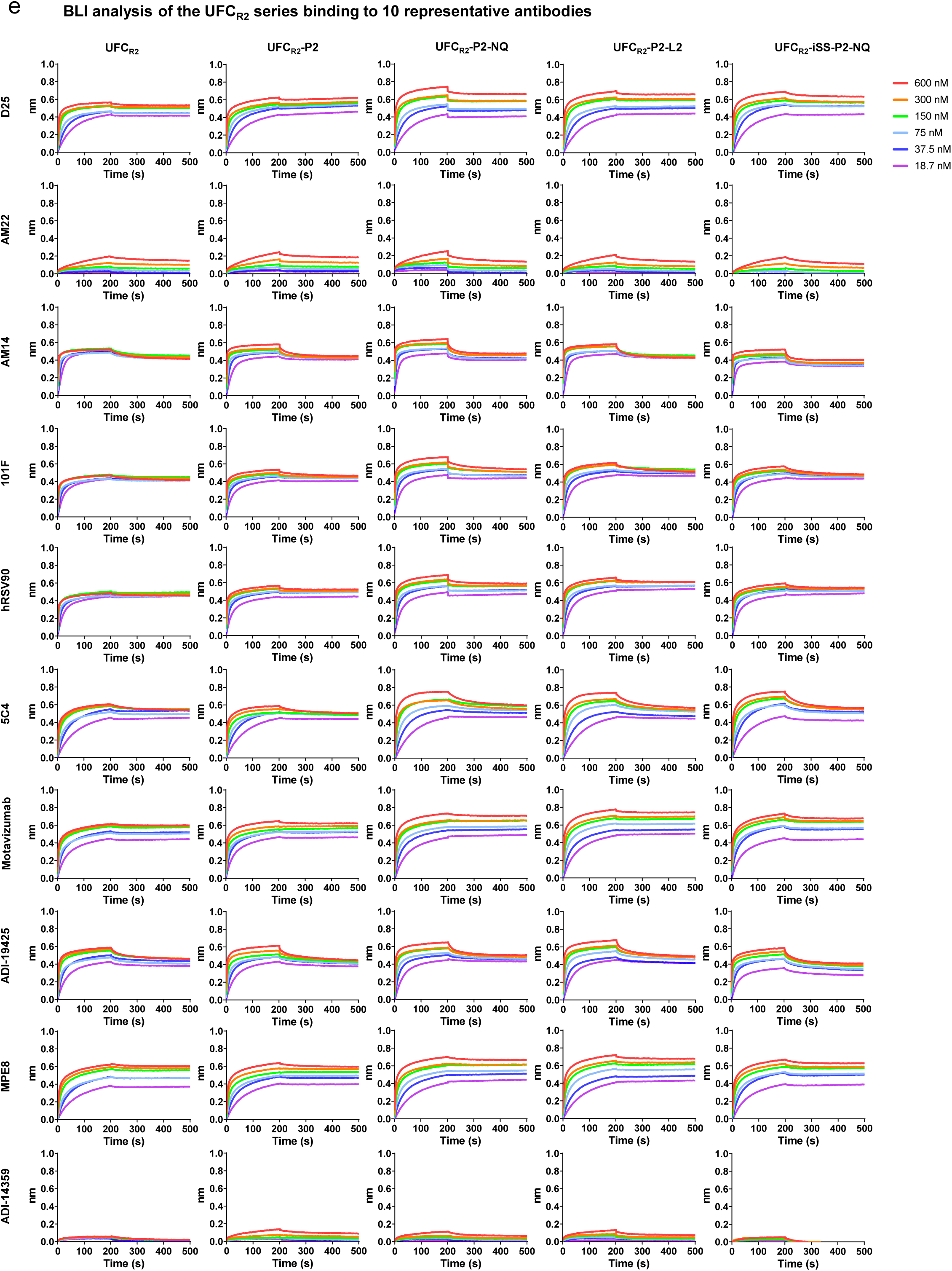
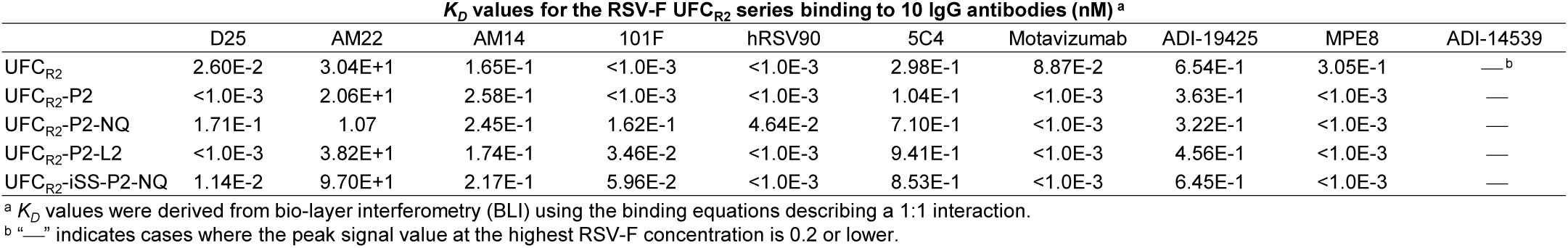
In vitro characterization and additional structural analysis of the RSV-F UFC_R2_ series. (**a**) Construct design of UFC_R2_, UFC_R2_-P2, UFC_R2_-P2-NQ, and UFC_R2_-P2-L2. The sequence of UFC_R2_ is shown with the S46G/K465Q, P2, NQ, and L2 mutations labeled. The leader sequence, restriction site, and foldon trimerization motif are highlighted in yellow, pink, and green shades. (**b**) SDS-PAGE analysis of four UFC_R2_ trimers under reducing conditions. Prior to the analysis, UFC_R2_ trimers were treated with 5 mM disuccinimidyl glutarate (DSG) for crosslinking F protomers. Each well was loaded with 2 ug of the appropriate antigen. (**c**) Structural comparison between UFC_R2_-iSS and the two DS-Cav1 variants. Left: Crystal structures of UFC_R2_-iSS and sc9-10 DS-Cav1 are superimposed and shown as green and pink ribbons models, respectively, within the gray trimer surface. The Cα-RMSD between UFC_R2_-iSS and sc9-10 DS-Cav1 is 0.55 Å. Right: Crystal structures of UFC_R2_-iSS and DS-Cav1 superimposed and shown as green and gold ribbons models, respectively, within the gray trimer surface. The Cα-RMSD between UFC_R2_-iSS and DS-Cav1 is 1.09 Å. (**d**) ELISA binding curves of five UFC_R2_–series constructs with ten representative antibodies. Briefly, each well were coated with 0.1 μg of the appropriate antigen and antibodies were diluted from a starting concentration of 1 μg/ml with a 10-fold dilution series except for AM22, which started from a concentration of 10 μg/ml. (**e**) BLI analysis of five UFC_R2_-series constructs using 10 representative antibodies. Octet binding curves are shown for five UFC_R2_–series constructs with 10 IgG antibodies in the IgG form. Sensorgrams were obtained from an Octet RED96 instrument using the AHC biosensor. A two-fold concentration gradient of antigen, starting at 600 nM, was used in a dilution series of six. *K_D_* values were derived from a 1:1 fitting model. UFC_R2_-iSS-P2-NQ was included in both (**d**) and (**e**) as the last (fifth) construct in antigenic evaluation.

**Fig. S5.**
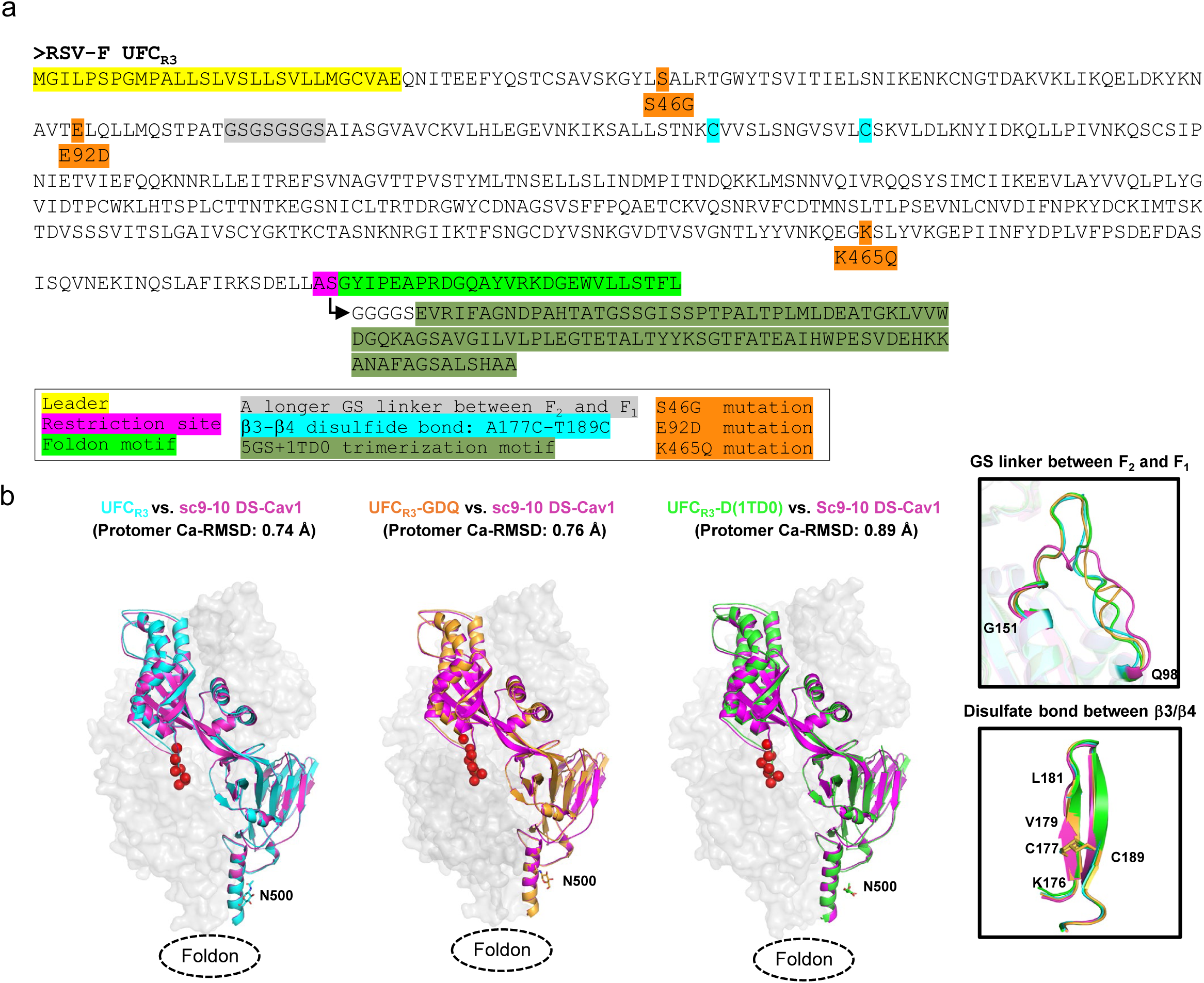
In vitro characterization and additional structural analysis of the RSV-F UFC_R3_ series. (**a**) Construct design of UFC_R3_, UFC_R3_-GDQ, UFC_R3_-D(1TD0). The sequence of UFC_R3_ is shown with the leader sequence, restriction site, and foldon trimerization motif are highlighted in yellow, pink, and green shades. The longer F_2_-F_1_ GS linker and the A177C-T89C disulfide bond between β3 and β4 that are unique to UFC_R3_ are highlighted in gray and cyan shades, respectively. The S46G. E92D, and K465Q mutations in UFC_R3_-GDQ are highlighted in orange shade, and the 1TD0 trimerization motif unique to UFC_R3_-D(1TD0) is highlighted in dark green shade. (**b**) Structural comparison of the UFC_R3_-series with respect to sc9-10 DS-Cav1. The sc9-10 DS-Cav1 (pink)-superimposed UFC_R3_, UFC_R3_-GDQ, and UFC_R3_-D(1TD0) crystal structures are shown as cyan, gold, and green ribbons models within the gray trimer surface. The small panel on the right shows the structural details of the extended F_2_-F_1_ linker (top) and the interstrand disulfide bond A177C-T189C (bottom).

**Fig. S6.**
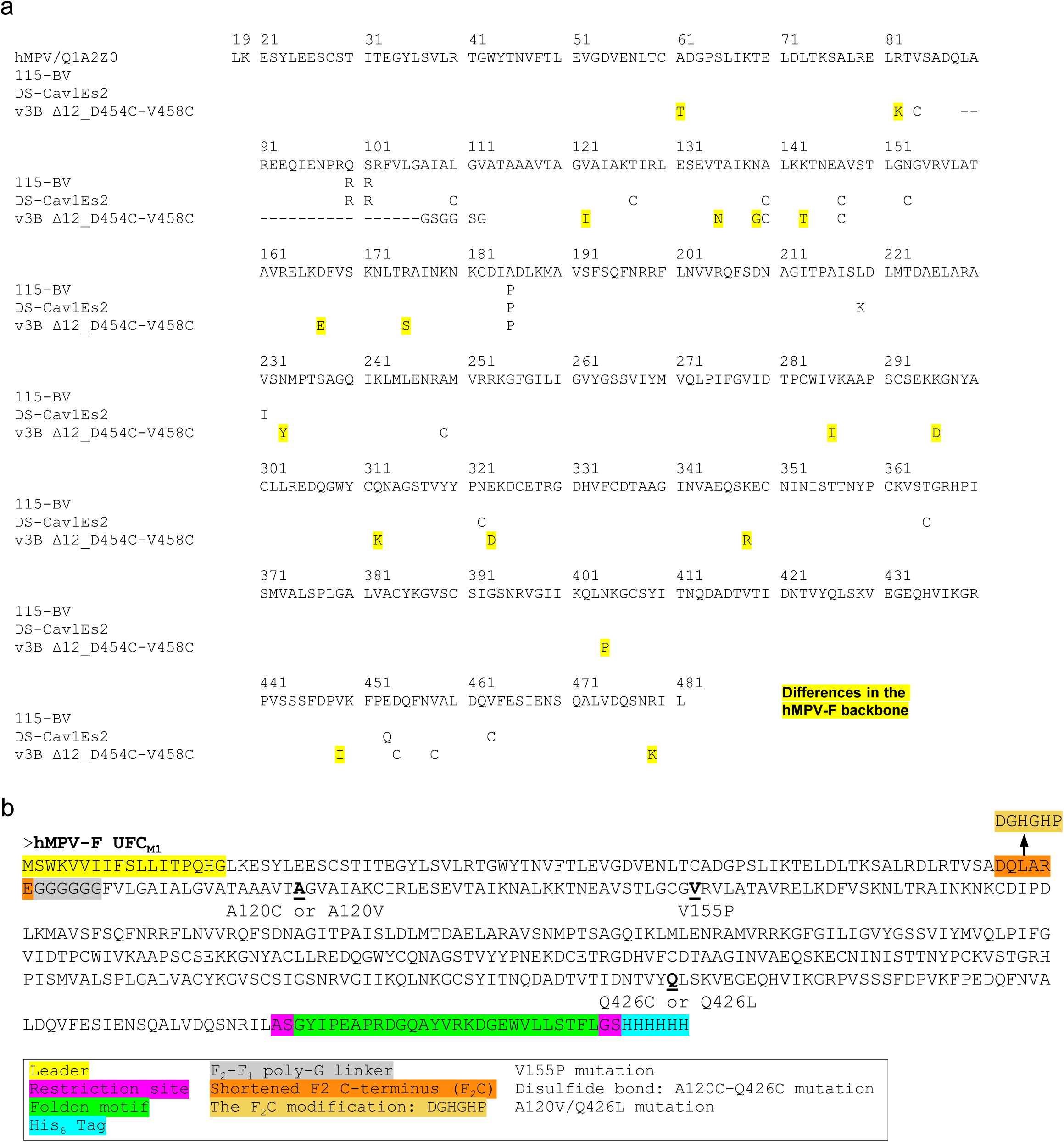
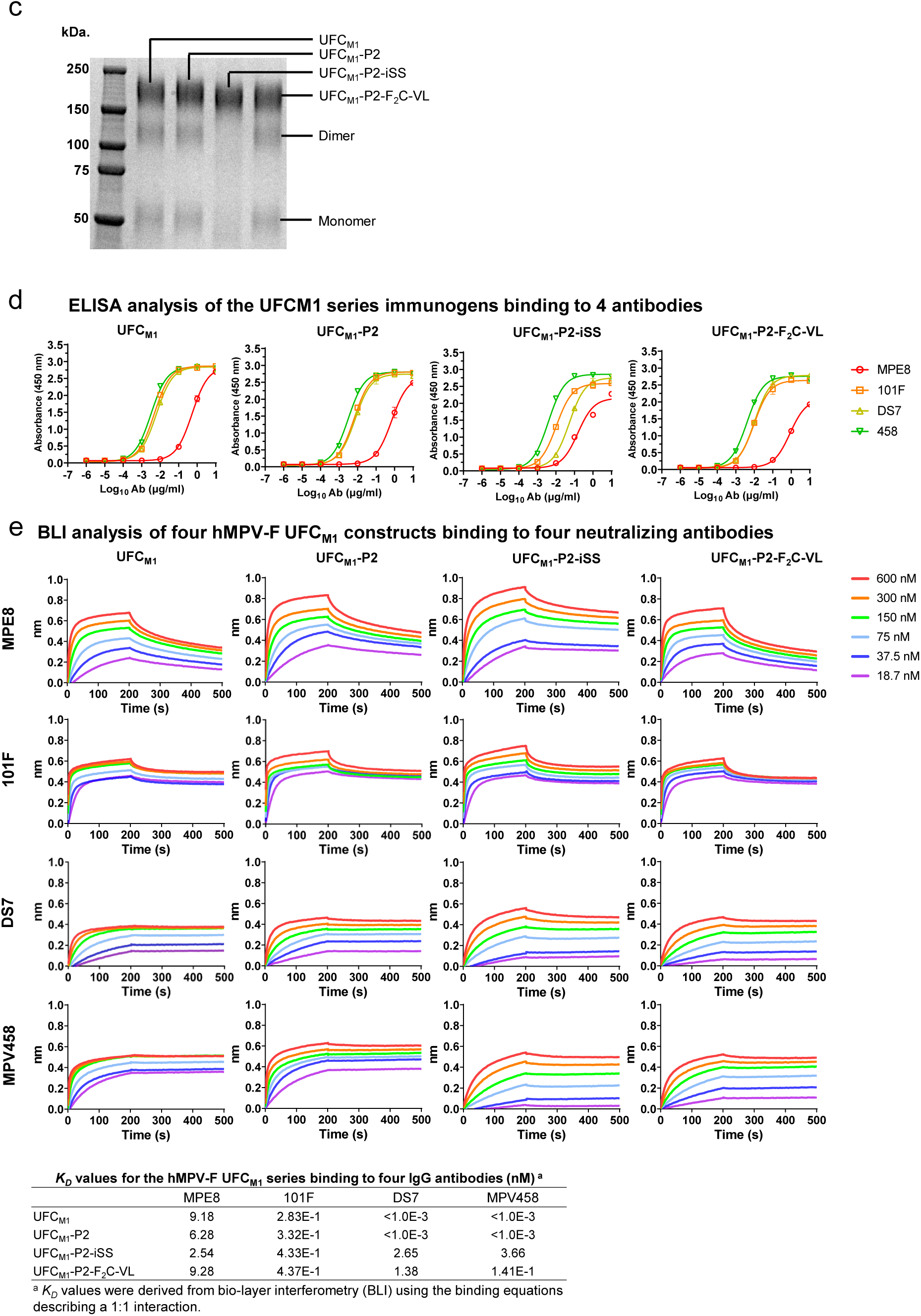
In vitro characterization of the hMPV-F UFC_M1_ series. (**a**) Sequence alignment of three previously reported hMPV-F designs. (**b**) Construct design of UFC_M1_, UFC_M1_-P2, UFC_M1_-P2-iSS, and UFC_M1_-P2-F_2_C-VL, The sequence of UFC_M1_ is shown with the leader sequence, restriction site, foldon trimerization motif, and His_6_ tag are highlighted in yellow, pink, green, and cyan shades, respectively. The V155P mutation in UFC_M1_-P2, A120C/Q426C mutation in UFC_M1_-P2-iSS, and A120V/Q426L (VL) mutation in UFC_M1_-P2-F_2_C-VL are listed. The shortened F2 C-terminus (F_2_c) and its modification in UFC_M1_-P2-F_2_C-VL are highlighted in orange and light orange shades, respectively. The linker (G6) between F2 and F1 is highlighted in gray shade. (**c**) SDS-PAGE analysis of four UFC_M1_ trimers under reducing conditions. Prior to the analysis, UFC_M1_ trimers were treated with 5 mM disuccinimidyl glutarate (DSG) for crosslinking F protomers. Each well was loaded with 2 ug of the appropriate antigen. (**d**) ELISA binding curves of four UFC_M1_-series constructs with four NAbs. Briefly, each well were coated with 0.1 μg of the appropriate antigen and antibodies were diluted from a starting concentration of 1 μg/ml with a 10-fold dilution series. (**e**) BLI analysis of four UFC_M1_–series constructs using four NAbs. Octet binding curves are shown for four UFC_M1_-series constructs with four NAbs in the IgG form. Sensorgrams were obtained from an Octet RED96 instrument using the AHC biosensor. A two-fold concentration gradient of antigen, starting at 600 nM, was used in a dilution series of six. *K_on_* and *K_D_* values were derived from a 1:1 fitting model.

**Fig. S7.**
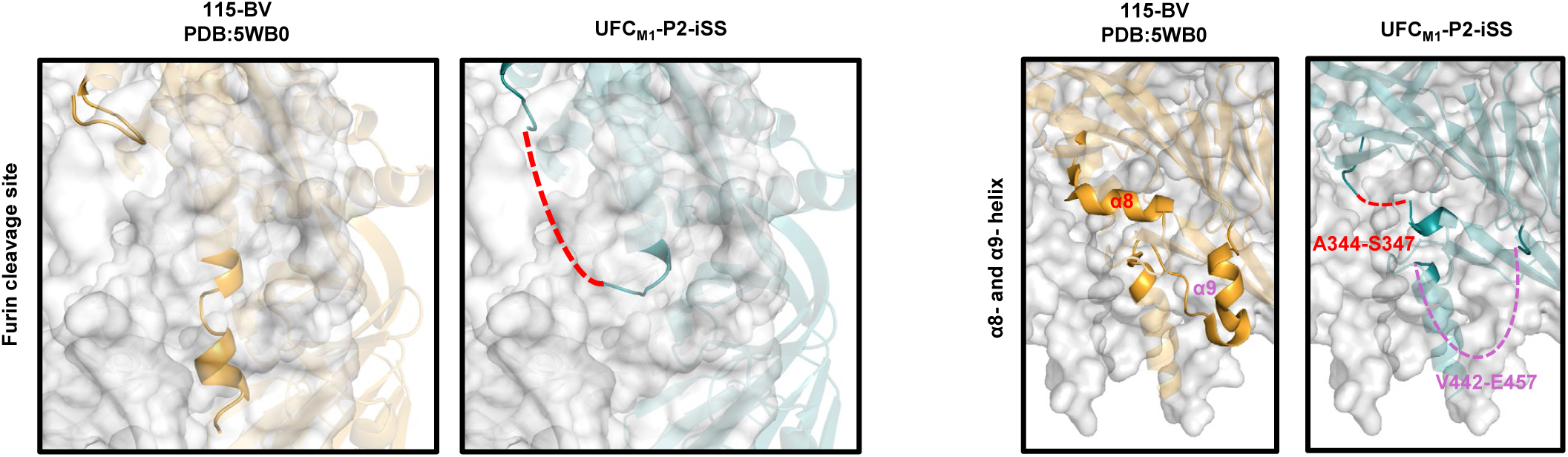
Additional structural analysis of the hMPV-F UFC_M1_-P2-iSS trimer. Close-up view of regions surrounding the F_2_-F_1_ linkage (left), part of α8 (middle, A344-S347), and part of α9-α10-β23 (right, V442-E457) in the 6 Å-resolution crystal structure of UFC_M1_-P2-iSS, which is shown as green ribbons models in the gray trimer surface. A red dotted line is added to indicate where the F_2_-F_1_ linker may be within the closed trimer structure. For the F_2_-F_1_ linkage, the structure of115-VB (PDB ID:5WB0) is shown on the leftmost for comparison, as the cleavage site in the 115-BV sequence allows the expressed F structure to be cleaved between F_2_ and F_1_. The crystal structure of 115-VB is shown as gold ribbons models in the gray trimer surface.

**Fig. S8.**
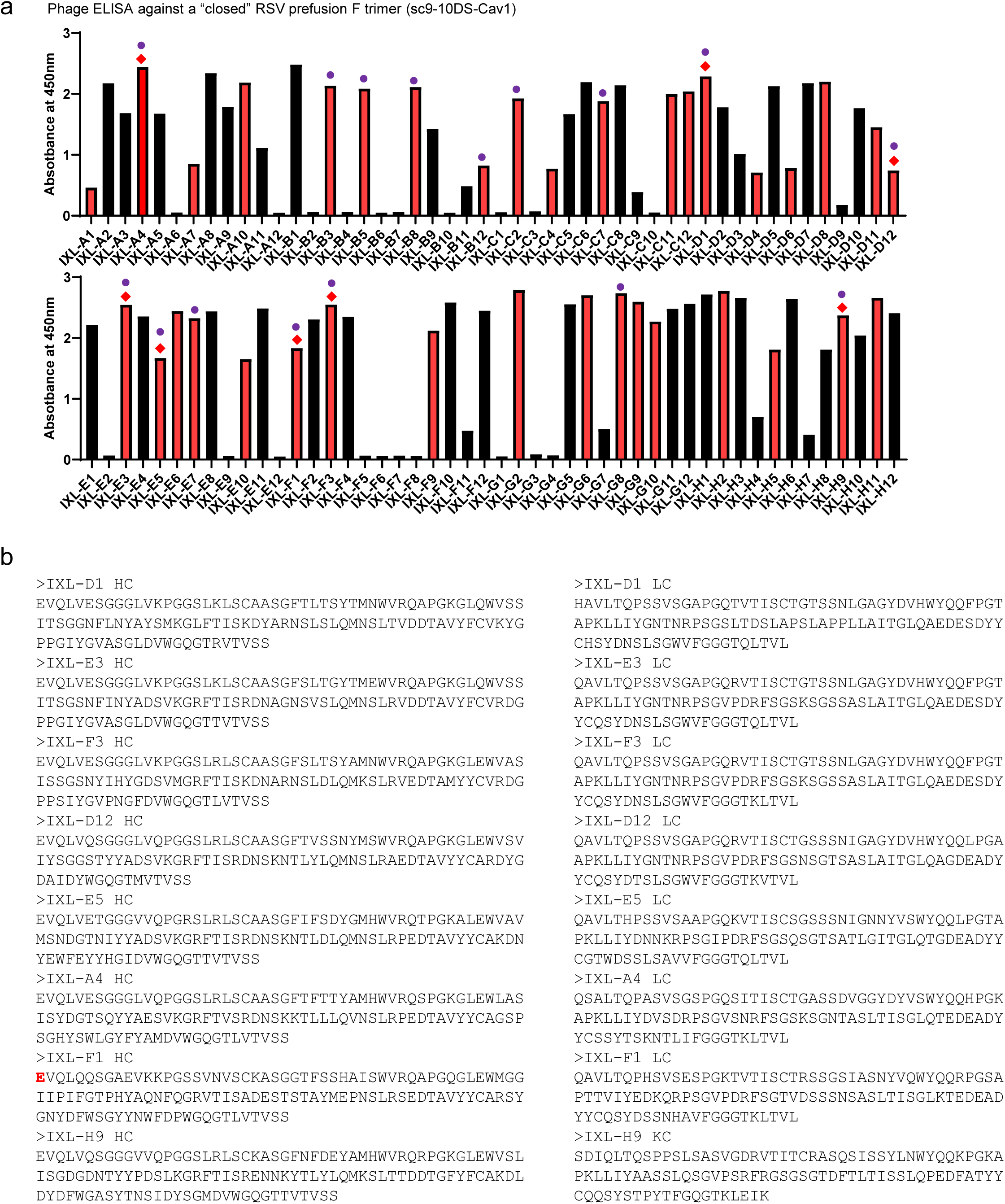
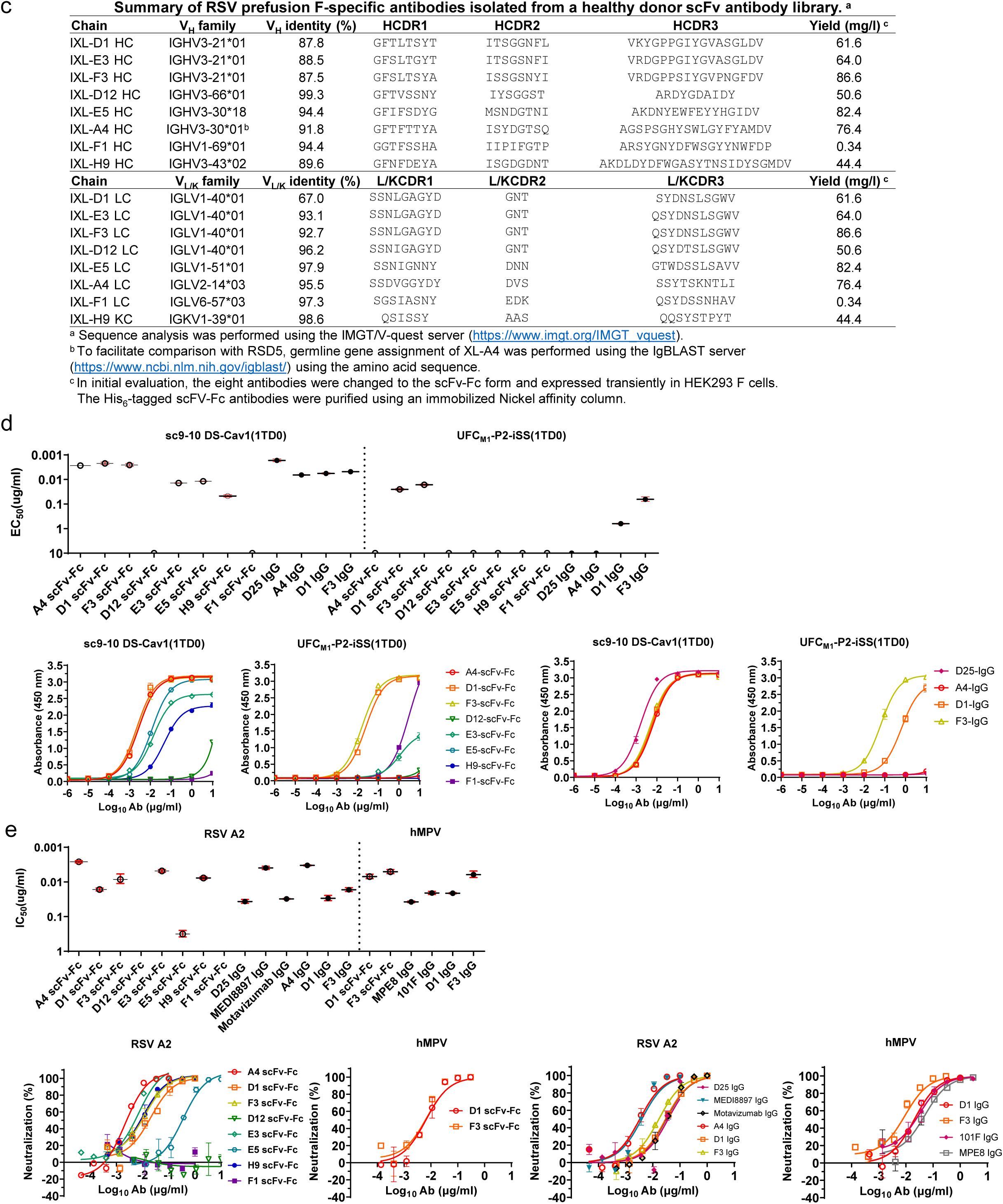
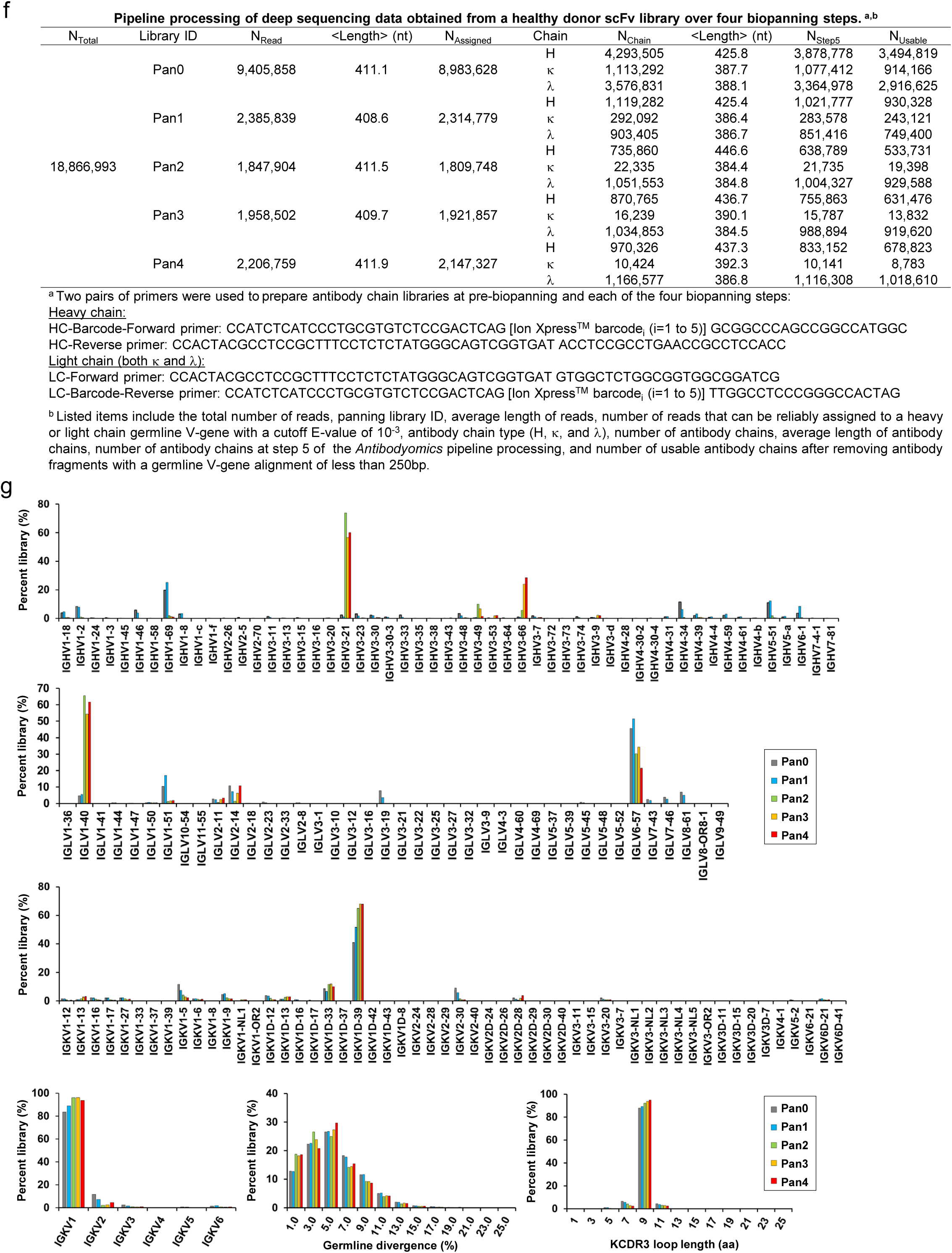
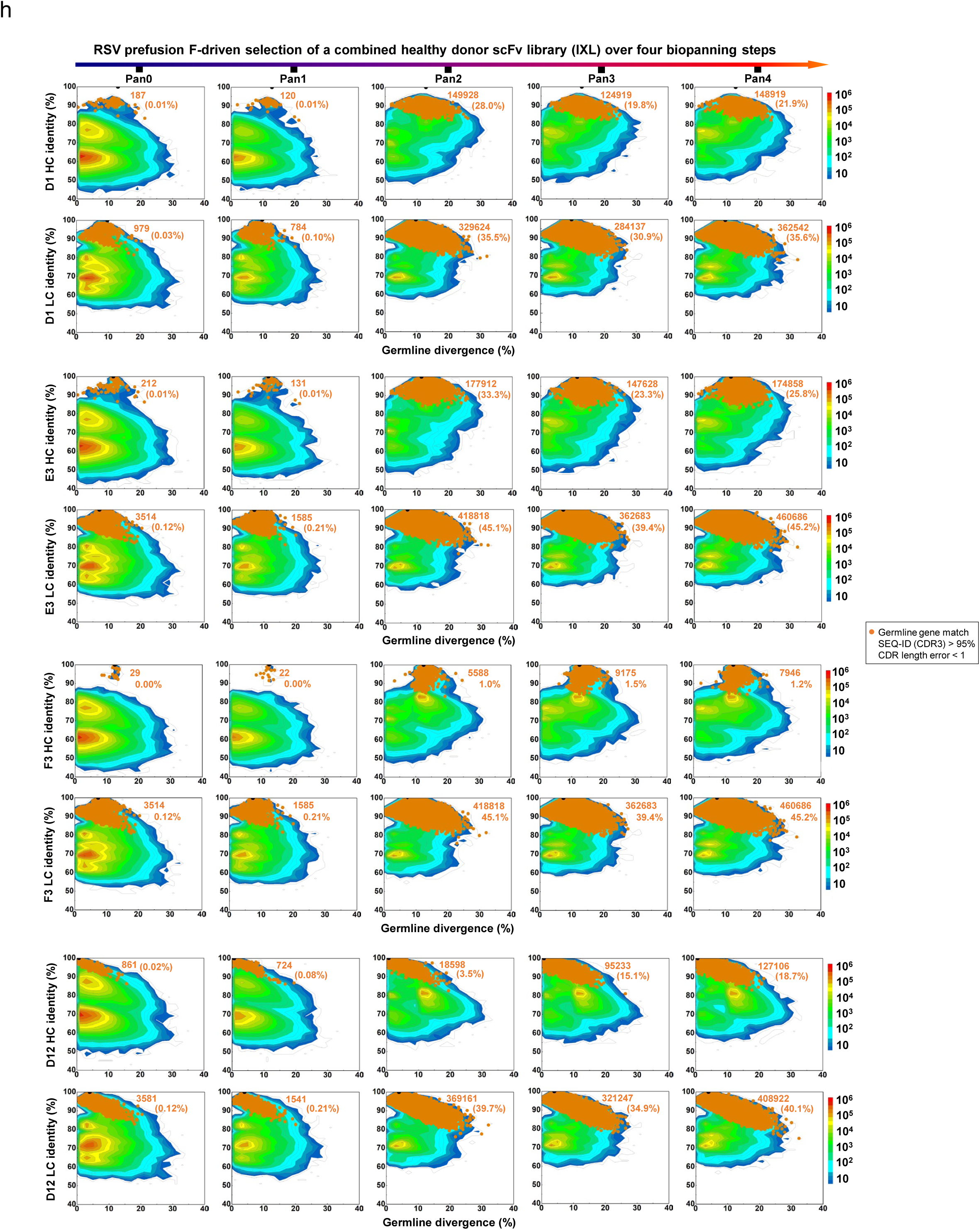
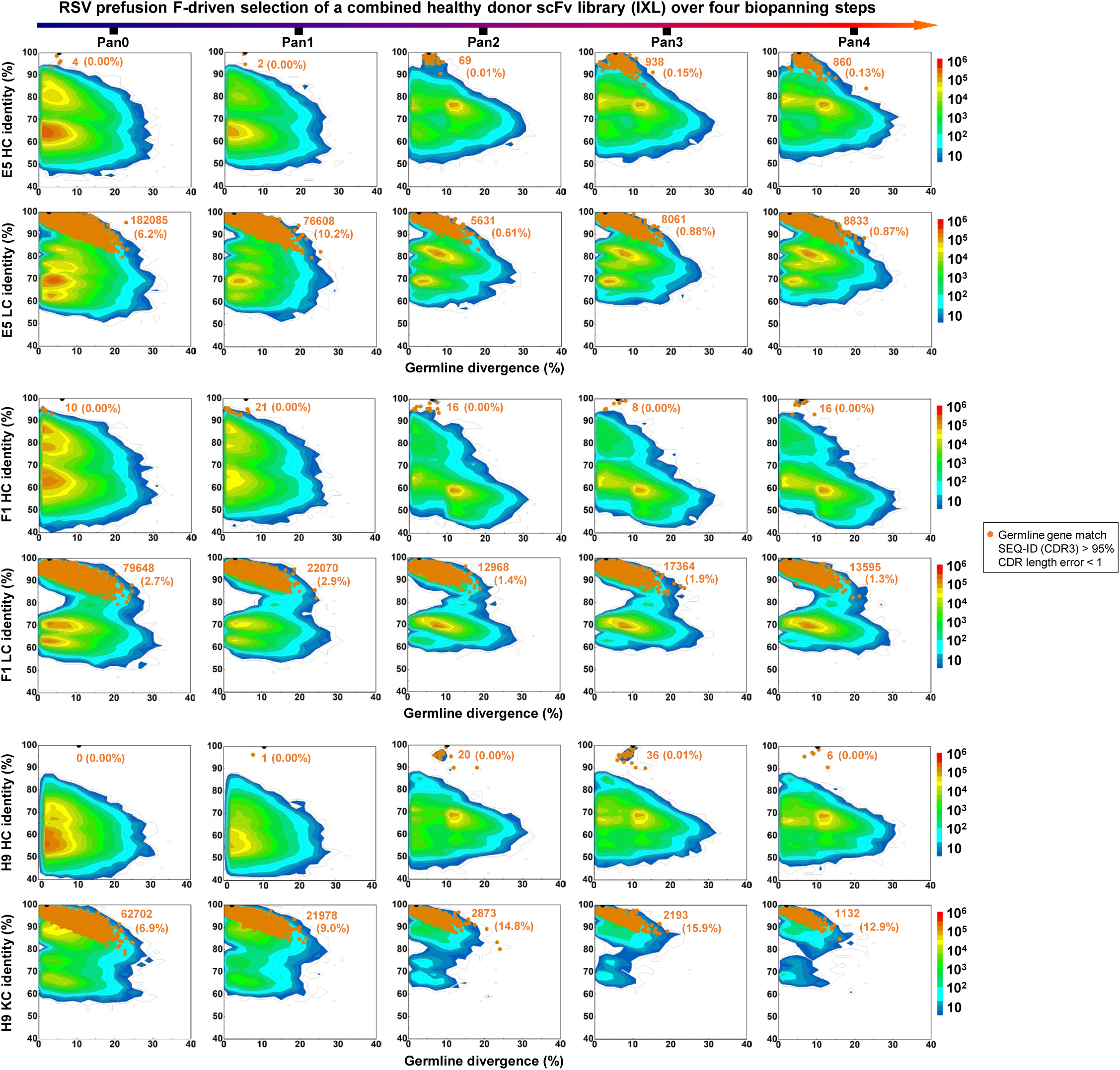
Biopanning and deep sequencing of a phage-displayed human antibody library. Biopanning of a phage scFv library prepared from PBMCs of ten health adult individuals using the RSF UFC_R1_-P2-NQ trimer as a probe was performed to isolate RSV (and potentially hMPV) neutralizing antibodies. (**a**) Phage ELISA was performed to assess randomly picked phage clones for their ability to bind an RSV prefusion F trimer, sc9-10 DS-Cav1, which has a disulfide bond-locked “closed” conformation. The 36 scFv clones that were sequenced are shown in red bars. Of these, 16 scFv clones with complete variable sequences are labeled with solid purple circles and 8 representative scFv clones with red diamond symbols. (**b**) Amino acid sequences of heavy and light chains from 8 representative scFv clones that have complete variable regions from sequencing. A sequencing error was manually corrected for the first amino acid of IXL-F1 HC, which is shown in red. (**c**) Gene family analysis of the phage library-derived antibodies and their yield in transient expression of HEK293 F cells. Key antibody features are summarized in the table. Notably, the scFv gene was cloned into an Fc vector for expression, and as such, the antibodies tested in the initial characterization were in the scFv-Fc form. (**d**) ELISA binding of eight phage library-derived antibodies, in scFv-Fc and some in IgG forms, tested against RSV-F (sc9-10 DS-Cav1) and hMPV-F (UFC_M1_-P2-iSS) antigens. The EC_50_ values are plotted to facilitate comparison between different clones, while the binding curves are shown below the EC_50_ plot. (**f**) Neutralization of eight phage library-derived antibodies, in scFv-Fc and some in IgG forms, tested against live RSV-A2-GFP and hMPV-GFP viruses. The ID_50_ values are plotted to facilitate comparison between different clones, while the neutralization curves are shown below. For (**f**)-(**h**), A total of five libraries, including the pre-panning library and the library after each of the four biopanning steps, were processed into heavy (HC), kappa (KC), and lambda chain (LC) libraries, which were deep sequenced on an Ion GeneStudio S5 sequencer for detailed antibodyomics analysis. (**f**) Pipeline processing of deep sequencing data obtained from the five scFv libraries. (**g**) Quantitative library profiles, including full germline gene usage profiles for the scFv libraries before and after each of the four panning steps (top three panels) and kappa chain-specific profiles for the scFv libraries before and after each of the four panning steps, including germline gene usage (gene families shown), germline divergence (or somatic hypermutation, SHM), and KCDR3 loop length (bottom panel). (**h**) Divergence-identity analysis of seven phage library-derived antibodies (IXL-D1, -E3, -F3, -D12, -E5, -F1, and -H9) within the scFv library before and after each of the four panning steps. To simplify the labeling, the prefix in the clone name “IXL” is not shown for any of the 2D plots. HC and LC/KC sequences are plotted as a function of sequence identity to a given template antibody and sequence divergence from putative germline V genes. Color coding indicates sequence density. On the 2D plots, reference antibodies are shown as black dots, whereas NGS-derived somatic variants that were identified based on the germline V gene match, CDR3 length (with ≤ 1-residue error of margin), CDR3 identity of 95% or greater are shown as orange dots, with the number of sequences and percentage within the germline gene family labeled accordingly.

**Fig. S9.**
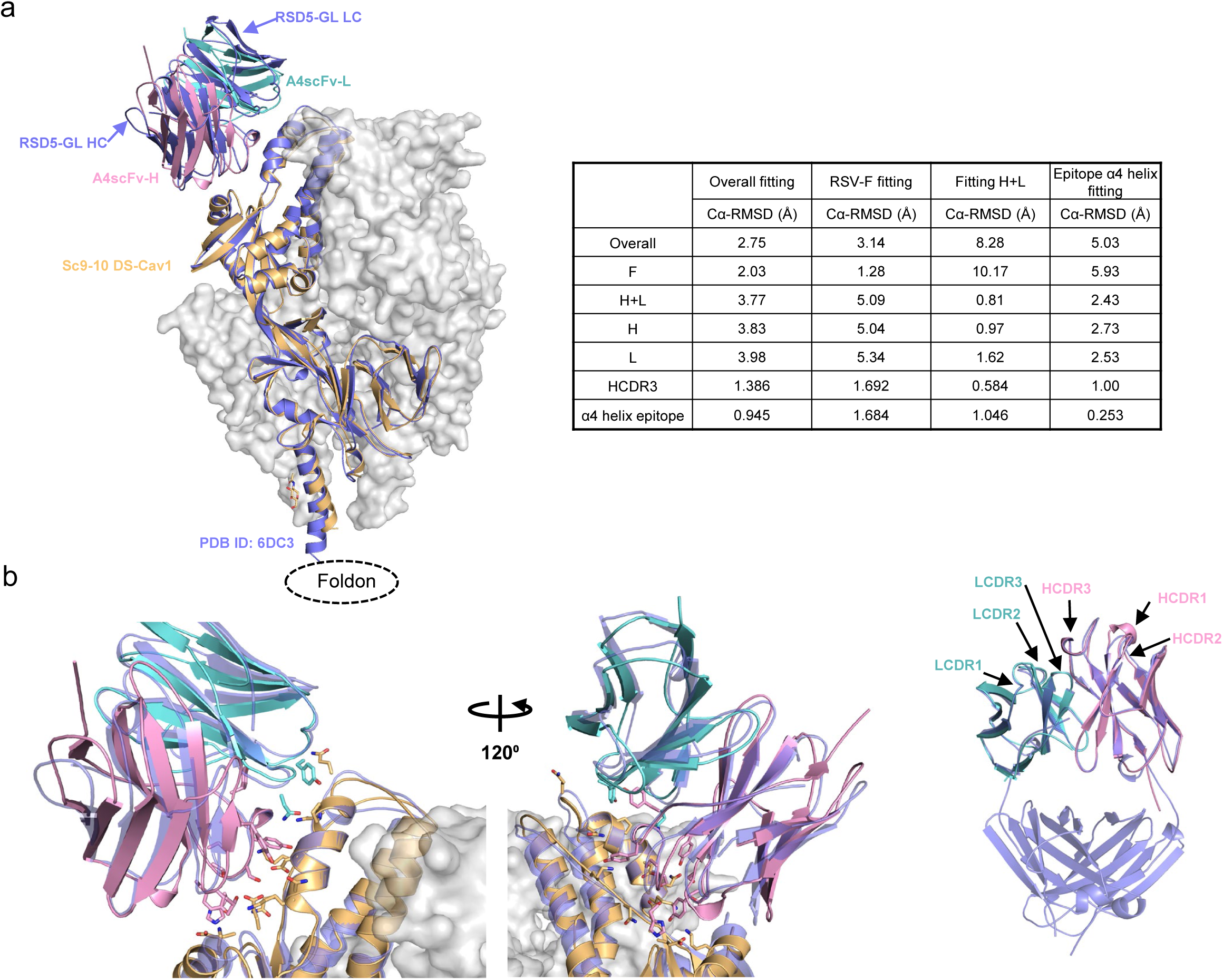
Additional structural analysis of sc9-10 DS-Cav1 in complex with A4 scFv. (**a**) Left: Superimposed structures of sc9-10 DS-Cav1/A4-scFv and DS-Cav1/RSD5-GL (PDB ID: 6DC3) complexes using Cα atoms of both RSV-F and antibody variable regions for fitting. The sc9-10 DS-Cav1/A4-scFv structure is shown as ribbons models with RSV-F colored in gold and A4 scFv heavy and light chains in pink and cyan, respectively, while the DS-Cav1/RSD5-GL Fab complex is shown as blue ribbons models. The overall Cα-RMSD is 2.75 Å at the monomer level. Right: Summary of Cα-RMSD values calculated using different fitting schemes, including overall (RSV-F + antibody HC/LC), RSV-F, HC, LC, HCDR3 (paratope), and α4 helix (epitope). RSD5-GL residues S100-V102 were used for HCDR3 fitting, whereas residues L195-L207 in RSV-F were used for α4 helix fitting (**b**) Left: Closed-up view of the RSV-F/antibody interface for A4 and RSD5-GL after fitting the Cα atoms of the α4 helix in RSV-F. Crystal structures are shown as ribbons models with the same color-coding scheme as in (a). Side chains of key interface residues are shown as stick models. Right: Superimposed A4-scFv and RSD5-GL Fab (PDB ID:6DC3) structures, with the HCDR1-3 and LCDR1-3 loops labeled on the top. Crystal structures are shown as ribbons models with the same color-coding scheme as in (a). The Cα-RMSD for antibody variable regions is 0.81 Å, and the two HCDR3 loops (S100-V102 in RSD5-GL and H100-V102 in A4) have almost identical conformations. All structure alignments and Cα-RMSD calculations were done in Chimera X.

**Fig. S10.**
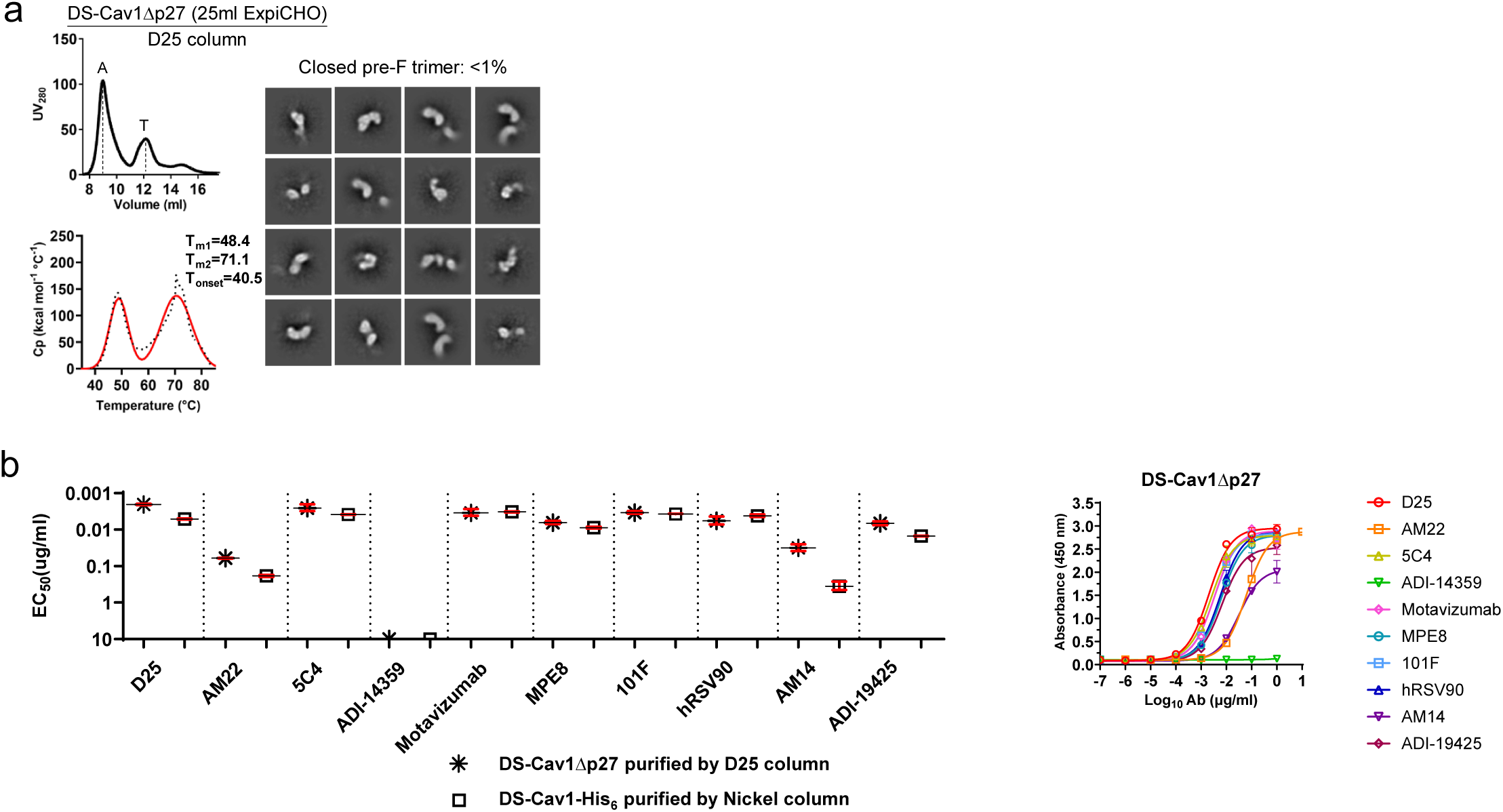
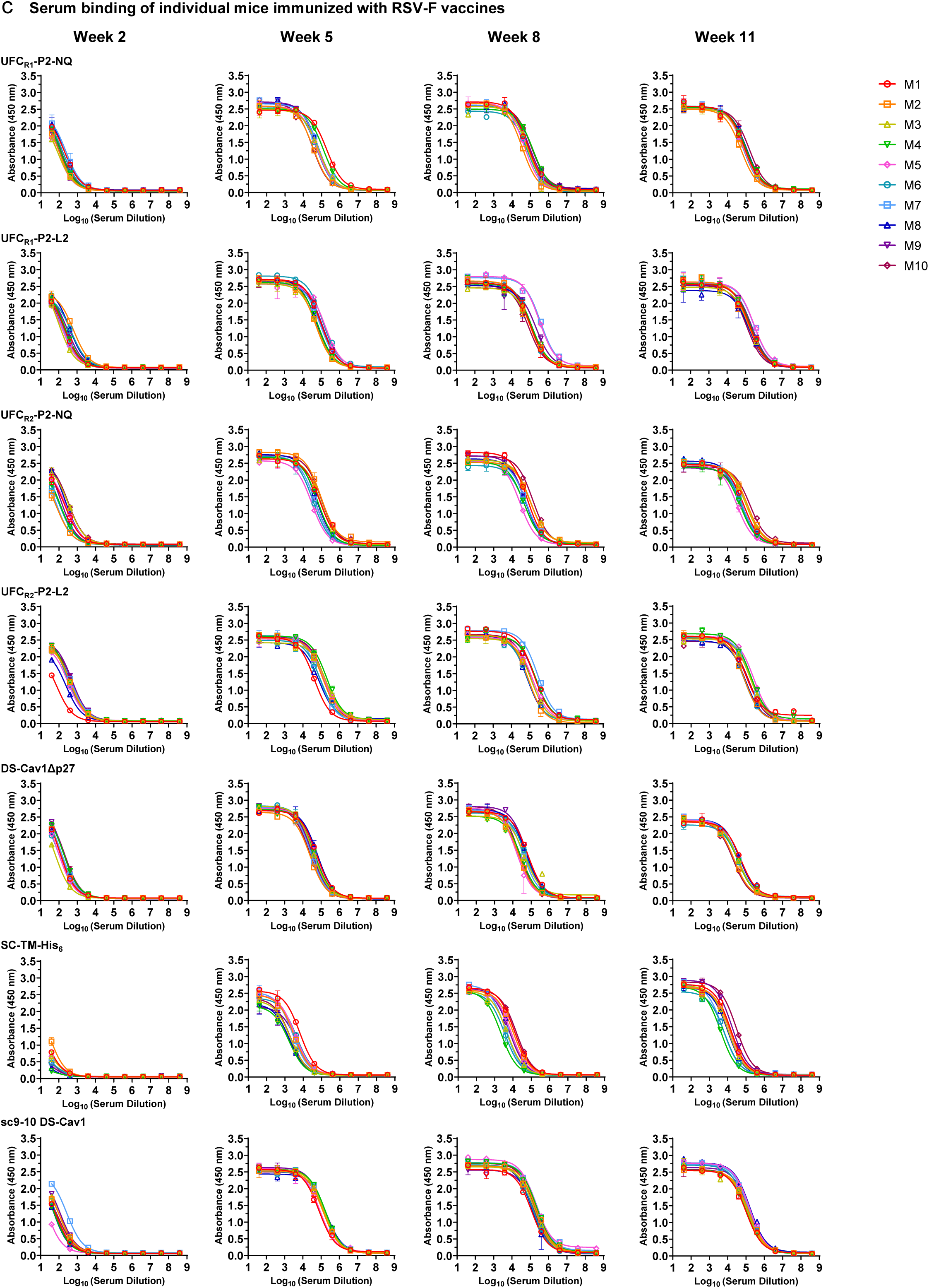
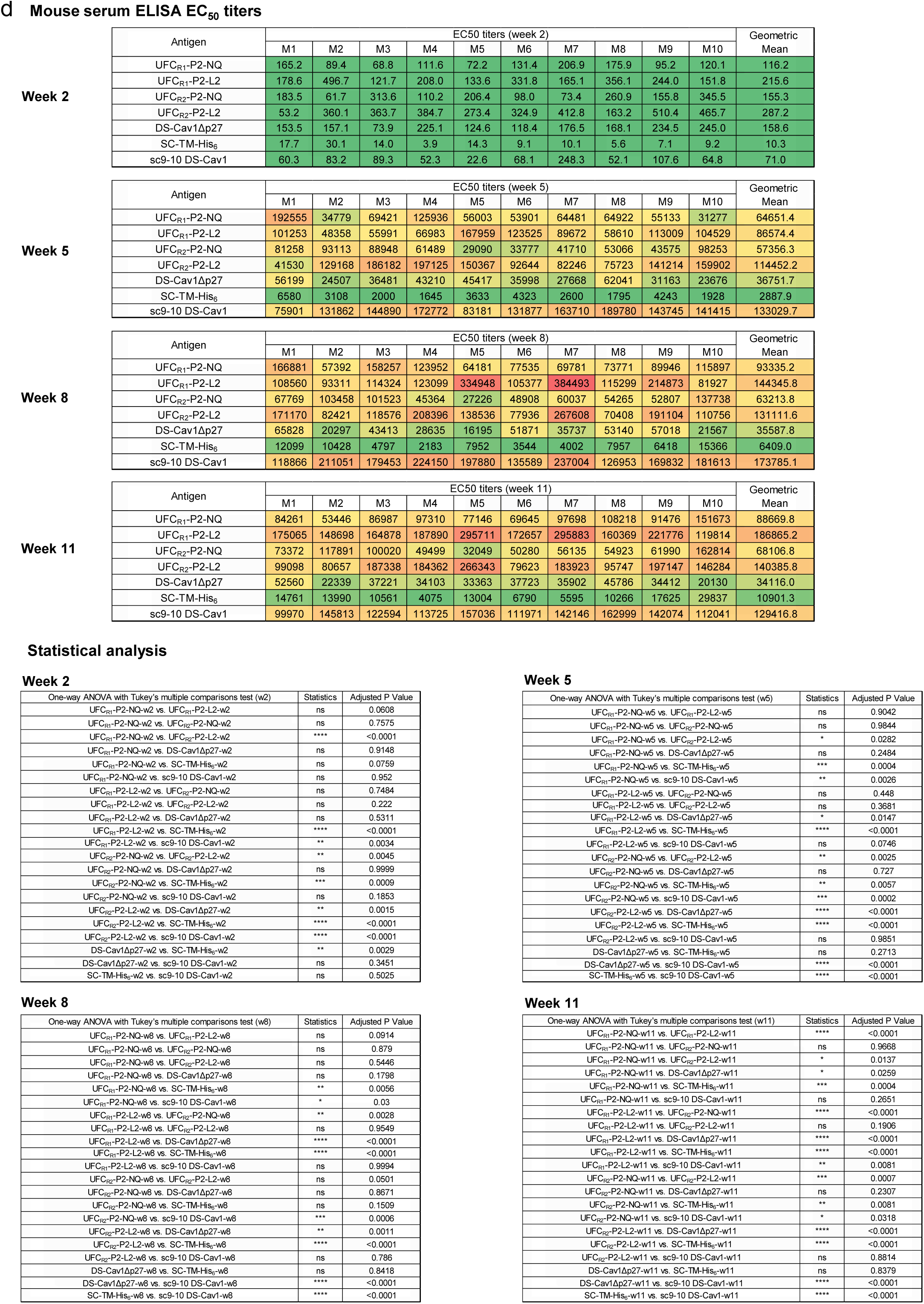
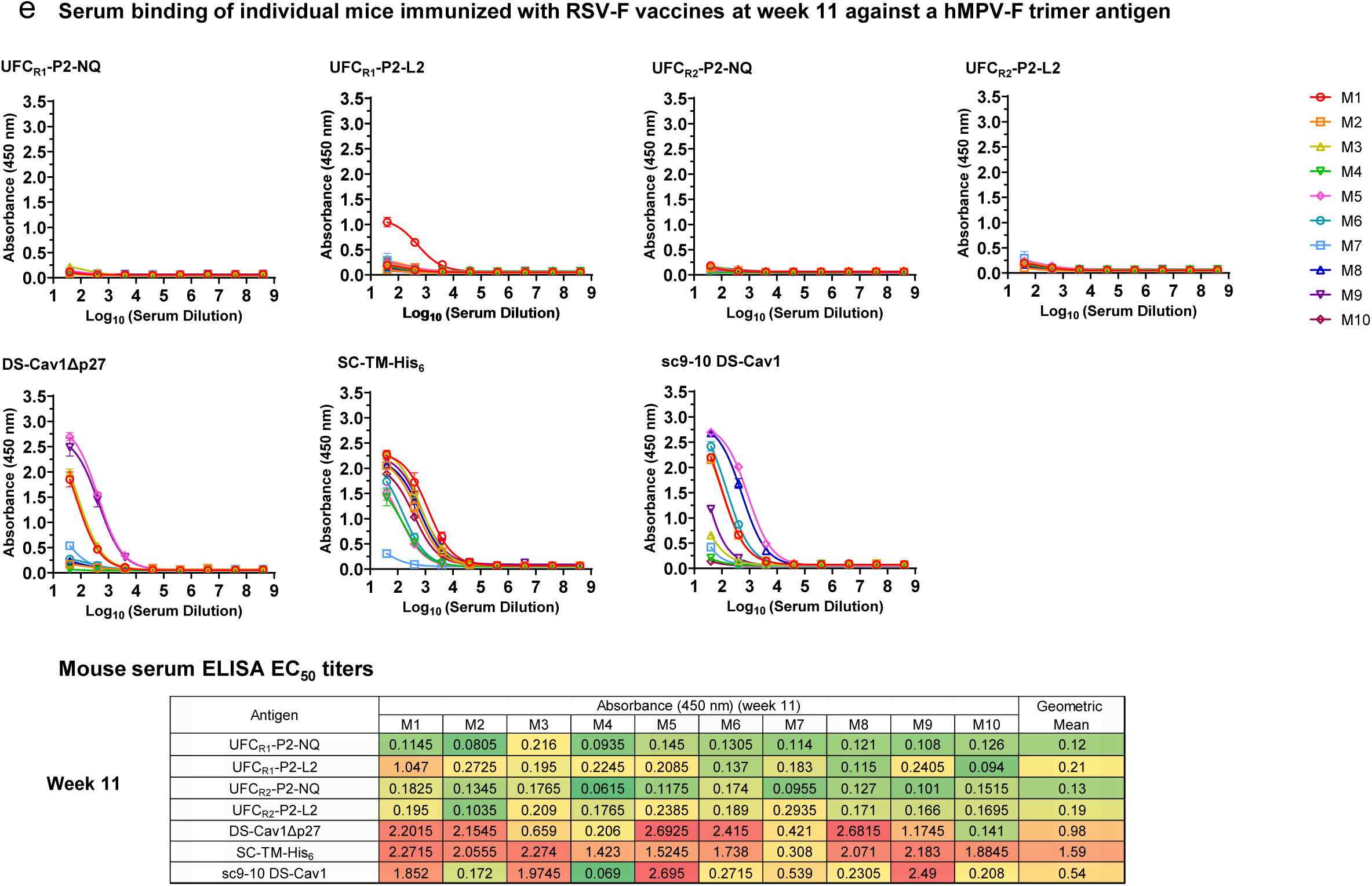
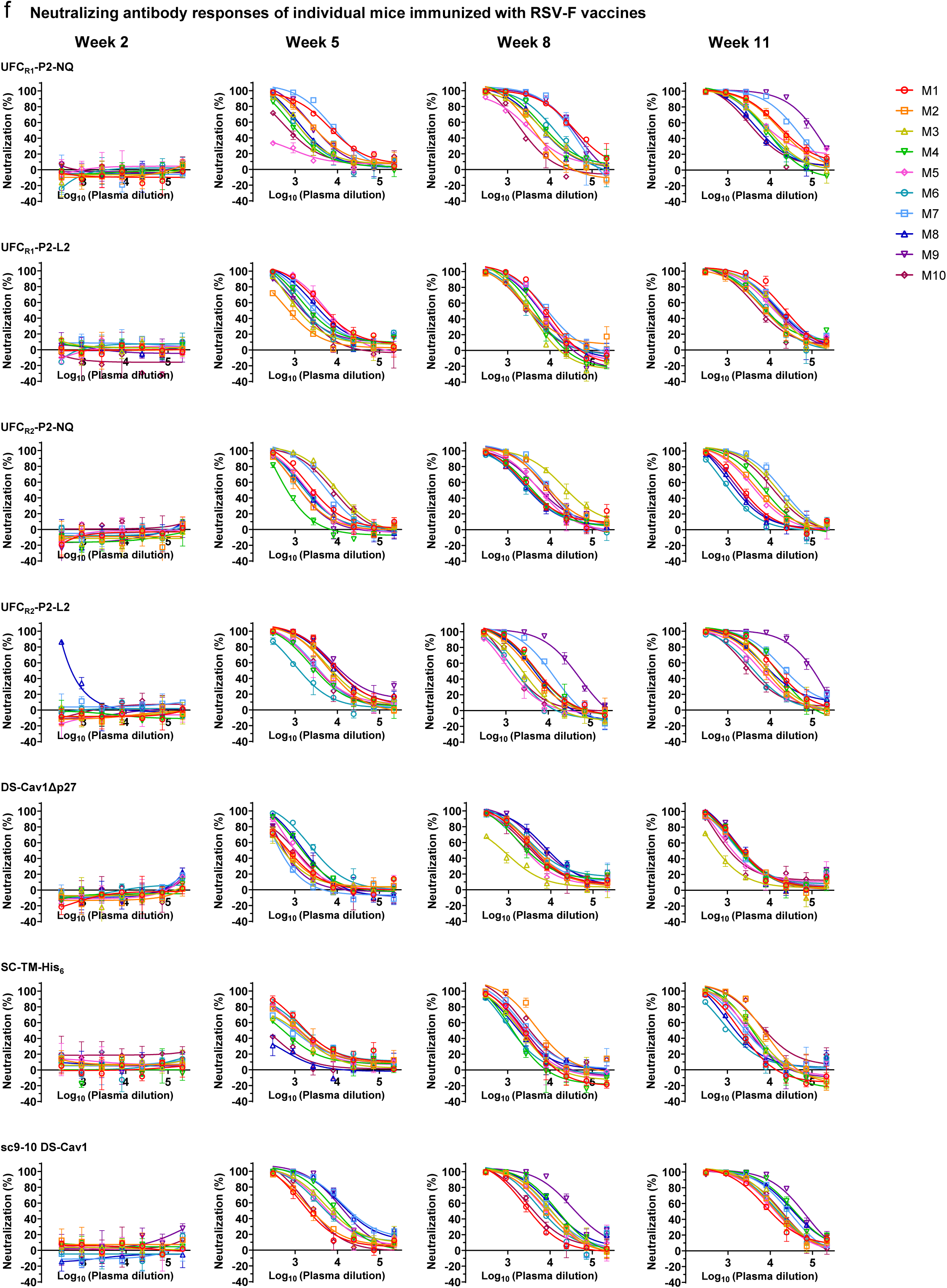
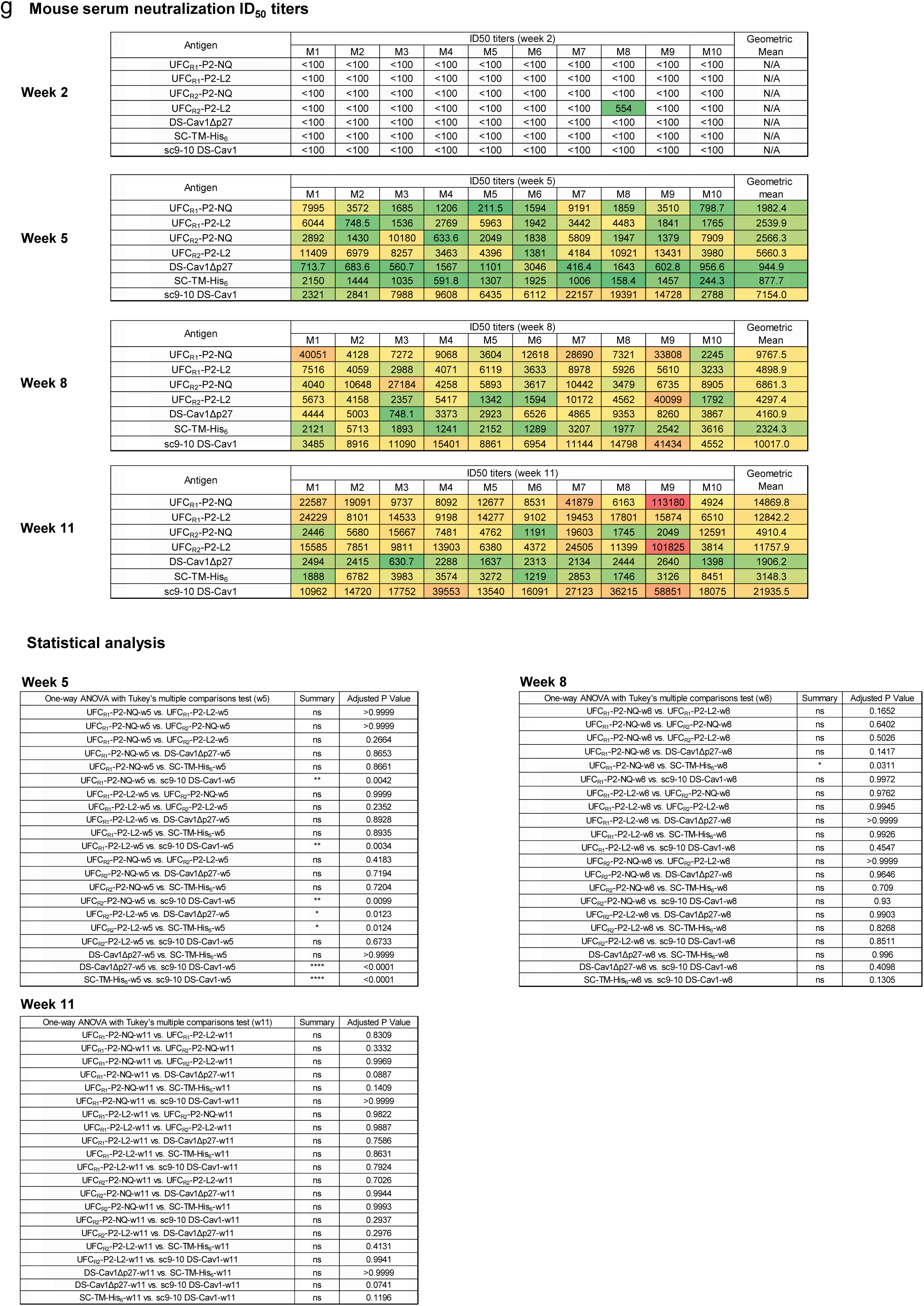
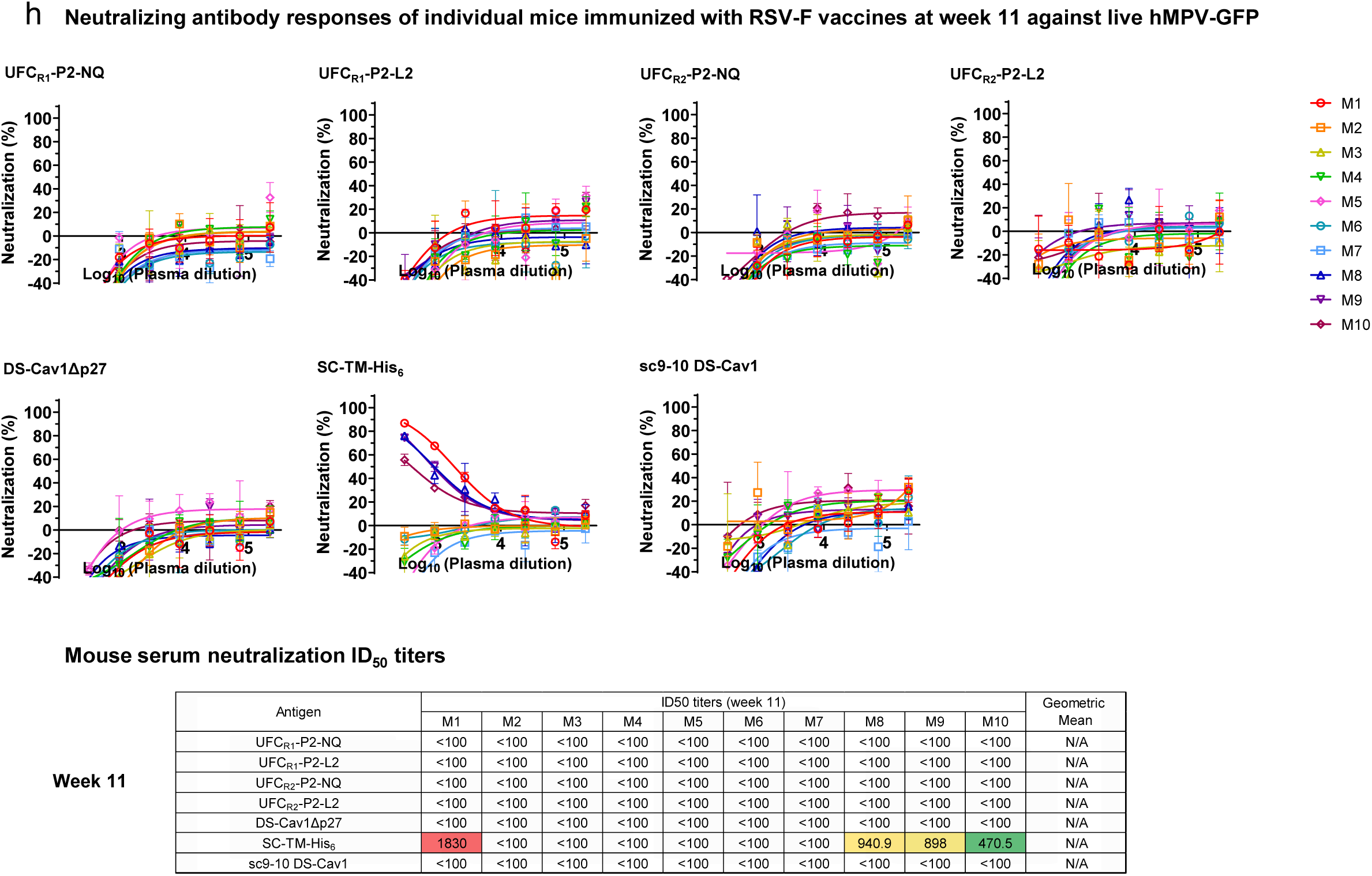
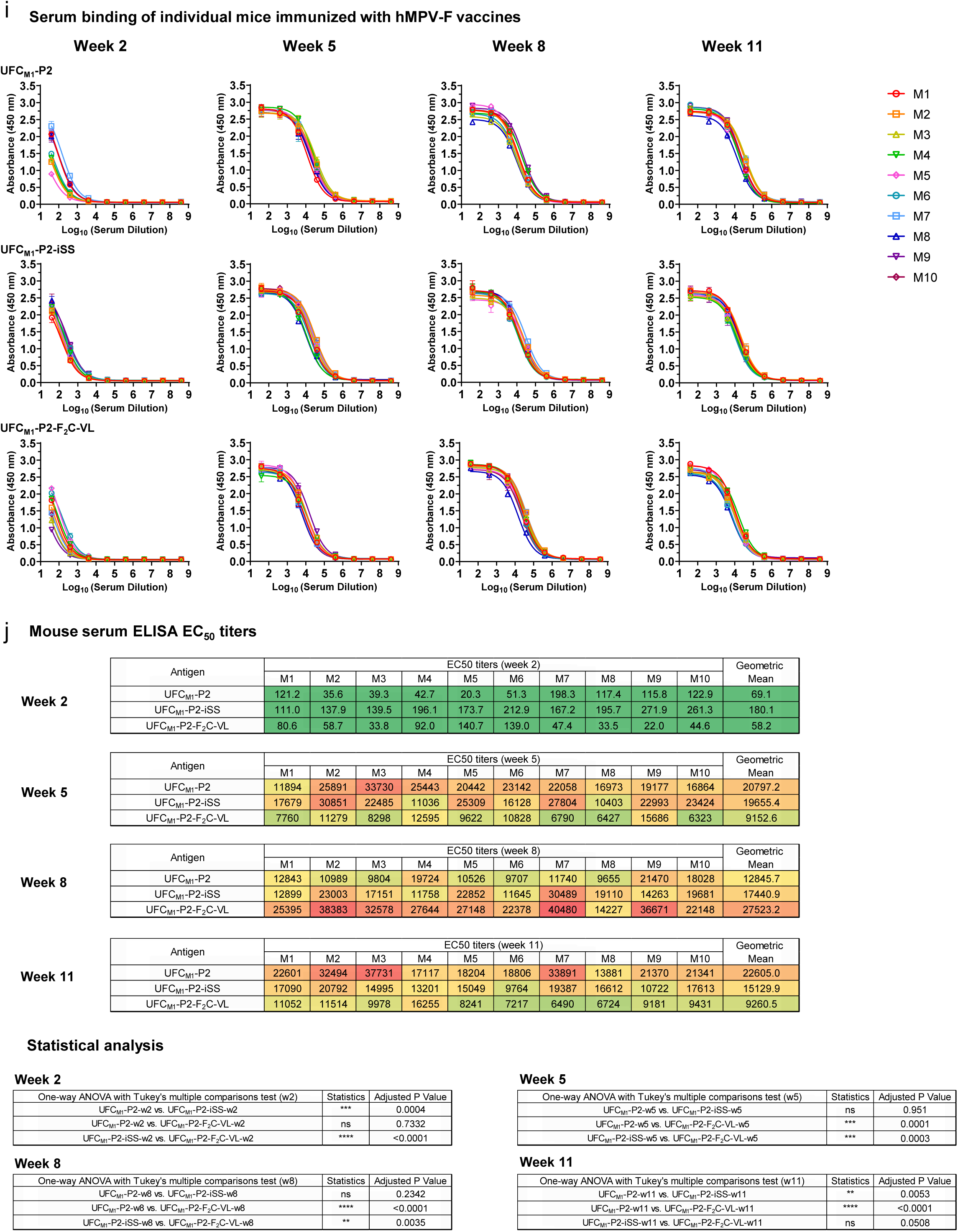
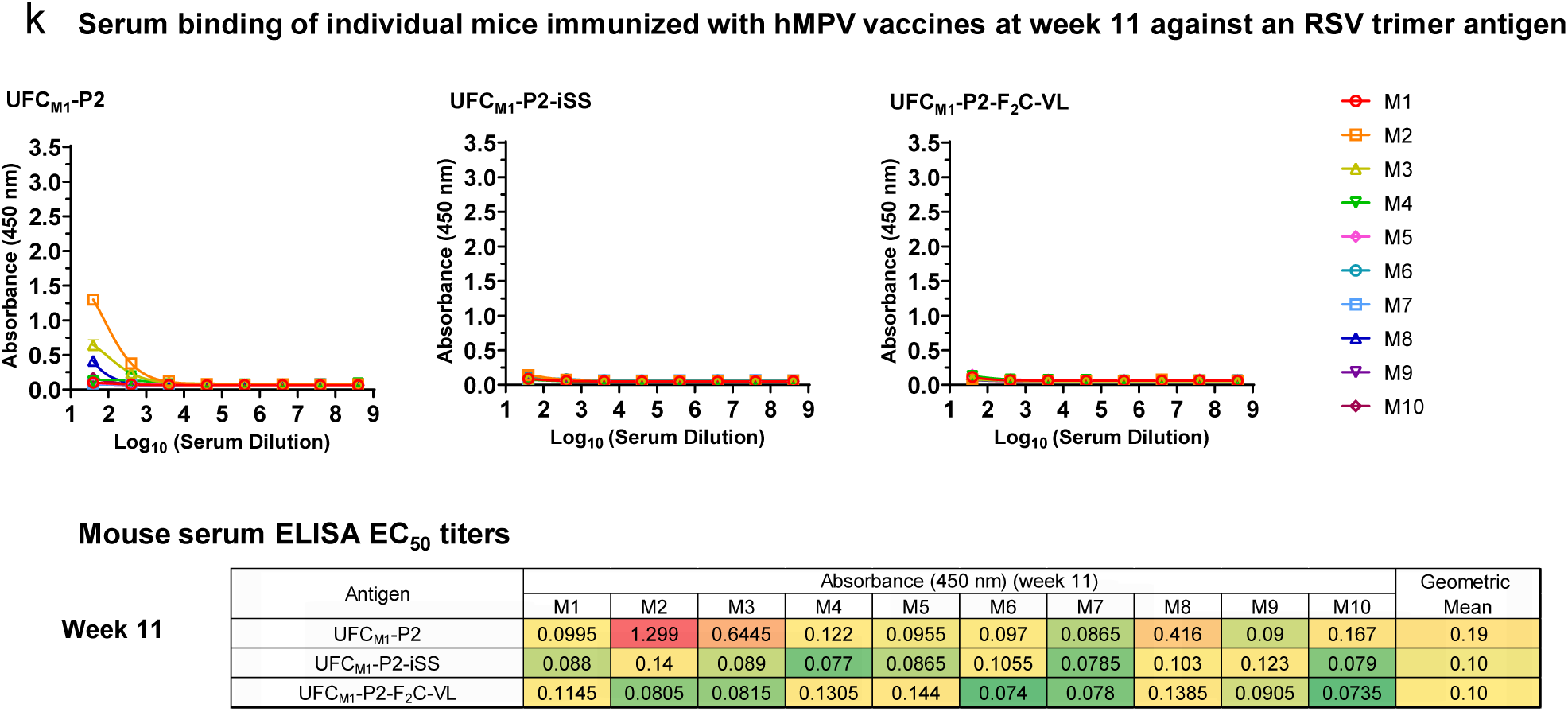
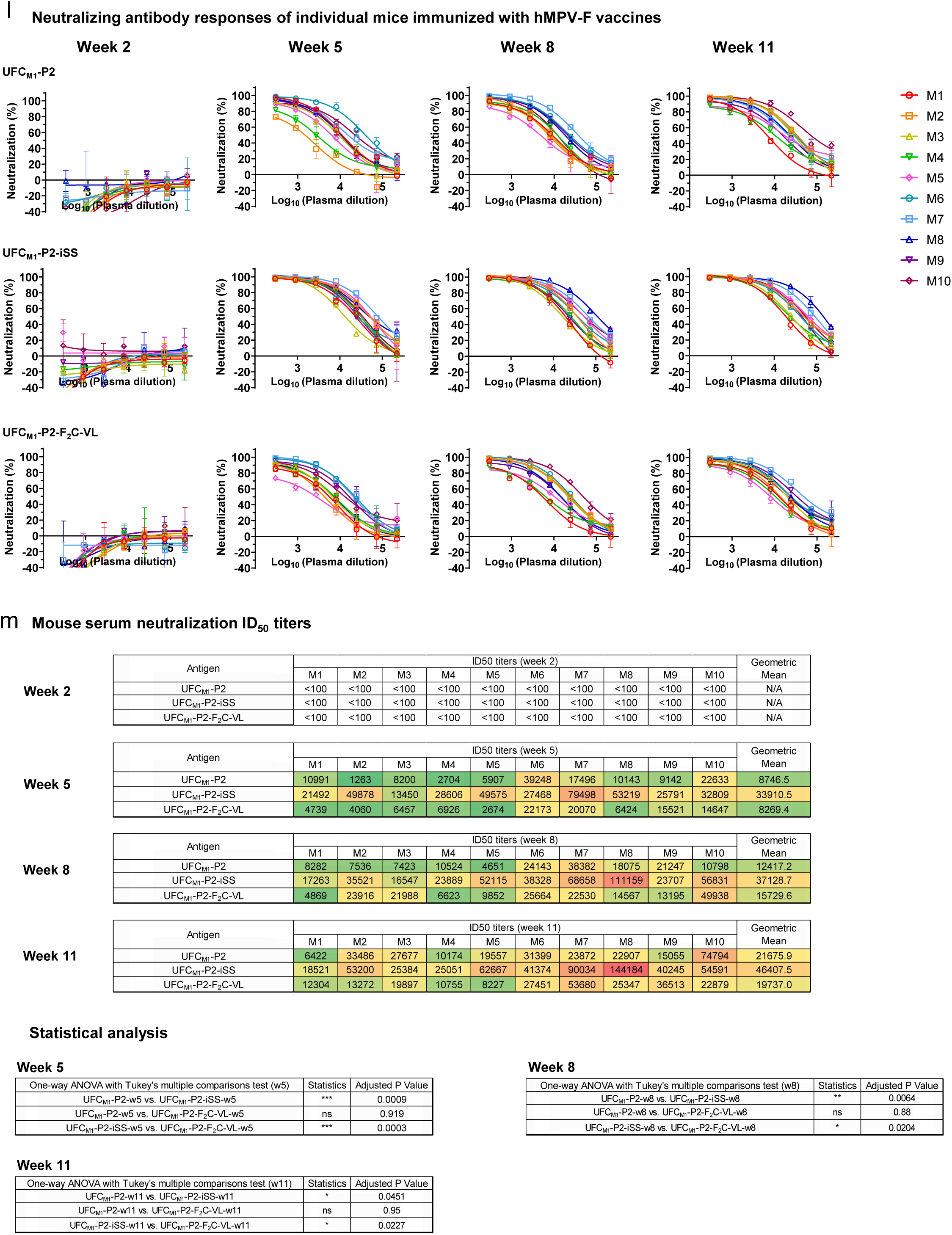
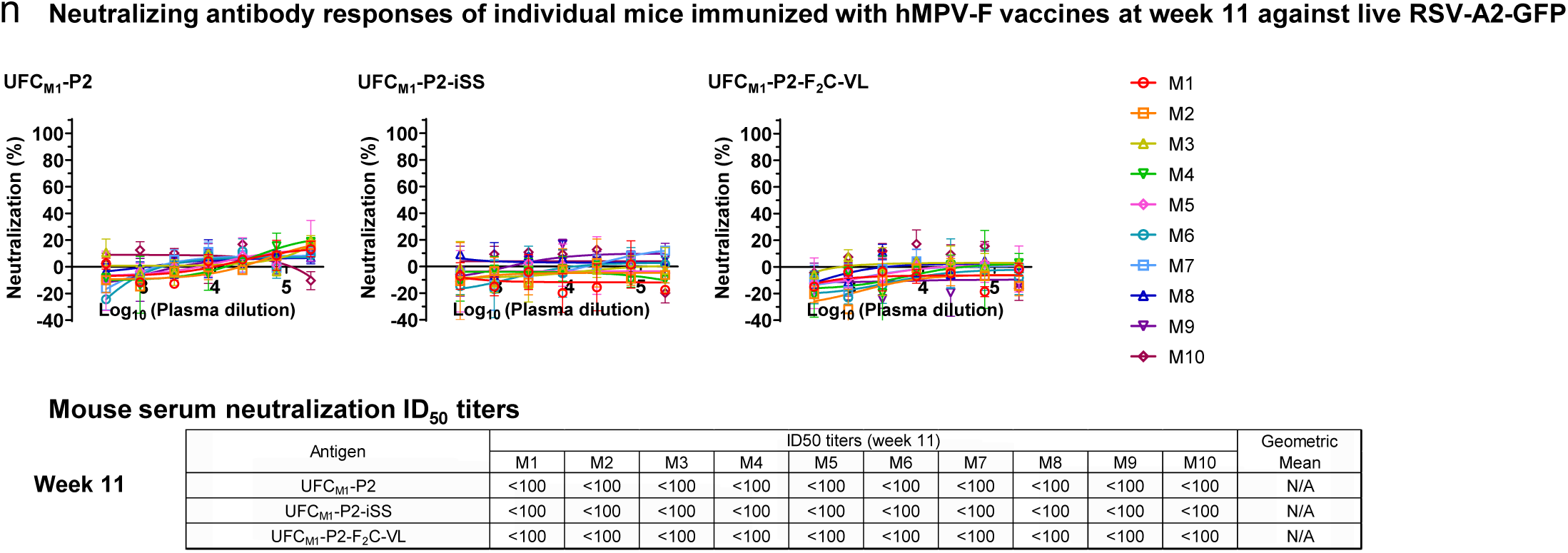
Immunogenicity of RSV-F and hMPV-F vaccines in mice. (**a**) In vitro characterization of DS-Cav1Δp27 following expression in 25 ml ExpiCHO cells and D25 purification, including SEC, DSC, and representative 2D classification images. (**b**) Left: Comparison of ELISA-derived EC_50_ (µg/ml) values of two RSV-F DS-Cav1 trimers binding to 10 antibodies, which are labeled on the plots. Of note, DS-Cav1Δp27 was purified using a D25 antibody column and DS-Cav1-His_6_ was purified by a Nickel column. Right: ELISA binding curves of D25-purified DS-Cav1Δp27 with 10 representative antibodies. (**c**) ELISA binding curves of mouse serum from seven RSV-F trimer vaccine groups to the coating antigen RSV sc9-10 DS-Cav1-1TD0. (**d**) Summary of geometric mean values of EC_50_ titers measured for seven RSV-F trimer vaccine groups against the coating antigen RSV sc9-10 DS-Cav1-1TD0. color coding indicates the level of EC50 titers (green to red: low to high binding). Table of statistical analysis was performed using a one-way ANOVA, followed by post-hoc analysis using Tukey’s multiple comparison test for each timepoint. Of note, the EC_50_ values at week 2 were derived by setting the lower/upper constraints of OD_450_ at 0.0/2.6 to achieve greater accuracy. (**e**) ELISA binding curves of mouse serum from seven RSV-F trimer vaccine groups at week 11 after four immunizations to the coating antigen hMPV UFC_M1_-P2-iSS-1TD0. Summary of absorbance values at 450 nm was included. (**f**) Neutralization curves of mouse serum from seven RSV-F trimer vaccine groups against the live RSV-A2-GFP. (**g**) Summary of geometric mean values of ID_50_ titers measured for seven RSV-F trimer vaccine groups against the RSV-A2-GFP. Color coding indicates the level of ID_50_ titers (white: no neutralization; green to red: low to high neutralization). Of note, the ID_50_ values were derived by setting the lower/upper constraints of % neutralization set at 0.0/100.0. Table of statistical analysis was performed. (**h**) Neutralization curves of mouse serum from seven RSV-F trimer vaccine groups at week 11 against the live hMPV-GFP. Summary of ID_50_ values was included. DS-Cav1Δp27, SC-TM-His_6_, and sc9-10 DS-Cav1 groups are included for comparison. (**i**) ELISA binding curves of mouse serum from three hMPV-F trimer vaccine groups to the coating antigen hMPV UFC_M1_-P2-iSS-1TD0. (**j**) Summary of geometric mean values of EC_50_ titers measured three hMPV-F trimer vaccine groups against the coating antigen hMPV UFC_M1_-P2-iSS-1TD0. Table of statistical analysis was performed. Of note, the EC_50_ values at week 2 were derived by setting the lower/upper constraints of OD_450_ at 0.0/2.7 to achieve greater accuracy. (**k**) ELISA binding curves of mouse serum from three hMPV-F trimer vaccine groups at week 11 to the coating antigen RSV sc9-10 DS-Cav1-1TD0. Summary of absorbance values at 450 nm was included. (**l**) Neutralization curves of mouse serum from three hMPV-F trimer vaccine groups against the live hMPV-GFP. (**m**) summary of geometric mean values of ID_50_ titers measured for three hMPV-F trimer vaccine groups against the hMPV-GFP. Table of statistical analysis was performed. (**n**) Neutralization curves of mouse serum from three hMPV-F trimer vaccine groups at week 11 against the live RSV-A2-GFP. Summary of ID_50_ values was included. EC_50_ and ID_50_ titers were calculated in GraphPad Prism 10.0.2. For significance, ns (not significant), \**p* < 0.05, \*\**p* < 0.01, \*\*\**p* < 0.001, and \*\*\*\**p* < 0.0001.

**Table S1.**
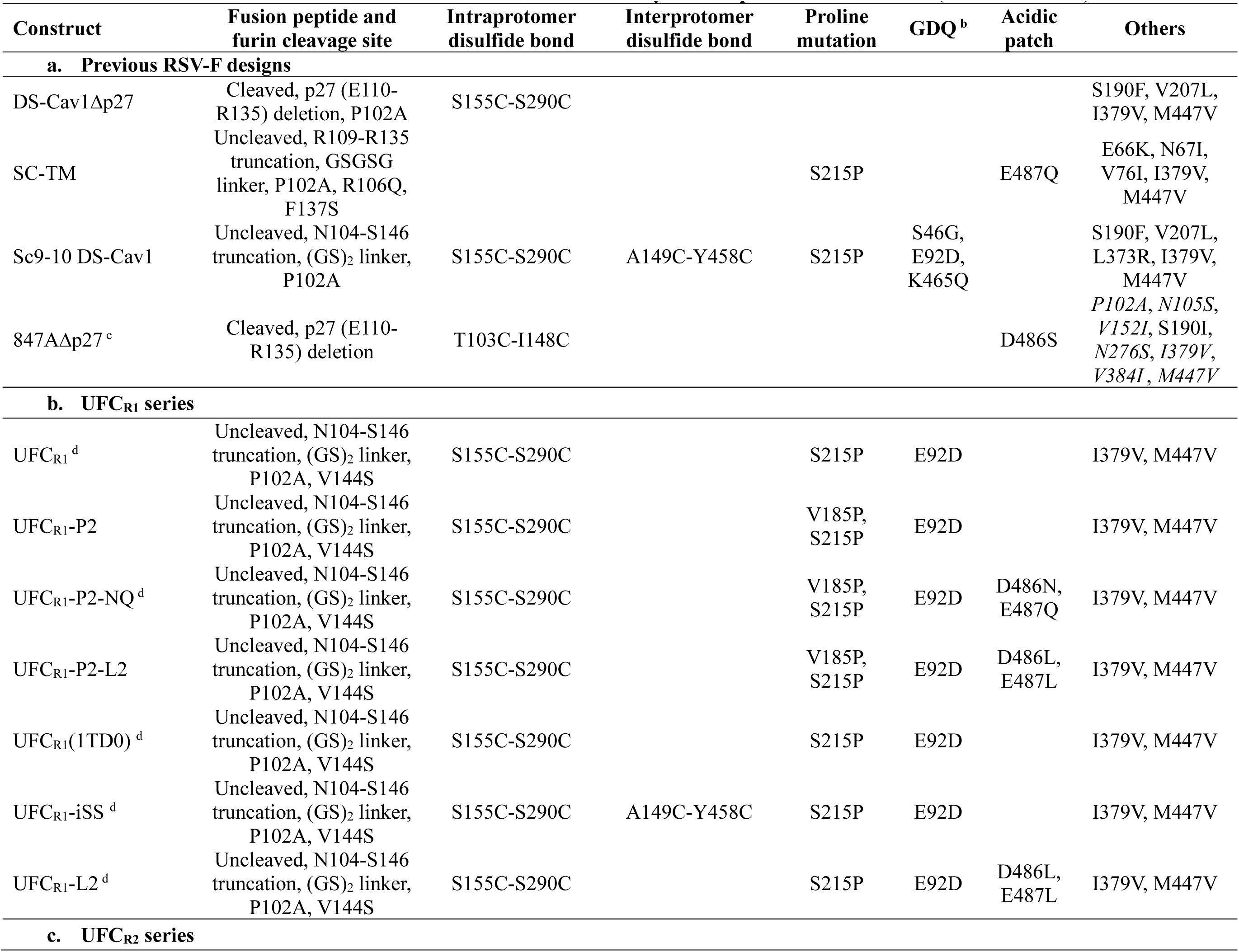
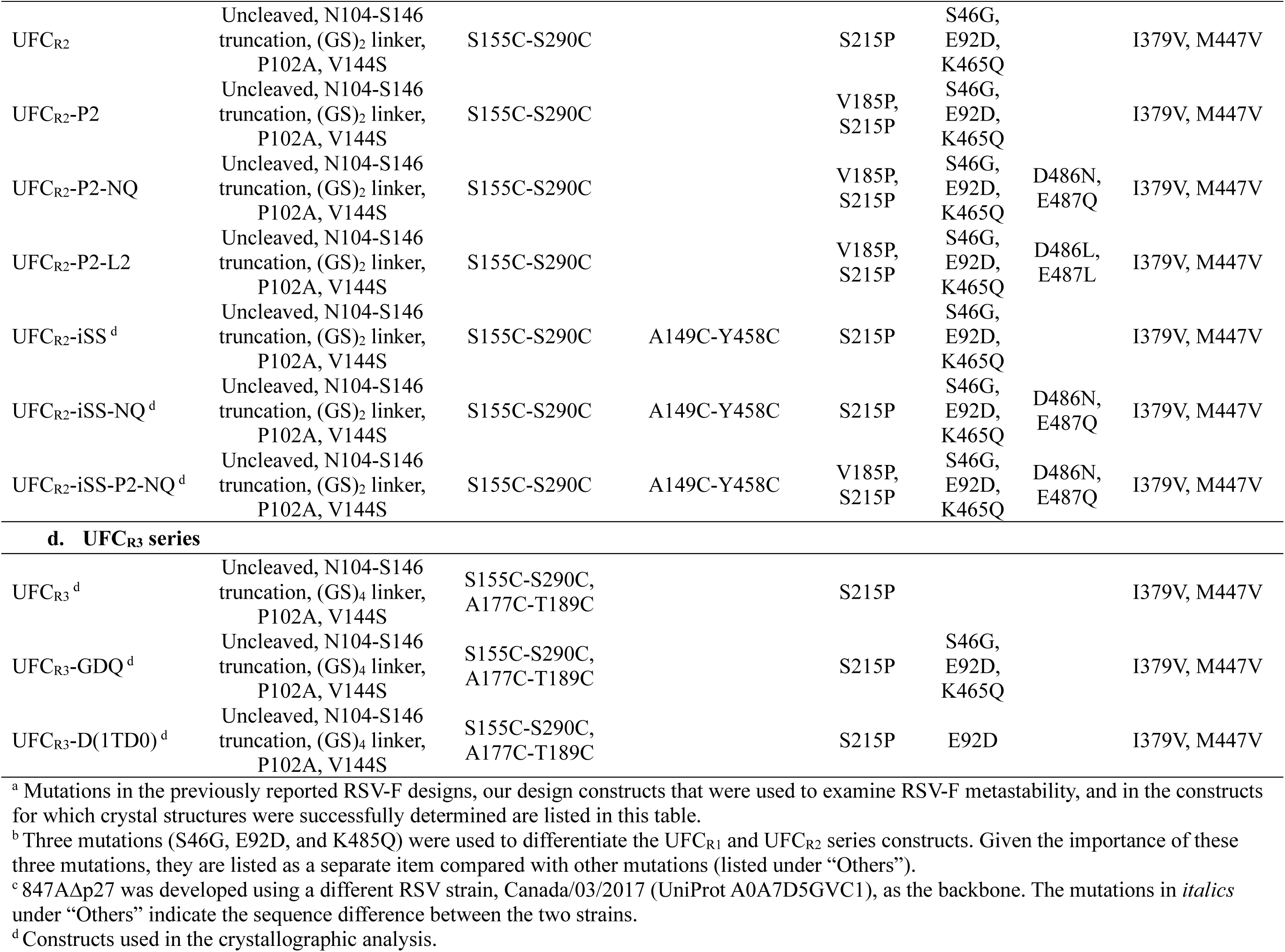
Mutations in RSV-F constructs tested in this study with respect to RSV strain A2 (UniProt P03420) ^a^.

**Table S2.**
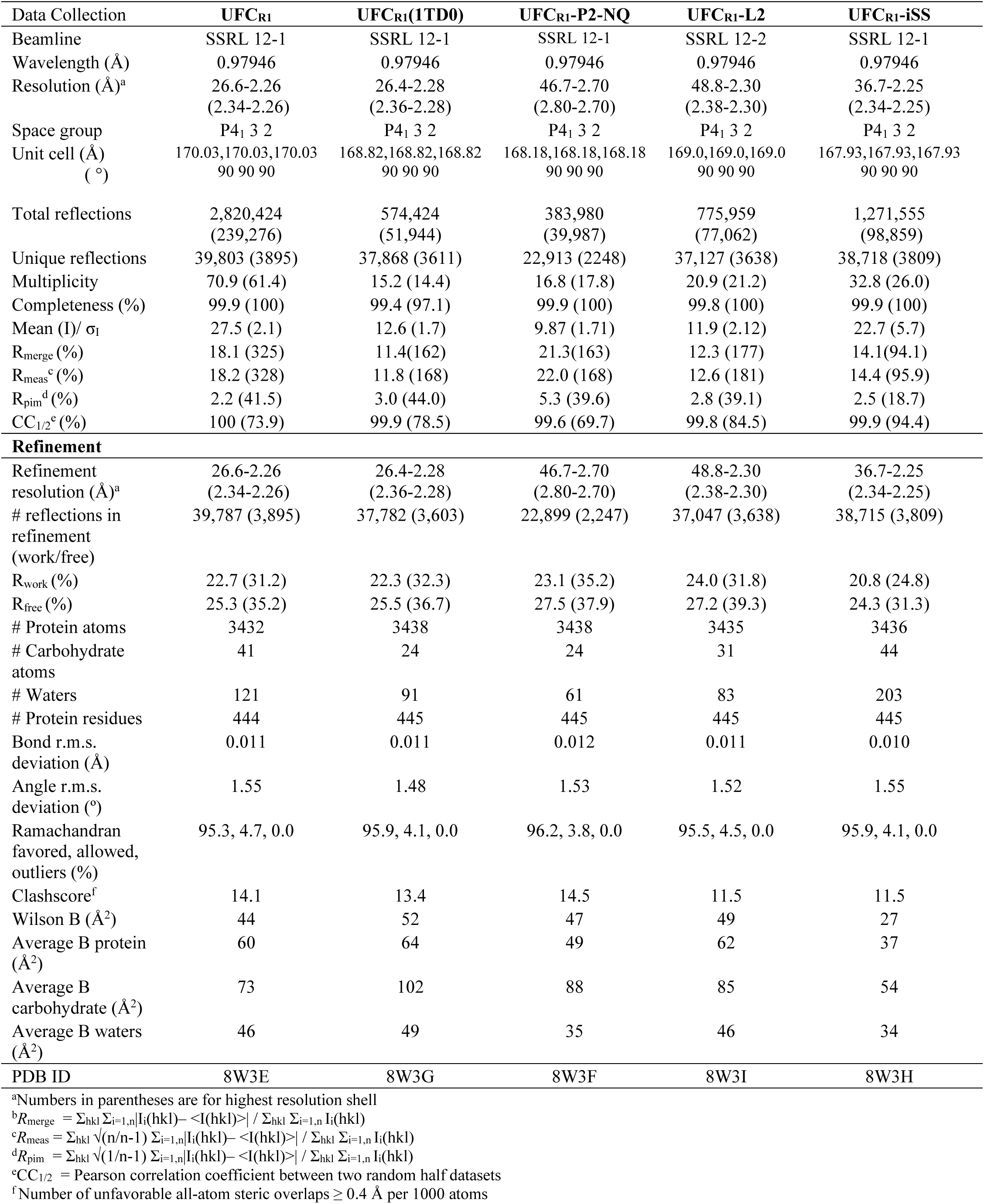
Data collection and refinement statistics for the RSV-F UFC_R1_ series.

**Table S3.**
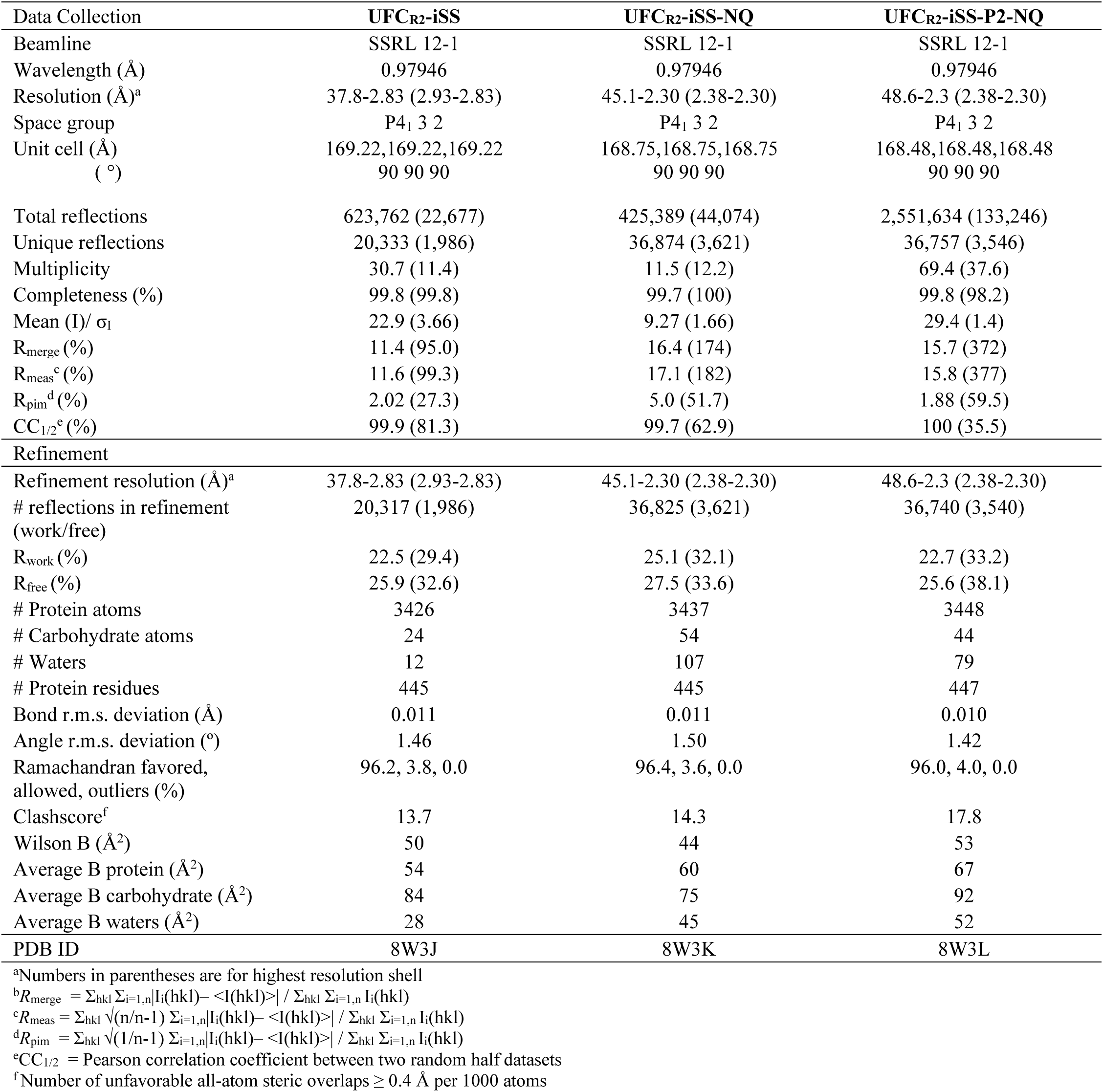
Data collection and refinement statistics for the RSV-F UFC_R2_ series.

**Table S4.**
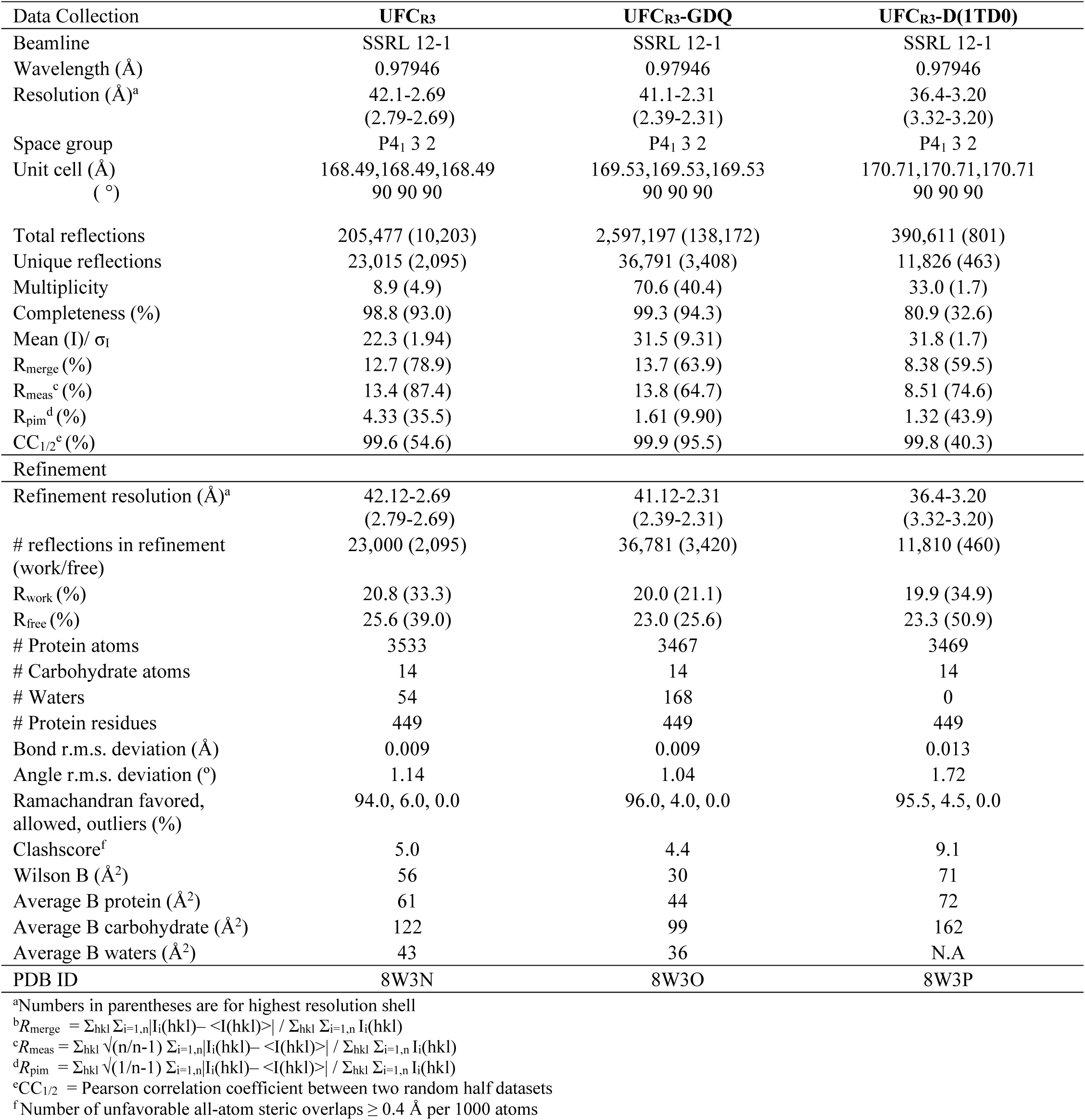
Data collection and refinement statistics for the RSV-F UFC_R3_ series.

**Table S5.**
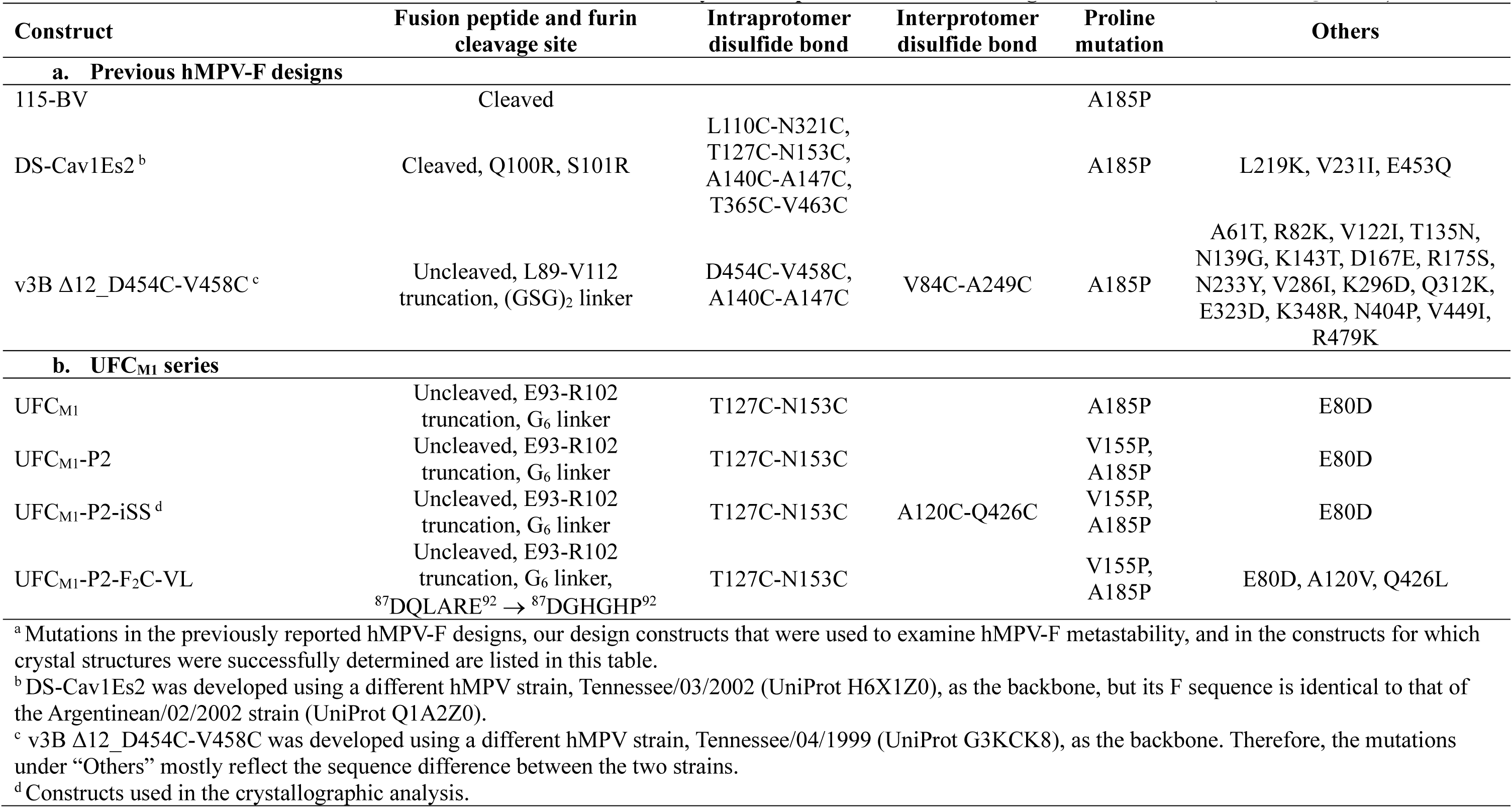
Mutations in hMPV-F constructs tested in this study with respect to hMPV strain Argentinean/02/2002 (UniProt Q1A2Z0) ^a^.

**Table S6.**
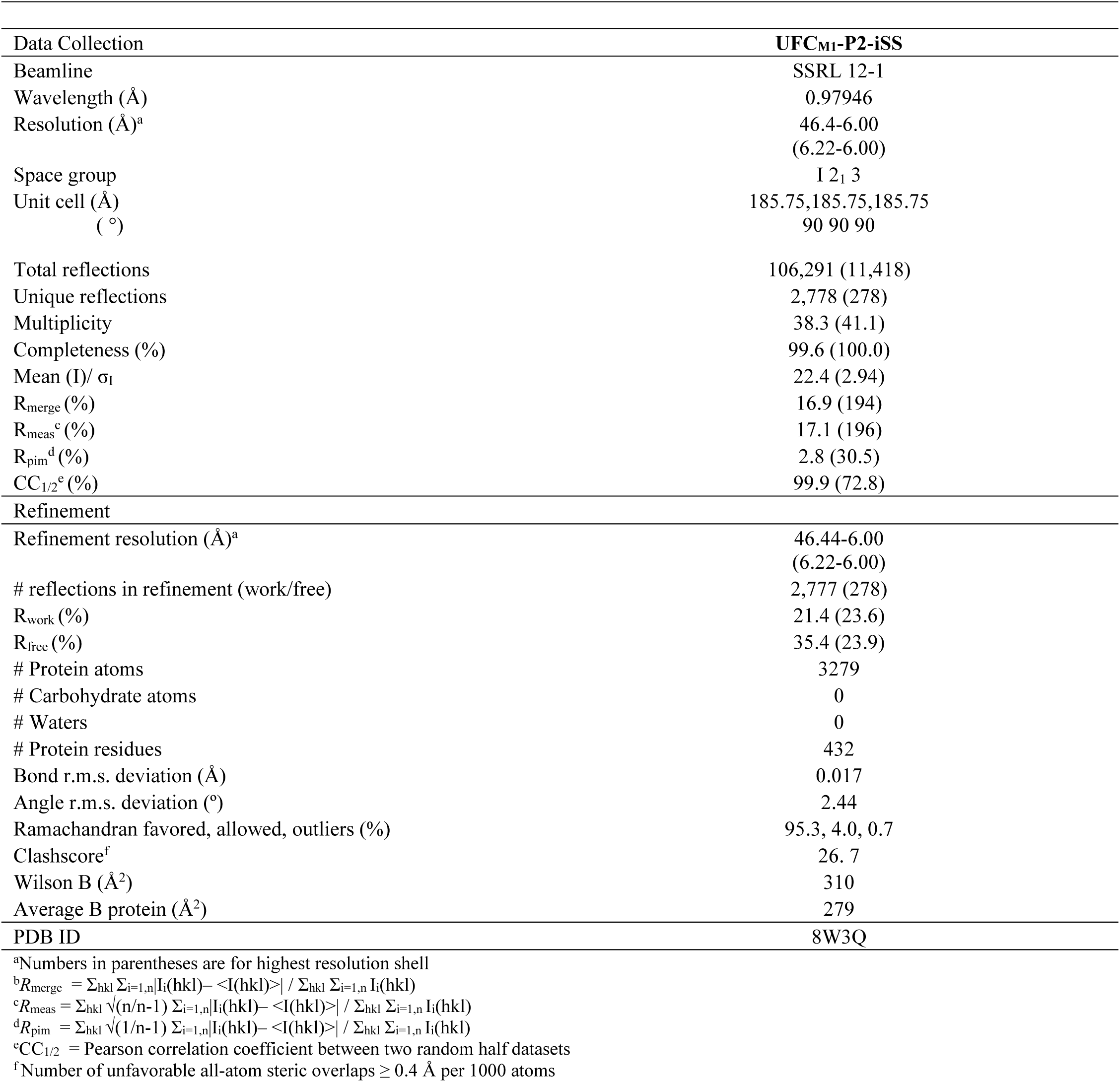
Data collection and refinement statistics for hMPV-F UFC_M1_-P2-iSS.

**Table S7.**
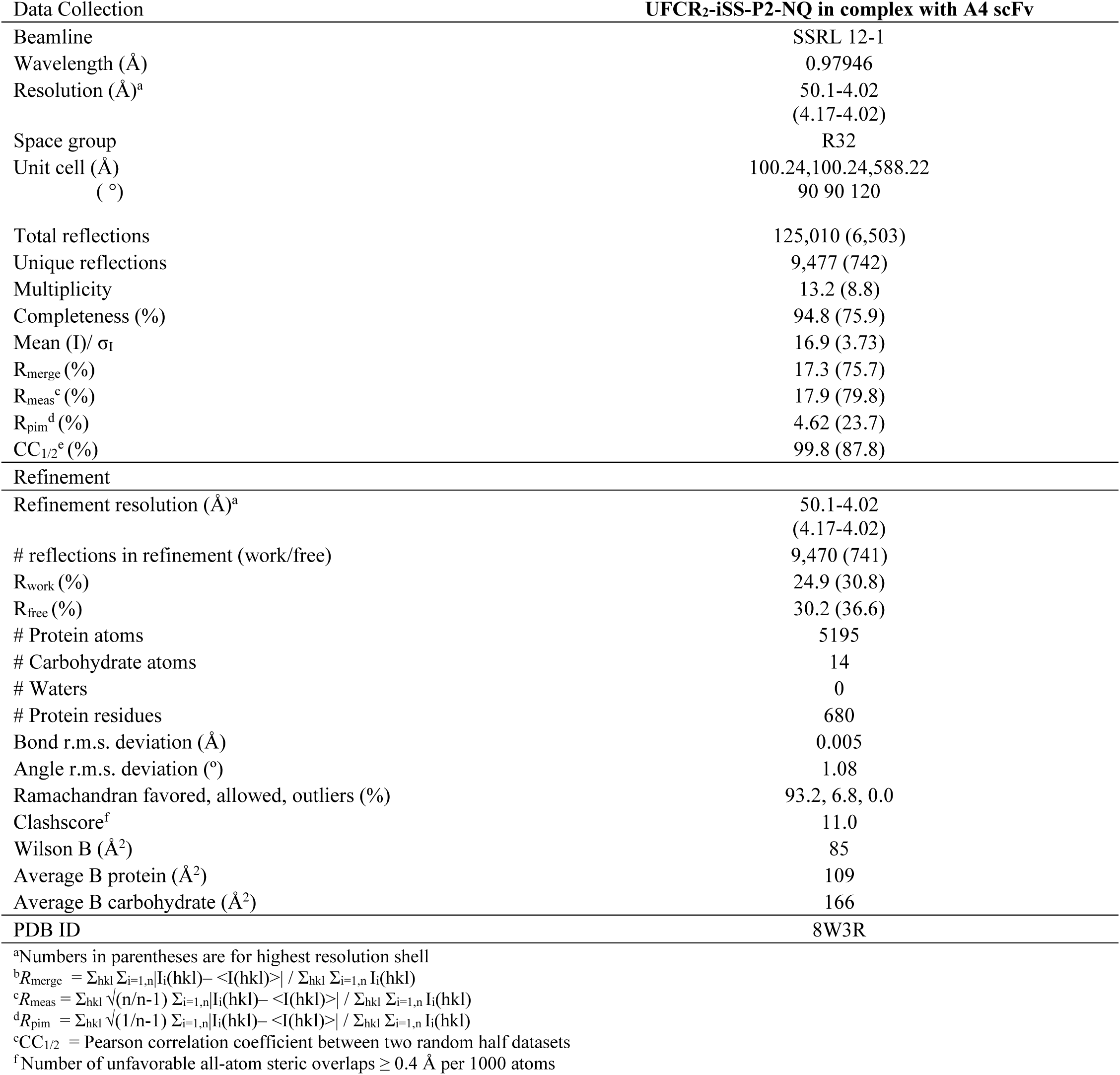
Data collection and refinement statistics for the prefusion RSV-F/A4 antibody complex.

**Table S8.**
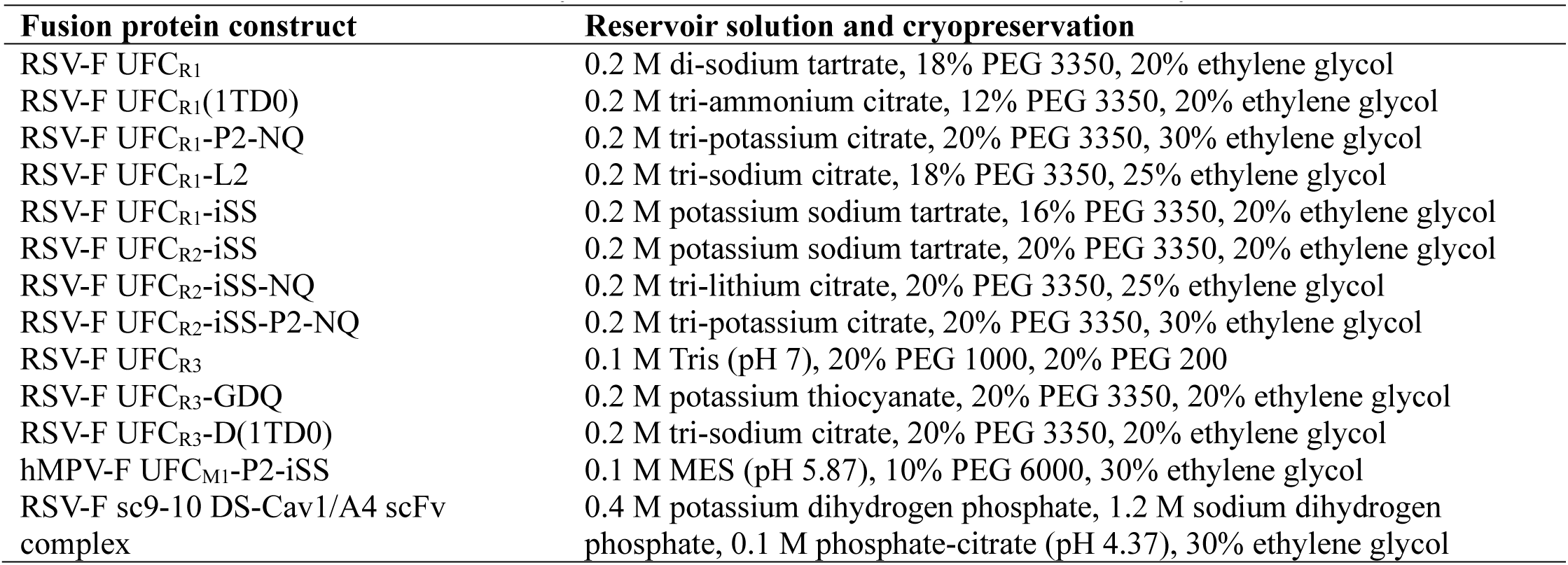
Reservoir solution used for crystallization of various constructs in this study.

## Notes

### Competing Interest Statement

The authors have declared no competing interest.

